# A neural mechanism for learning from delayed postingestive feedback

**DOI:** 10.1101/2023.10.06.561214

**Authors:** Christopher A. Zimmerman, Scott S. Bolkan, Alejandro Pan-Vazquez, Bichan Wu, Emma F. Keppler, Jordan B. Meares-Garcia, Eartha Mae Guthman, Robert N. Fetcho, Brenna McMannon, Junuk Lee, Austin T. Hoag, Laura A. Lynch, Sanjeev R. Janarthanan, Juan F. López Luna, Adrian G. Bondy, Annegret L. Falkner, Samuel S.-H. Wang, Ilana B. Witten

## Abstract

Animals learn the value of foods based on their postingestive effects and thereby develop aversions to foods that are toxic^1–6^ and preferences to those that are nutritious^7–14^. However, it remains unclear how the brain is able to assign credit to flavors experienced during a meal with postingestive feedback signals that can arise after a substantial delay. Here, we reveal an unexpected role for postingestive reactivation of neural flavor representations in this temporal credit assignment process. To begin, we leverage the fact that mice learn to associate novel^15–18^, but not familiar, flavors with delayed gastric malaise signals to investigate how the brain represents flavors that support aversive postingestive learning. Surveying cellular resolution brainwide activation patterns reveals that a network of amygdala regions is unique in being preferentially activated by novel flavors across every stage of the learning process: the initial meal, delayed malaise, and memory retrieval. By combining high-density recordings in the amygdala with optogenetic stimulation of genetically defined hindbrain malaise cells, we find that postingestive malaise signals potently and specifically reactivate amygdalar novel flavor representations from a recent meal. The degree of malaise-driven reactivation of individual neurons predicts strengthening of flavor responses upon memory retrieval, leading to stabilization of the population-level representation of the recently consumed flavor. In contrast, meals without postingestive consequences degrade neural flavor representations as flavors become familiar and safe. Thus, our findings demonstrate that interoceptive reactivation of amygdalar flavor representations provides a neural mechanism to resolve the temporal credit assignment problem inherent to postingestive learning.

## Introduction

Postingestive feedback signals arise from the gut as food is digested and absorbed, and animals are able to associate this delayed feedback with flavors experienced during a meal minutes or hours earlier^1–14^. This learning process is essential for survival — nutritious foods are valuable, while poisonous foods can be deadly — but it remains unknown how the brain is able to associate a stimulus (flavor) with a delayed reinforcement signal (postingestive feedback from the gut) that can arrive much later.

Conditioned taste aversion (CTA) offers a classic example of this credit assignment problem. Humans^3^ and other animals^1,2^ develop CTAs when they experience symptoms of food poisoning (visceral malaise, nausea, diarrhea), which produces a long-lasting aversion to the potentially poisonous food. Remarkably, animals can develop a CTA to novel foods after a single pairing (that is, one-shot learning) even with meal-to-malaise delays of several hours^19–21^.

Previous work on CTA has focused on two primary anatomical pathways. The first begins in the mouth and sends taste signals to the gustatory insular cortex^22–27^, which then transmits this signal to the basolateral amygdala (BLA)^28–34^. The second begins in the gut and sends malaise signals to a genetically defined population of glutamatergic neurons in the hindbrain parabrachial nucleus (CGRP, calcitonin gene-related peptide, neurons)^4,35–37^ that then project to the central amygdala (CEA) and bed nucleus of the stria terminalis (BST). However, it remains unclear where and how temporally separated taste and malaise signals ultimately converge in the brain to support learning.

### Novel but not familiar flavors support one-shot CTA learning

To gain insight into this longstanding question, we leveraged the fact that novel flavors support CTA after a single pairing with malaise (one-shot learning), whereas familiar flavors that are already known to be safe do not^15–18^. Mice consumed a palatable flavor (sweetened grape Kool-Aid) that was either novel (no prior exposure before conditioning) or familiar (four daily pre-exposures before conditioning) followed, after a 30-min delay, by injection of the emetic drug lithium chloride (LiCl) to induce visceral malaise^21^. We then assessed learning two days later with a two-bottle memory retrieval test (Fig. 1a). We used a flavor (combination of taste and odor) in our study for ethological validity — animals rarely encounter a taste alone and use both taste and odor to avoid foods that made them ill^38,39^ — and because even CTA to pure tastants appears to involve odor conditioning^40^. Consistent with previous work^15–18^, mice for which the malaise-paired flavor was novel developed a strong and stable aversion, whereas mice for which the same flavor was familiar continued to prefer it to water even after the pairing with malaise (Fig. 1b).

**Fig. 1.**
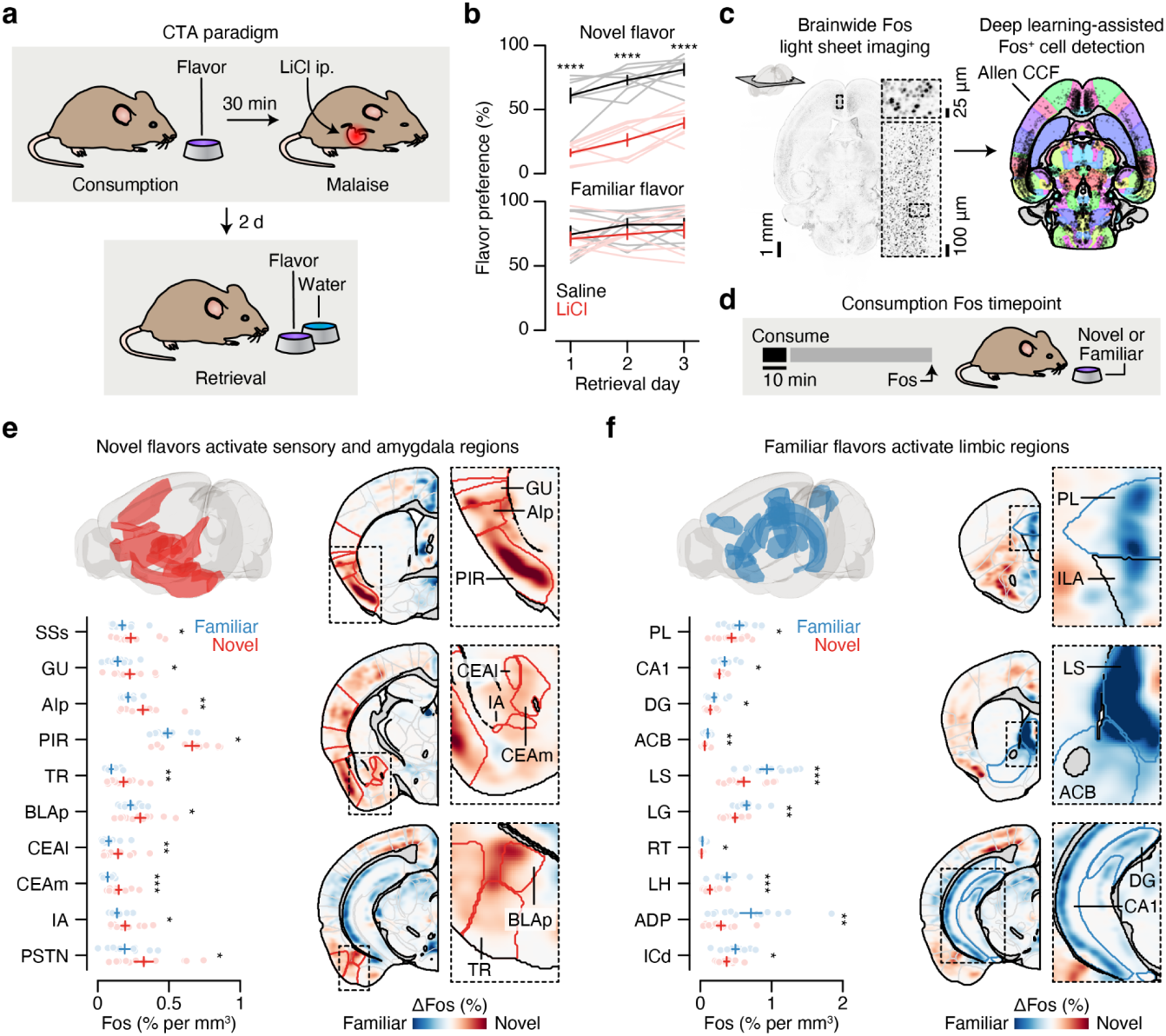
Consumption of a novel flavor supports one-shot CTA learning and activates different brain regions than the same flavor when familiar. **a.** Schematic of the CTA paradigm. **b,** Flavor preference across three consecutive daily retrieval tests for mice that consumed either a novel (top) or familiar (bottom) flavor and then were injected with either LiCl (red) or saline (black) on pairing day (*n* = 8 mice per group). The specific flavor (sweetened grape kool-aid) and amount consumed (1.2 ml) was the same for all groups. The familiar group was pre-exposed to the flavor on four consecutive days before conditioning, whereas the novel group was completely naïve. **c,** Schematic and example Fos expression data (100-µm maximum intensity projection) for the brainwide light sheet imaging pipeline. **d,** Schematic of the Consumption Fos timepoint (*n* = 12 mice per group). The line above “10 min” is a scale bar and the gray bar represents the 60-min wait before perfusion. **e,** Novel flavors preferentially activate sensory and amygdala regions. Left: Comparison of individual familiar (blue) and novel (red) flavor condition mice for every significantly novel flavor-activated brain region. Right: Visualization of the spatially resolved difference in Fos^+^ cell density across flavor conditions with Allen CCF boundaries overlaid. **f,** Familiar flavors preferentially activate limbic regions. Panels are analogous to **e**. *P*-values in **b** are from GLMM marginal effect *z*-tests corrected for multiple comparisons across retrieval days within each flavor condition. *P*-values in **e,f** are from GLMM marginal effect *z*-tests corrected for multiple comparisons across timepoints within each brain region. Error bars represent mean ± s.e.m. \**P* ≤ 0.05, \*\**P* ≤ 0.01, \*\*\**P* ≤ 0.001, \*\*\*\**P* ≤ 0.0001. See Extended Data Table 1 for list of brain region abbreviations.

### Brainwide Fos imaging uncovers brain regions activated by novel or familiar flavors

Using this paradigm, we compared brainwide activation in response to the same flavor when it was novel versus familiar, in order to determine where novel flavors that support learning are represented and where this representation converges with postingestive malaise signals. After each stage of CTA learning (consumption, malaise, retrieval), we cleared animals’ brains using iDISCO+^41,42^, immunolabeled the immediate early gene Fos as a proxy for neural activation, and acquired high-resolution and high-signal-to-noise whole-brain light sheet imaging volumes (∼2-μm isotropic resolution). We employed an automated deep learning-assisted cell detection pipeline (see Methods for details; 258,555 ± 14,421 Fos^+^ neurons/animal across all experiments; mean ± s.e.m.) and registered the location of each Fos^+^ neuron to the Allen Common Coordinate Framework (CCF)^43^ for downstream analysis (Fig. 1c).

We initially investigated the first stage of the CTA paradigm: consumption of flavored water (Fig. 1d). We observed striking differences in the brainwide activation patterns of mice that consumed novel versus familiar flavors, despite the fact that each group had precisely the same sensory experience during Fos induction (Fig. 1e,f; Extended Data Fig. 1a,b; Extended Data Table 1; interactive visualization at https://www.brainsharer.org/ng/?id=872, left column). Novel flavors that support learning preferentially activated a set of sensory and amygdala structures (for example, CEA, BLA, insular cortex, and piriform cortex; Fig. 1e). These observations are consistent with previous anatomically targeted Fos studies^44–46^ and with loss-of-function experiments^24,28,31,32^ that demonstrated a causal role for many of these regions in CTA. In contrast, a familiar flavor that animals had previously learned was safe primarily engaged a network of limbic regions (for example, lateral septum (LS), ventral hippocampus, prefrontal cortex, and nucleus accumbens; Fig. 1f). Unlike the novel flavor-activated regions, most of these regions had not previously been implicated in CTA. The LS showed the strongest familiar flavor-dependent activation, and chemogenetic activation of this region during consumption was sufficient to block CTA learning (Extended Data Fig. 2a,b) and amygdala activation by novel flavors (Extended Data Fig. 2c–g), thus validating the potential of our brainwide imaging to identify new regions that contribute to CTA.

### An amygdala network preferentially responds to novel flavors across every stage of learning

We next investigated how the brainwide activation patterns elicited by the novel versus familiar flavor change during postingestive malaise and, days later, memory retrieval (Fig. 2a; interactive visualization at https://www.brainsharer.org/ng/?id=872, middle and right columns). We reasoned that preferential novel flavor activation at these timepoints, respectively, may reveal where flavor and malaise signals converge to support CTA learning, and where the CTA memory is stored and recalled. To accurately estimate the contribution of flavor novelty and experimental timepoint to neural activation in each brain region, we applied a generalized linear mixed model (GLMM) that accounts for variation associated with different technical batches and with sex (Fig. 2b; Extended Data Table 1; see Methods for full description).

**Fig. 2.**
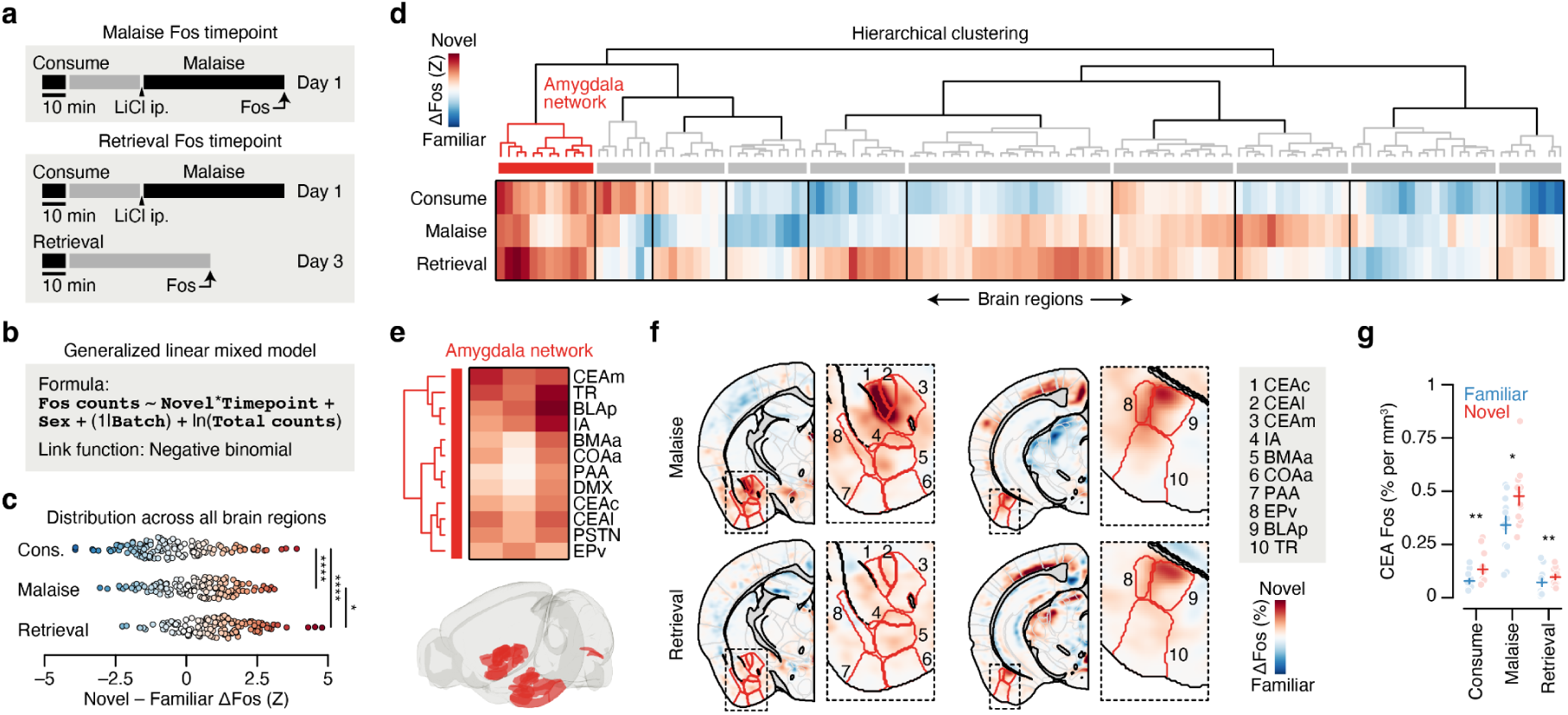
An amygdala network responds to novel flavors across every stage of CTA learning. **a,** Schematic of the Malaise and Retrieval Fos timepoints (*n* = 12 mice per flavor condition per timepoint). There were six total groups of mice in the main Fos dataset: two flavor conditions (Novel, Familiar) × three timepoints (Consumption (from Fig. 1), Malaise, Retrieval). **b,** Description of the GLMM for brainwide Fos data. Briefly, we modeled the number of Fos^+^ neurons in each brain region (**`Fos counts`**) as a fixed effect interaction of flavor condition (**`Novel`**) and experimental timepoint (**`Timepoint`**; in the formula, * represents all possible main effects and interactions) with additional contributions from sex (fixed effect: **`Sex`**), technical batch (random effect: (1|**`Batch`**)), and total brainwide Fos^+^ cell count (offset term: ln(**`Total Counts`**)) using a negative binomial link function. This model properly accounts for the statistical structure of brainwide Fos data as well as batch-to-batch variation in tissue clearing, immunolabeling, and imaging. We then used this model to calculate the average marginal effect (Novel – Familiar ΔFos, in standardized units) of flavor on Fos^+^ cell counts for each brain region (see Methods and Equation 2 for more details). **c,** Novel – Familiar ΔFos effect distribution at each timepoint across brain regions, for all regions that were significantly modulated by **`Novel`**, **`Timepoint`**, or their interaction (*n* = 130 brain regions). Each point represents a single brain region. **d,** Hierarchical clustering of Novel – Familiar ΔFos effects. See Extended Data Fig. 3d for an expanded version. **e,** Detail of the amygdala network (Cluster 1 from **d**) that is preferentially activated by novel flavors at every stage of learning. The heatmap columns in **e** are in the same order as the rows in **d** (left to right: Consumption, Malaise, Retrieval). **f,** Visualization of the spatially resolved difference in Fos^+^ cell density across flavor conditions with Allen CCF boundaries overlaid. **g,** Comparison of individual familiar (blue) and novel (red) flavor condition mice for the CEA at each timepoint. *P*-values in **c** are from Kolmogorov-Smirnov tests corrected for multiple comparisons across timepoints. *P*-values in **g** are from GLMM marginal effect *z*-tests corrected for multiple comparisons across timepoints. Error bars represent mean ± s.e.m. \**P* ≤ 0.05, \*\**P* ≤ 0.01, \*\*\*\**P* ≤ 0.0001. See Extended Data Table 1 for list of brain region abbreviations and for GLMM statistics.

While novel and familiar flavors preferentially activated an equal fraction of brain regions during consumption (Fig. 1e,f), postingestive malaise triggered a brainwide shift towards activation by the novel flavor (Fig. 2c; Extended Data Fig. 3a–c). This shift towards representing the novel flavor was still present when the memory was retrieved days later (Fig. 2c; Extended Data Fig. 3a–c).

To investigate which brain regions contributed to this effect, we performed hierarchical clustering on weights from the GLMM of novel versus familiar flavor activation across experimental timepoints (Fig. 2d; Extended Data Fig. 3d–n). This analysis uncovered a network of amygdala regions (cluster 1) that was activated by novel flavors at every stage of learning (Fig. 2e,f; see Extended Data Fig. 4a–d for correlation analyses within and across clusters), most strikingly in the CEA (Fig. 2g). This discovery is exciting for two reasons. First, the fact that the representation of a novel flavor formed during the initial consumption is still present 30 minutes later during postingestive malaise implies that this network is a site for the convergence of flavor and malaise signals. Second, the observation that this novel flavor representation is still present upon memory retrieval suggests that the same network may also be a site of storage and recall.

### CGRP neurons mediate the effects of postingestive malaise on delayed CTA learning and novel flavor-dependent amygdala activation

The idea that the amygdala could be a critical node for convergence of the flavor representation and malaise signal is further supported by prior work establishing the amygdala as a site of associative learning^47–57^, as well as by the discovery of brainstem neurons that convey visceral malaise to the amygdala: parabrachial CGRP neurons^4,37,58^ (Fig. 3a). Indeed, we found that LiCl injection activated CGRP neurons in vivo^59^ (Fig. 3b), that CGRP neurons formed dense monosynaptic connections in the CEA^59–61^ (Fig. 3c; latency: 5.9 ± 0.3 ms; mean ± s.e.m.), and that CGRP neuron stimulation activated neurons throughout the amygdala network in vivo (Extended Data Fig. 5a–f).

**Fig. 3.**
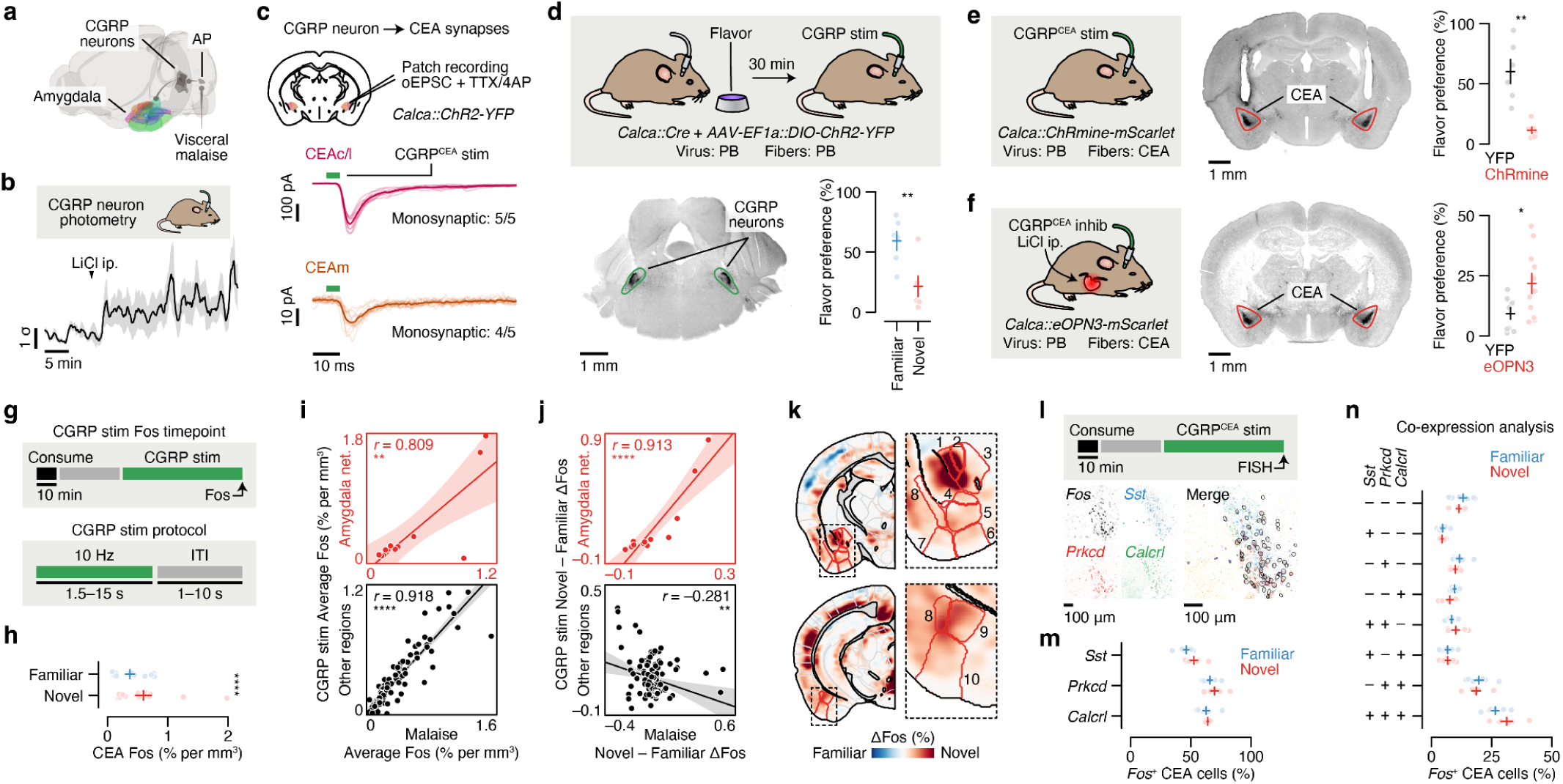
Parabrachial CGRP neurons mediate the effects of postingestive malaise on the amygdala network, and monosynaptic connections to the CEA support the acquisition of delayed CTA. **a,** Schematic of the neural pathway that conveys visceral malaise signals from the gut to the amygdala via the area postrema (AP) and CGRP neurons in the parabrachial nucleus (PB)^4–6,37,58,59^. **b,** Fiber photometry recordings showing that CGRP neurons are activated in vivo by LiCl-induced malaise (*n* = 5 mice). **c,** Top: Strategy for using ChR2-assisted circuit mapping^118^ to identify monosynaptic connections (optogenetically evoked excitatory postsynaptic currents (oEPSCs) in the presence of the voltage-gated sodium channel blocker tetrodotoxin (TTX) and the voltage-gated potassium channel blocker 4-aminopyridine (4AP)) between CGRP neurons and the CEA. Middle: Strong monosynaptic inputs from CGRP neurons to the CEAc/l (amplitude: –327.0 ± 136.3 pA; mean ± s.e.m.; *n* = 5/5 neurons from 3 mice). Bottom: Weaker monosynaptic inputs from CGRP neurons to the CEAm (amplitude: –15.6 ± 6.4 pA; mean ± s.e.m.; *n* = 4/5 neurons from 3 mice). The dark lines represent the average and the transparent lines represent individual trials for each example neuron. **d,** Top: Schematic for the CGRP neuron stimulation experiment. Cre-dependent ChR2-YFP virus was injected bilaterally into the PB of *Calca::Cre* mice and optical fibers were implanted bilaterally above the PB. Bottom left: Example ChR2-YFP expression in PB. Bottom right: Retrieval test flavor preference for the same experiment (*n* = 6 mice per group). **e**, Left: Schematic for the CGRP^CEA^ projection stimulation experiment. Cre-dependent ChRmine-mScarlet virus was injected bilaterally into the PB of *Calca::Cre* mice and optical fibers were implanted bilaterally above the CEA. During the behavioral experiment, CGRP^CEA^ projection stimulation began 30-min after novel flavor consumption. Middle: Example ChRmine-mScarlet expression in CEA. Right: Retrieval test flavor preference for the same experiment (*n* = 6 mice per group). **f**, Left: Schematic for the CGRP^CEA^ projection inhibition experiment. Cre-dependent eOPN3-mScarlet virus was injected bilaterally into the PB of *Calca::Cre* mice and optical fibers were implanted bilaterally above the CEA. During the behavioral experiment, CGRP^CEA^ projection inhibition and LiCl injection began 30-min after novel flavor consumption. Middle: Example eOPN3-mScarlet expression in CEA. Right: Retrieval test flavor preference for the same experiment (*n* = 11 eOPN3 mice, 9 YFP mice). **g,** Schematic of the CGRP stim Fos timepoint, which used *Calca::ChR2* mice with optical fibers implanted bilaterally above the PB (*n* = 14 novel flavor mice, 13 familiar flavor mice). **h,** Comparison of CEA Fos in individual familiar (blue) and novel (red) flavor condition mice. **i,** Correlation between the average Fos^+^ cell count across both flavor conditions in each brain region for the LiCl-induced malaise timepoint versus the CGRP stim timepoint for the amygdala network (Cluster 1 from Fig. 2d,e; top; *n* = 12 regions) and for all other regions (bottom; *n* = 117 regions). See also Extended Data Fig. 6c. **j,** Panels are analogous to **i**, but comparing the difference between Novel and Familiar flavor groups. See also Extended Data Fig. 6d. **k,** Visualization of the spatially resolved difference in Fos^+^ cell density across flavor conditions with Allen CCF boundaries overlaid. See Fig. 2f for brain region number legend. **l**, Top: Schematic of the CGRP^CEA^ projection stimulation RNAscope FISH experiment, which used *Calca::ChR2* mice with optical fibers implanted bilaterally above the CEA (*n* = 6 novel flavor mice, 7 familiar flavor mice; 490 ± 54 *Fos*^+^neurons per mouse). Bottom: Example FISH data showing *Fos* expression (as a marker of neural activation) and *Sst*, *Prkcd*, and *Calcrl* expression (as markers of known CEA cell types) in the CEA following CGRP^CEA^ projection stimulation. See Extended Data Fig. 6g,h for an expanded version. **m**, Comparison of marker gene expression in *Fos*^+^ neurons for individual familiar (blue) and novel (red) flavor condition mice. **n**, Comparison of marker gene co-expression. *P*-values in **d**–**f** are from Wilcoxon rank-sum tests. *P*-value in **h** is from a GLMM marginal effect *z*-test. *P*-values in **i**,**j** are from Pearson correlation *t*-tests. All within-gene Novel vs. Familiar comparisons in **m**,**n** are not significant with Wilcoxon rank-sum tests. Error bars represent mean ± s.e.m. Shaded areas in **b** represent mean ± s.e.m. and in **i,j** represent 95% confidence interval for linear fit. Units in **j** are % per mm^3^. \**P* ≤ 0.05, \*\**P* ≤ 0.01, \*\*\*\**P* ≤ 0.0001.

Moreover, CGRP neuron stimulation recapitulated the effects of LiCl-induced malaise in mediating novel flavor-dependent delayed CTA. Specifically, CGRP neuron cell-body stimulation that began 30 min after flavor consumption was sufficient to replace LiCl injection and condition an aversion to a novel but not familiar flavor (Fig. 3d). Similarly, stimulation of CGRP neuron → CEA (CGRP^CEA^) axon terminals 30-min after novel flavor consumption was also sufficient to condition a strong CTA (Fig. 3e), whereas CGRP^CEA^ inhibition^62^ during delayed LiCl-induced malaise significantly interfered with CTA acquisition (Fig. 3f).

Given these similarities between LiCl-induced malaise and CGRP neuron stimulation, we sought to determine if postingestive stimulation of these cells could recapitulate the novel flavor-dependent effects of LiCl-induced malaise on neural activation in the amygdala and across the brain (Fig. 3g). Indeed, postingestive CGRP neuron stimulation produced strikingly similar levels of overall neural activation (Fos^+^ cell counts in individual brain regions) across the entire brain when compared to LiCl-induced malaise (Fig. 3i; Extended Data Fig. 6c,e). Also similar to LiCl-induced malaise, CGRP neuron stimulation elicited stronger activation of the amygdala network when preceded by novel, rather than familiar, flavor consumption (Fig. 3j,k; Extended Data Fig. 6d,f). This effect was especially prominent in the CEA (Fig. 3h) and not observed in brain regions outside of the amygdala network (Fig. 3j; Extended Data Fig. 6d,f).

We next asked if the stronger effect of CGRP neuron stimulation on amygdala activation following novel versus familiar flavor consumption could be explained by recruitment of a specific CEA cell type. To address this, we performed multiplex fluorescence in situ hybridization (FISH) in the CEA in a separate cohort of mice that consumed a novel or familiar flavor followed by delayed CGRP^CEA^ projection stimulation and examined co-expression of *Fos* with markers for known CEA cell types^63–67^ (*Sst*, *Prkcd*, *Calcrl*). Most cells that expressed *Fos* also expressed *Prkcd* or *Calcrl*, and this did not depend on whether the mice had consumed a novel or familiar flavor (Fig. 3l–n; Extended Data Fig. 6g,h).

Taken together, these experiments show that postingestive CGRP neuron activity is necessary and sufficient to mediate delayed CTA, as well as the effects of malaise on novel flavor-dependent, amygdala-specific neural activation. This novelty-dependent activation does not appear to be instantiated by a specific CEA cell type, and instead novel (versus familiar) flavor consumption leads to a greater number of *Prkcd*^+^/*Calcrl*^+^ CEA cells being activated by subsequent CGRP neuron activity.

### Postingestive malaise specifically reactivates novel flavor-coding neurons in the amygdala, and this effect is mediated by CGRP neurons

While our brainwide Fos measurements and CGRP neuron manipulations point to the amygdala network as a unique site for the convergence of flavor representation and delayed malaise signals, these measurements do not resolve how these temporally separated signals are integrated at the single-cell level (Fig. 4a). One possibility is that individual novel flavor-coding neurons may be persistently activated long after a meal in a manner that provides passive overlap with delayed CGRP neuron malaise signals (Hypothesis 1 in Fig. 4a). Alternatively, CGRP neuron inputs may specifically reactivate novel flavor-coding neurons (Hypothesis 2 in Fig. 4a). Another possibility is that CGRP neuron inputs may activate a separate population of amygdala neurons that subsequently becomes incorporated into the novel flavor representation during memory consolidation (Hypothesis 3 in Fig. 4a). Testing these hypotheses requires tracking the activity of the same neurons across the stages of learning.

**Fig. 4.**
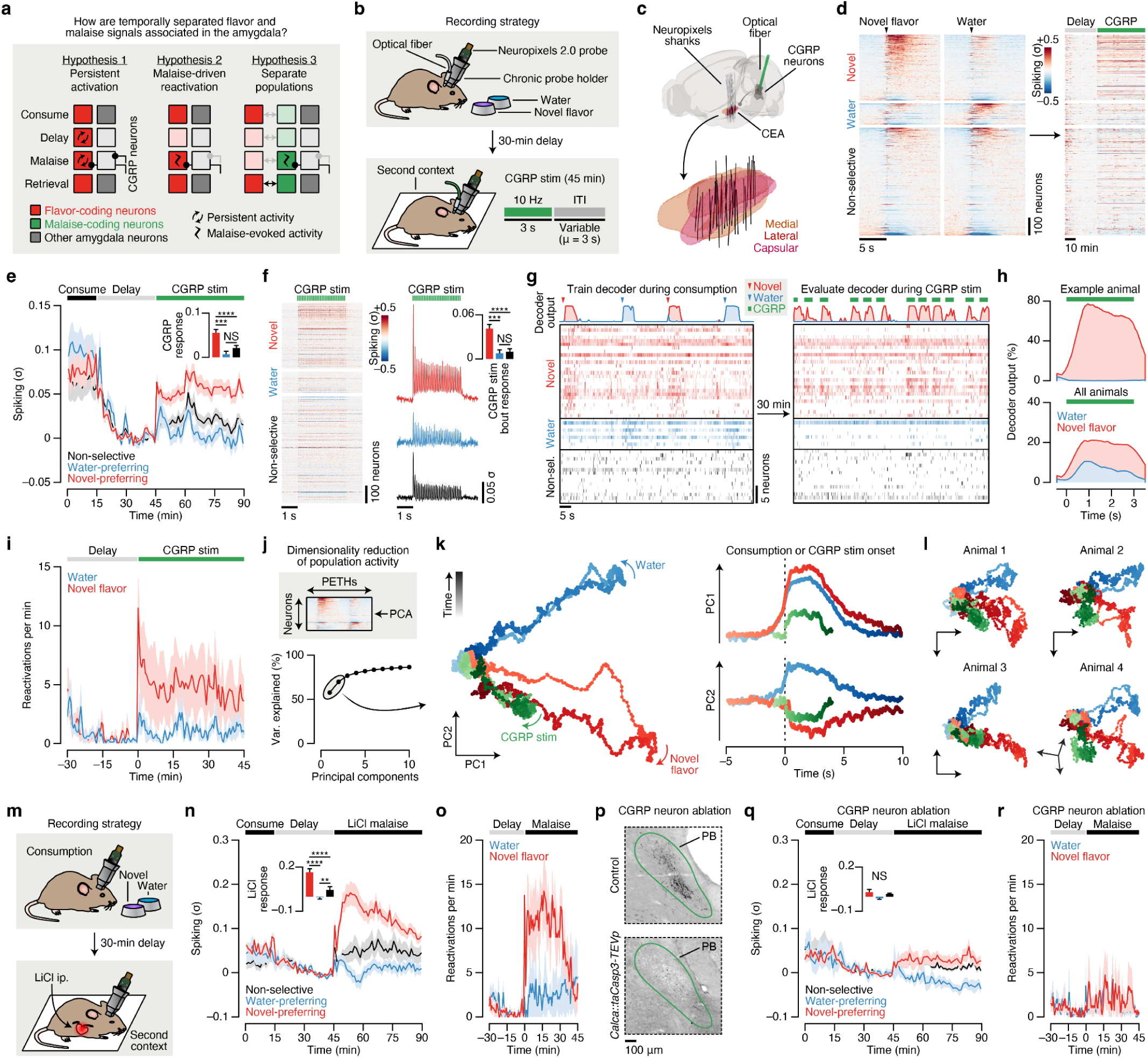
Postingestive CGRP neuron activity preferentially reactivates the representation of a recently consumed novel flavor in the amygdala. **a,** Hypotheses for how the amygdala associates temporally separated flavor and malaise signals to support CTA learning. Hypothesis 1: Individual novel flavor-coding neurons may be persistently activated long after a meal, either through cell-autonomous or circuit mechanisms, in a manner that provides passive overlap with delayed CGRP neuron malaise signals. Hypothesis 2: CGRP neuron inputs may specifically reactivate novel flavor-coding neurons. Hypothesis 3: CGRP neuron inputs may specifically activate a separate population of neurons that subsequently becomes incorporated into the novel flavor representation upon memory retrieval. Each of these hypotheses provides a plausible mechanism for linking flavors to malaise signals, but they make mutually exclusive predictions about single-neuron and populational-level activity across the stages of learning. **b,** Schematic of the CTA paradigm for chronic Neuropixels recordings and CGRP neuron stimulation. We trained mice to consume the novel flavor and water at relatively equal rates, and the total amount consumed was equal (Extended Data Fig. 8a; see Methods). **c,** Reconstruction of recording trajectories registered to the Allen CCF. Each line represents one shank of a four-shank Neuropixels 2.0 probe targeting CEA (*n* = 32 shanks from 8 mice). **d,** Heatmap showing the trial-average spiking of all recorded CEA neurons (*n* = 1,104 single- and multi-units from 8 mice) to novel flavor and water consumption during the consumption period (left) and during the delay and CGRP neuron stimulation periods (right). Neurons are grouped by their novel flavor/water preference and then sorted by consumption response magnitude. Consumption PETHs are time-locked to delivery of the flavor or water, which was triggered by the animal entering the port. **e,** Average spiking of the novel flavor-preferring (red; *n* = 373 neurons), water-preferring (blue; *n* = 121 neurons), and non-selective (black; *n* = 610 neurons) populations across the entire experiment. The inset quantifies the average response of each population during the entire 45-min CGRP neuron stimulation period. **f,** Left: Heatmap showing the trial-average spiking of all recorded neurons to individual 3-s bouts of 10-Hz CGRP neuron stimulation. Right: Average spiking of the novel flavor-preferring (red), water-preferring (blue), and non-selective (black) populations during CGRP neuron stimulation bouts. The inset quantifies the average response of each population within bouts of CGRP neuron stimulation. **g,** Example multinomial logistic regression decoder session. The top row shows the moment-by-moment decoder posterior for the novel flavor (red) and water (blue). The raster below shows time-locked neural activity for novel flavor-preferring, water-preferring, and non-selective neurons. The symbols in the legend represent the true event times (novel flavor delivery, water delivery, CGRP neuron stimulation), not decoder predictions. Only a subset of recorded neurons is shown for clarity (50 out of 90). **h,** Average decoder posterior time-locked to CGRP neuron stimulation for the example animal (top) and across all mice (bottom; *n* = 6 mice). **i,** Average reactivation rate for the novel flavor and water across the delay and CGRP neuron stimulation periods (*n* = 6 mice). We defined a reactivation event as any peak in the decoder posterior trace that was > 0.5. **j,** Top: Schematic of the population activity dimensionality reduction analysis. Bottom: The first two principal components explained >70% of the variance in trial-average population dynamics during novel flavor and water consumption. **k,** Left: Neural trajectories for novel flavor consumption (red), water consumption (blue), and CGRP neuron stimulation in PC-space. Right: Time-courses along the PC1 and PC2 axes for the trajectories to left. **l,** Dimensionality reduction analysis performed separately for four individual example mice, rather than on all mice combined as in **k**. **m,** Schematic of the CTA paradigm for chronic Neuropixels recordings and LiCl-induced malaise. **n,** Average spiking of the novel flavor-preferring (red; *n* = 280 neurons from 4 mice), water-preferring (blue; *n* = 80 neurons), and non-selective (black; *n* = 218 neurons) populations across the entire experiment. The inset quantifies the average response of each population following LiCl injection. **o**, Average population-level reactivation rate for the novel flavor and water across the delay and malaise periods (*n* = 4 mice). **p**, Example CGRP immunoreactivity data confirming genetic ablation of CGRP neurons by taCasp3-TEVp. See also Extended Data Fig. 9f,g. **q**, Analogous to **n**, but for mice with CGRP neuron ablation (*n* = 124 novel flavor-preferring neurons, 20 water-preferring neurons, 256 non-selective neurons from 4 mice). **r,** Analogous to **o**, but for mice with CGRP neuron ablation (*n* = 4 mice). *P*-values in **e,f,n,q** are from Wilcoxon rank-sum tests corrected for multiple comparisons across neuron groups. Error bars and shaded areas represent mean ± s.e.m. \*\**P* ≤ 0.01, \*\*\**P* ≤ 0.001, \*\*\*\**P* ≤ 0.0001, NS, not significant (*P* > 0.05).

Therefore, to distinguish between these possibilities, we performed high density recordings from individual neurons in CEA — the core node in our amygdala network (Figs. 1–3) — during consumption, subsequent malaise, and memory retrieval (Fig. 4b). Recordings were performed with chronically implanted four-shank Neuropixels 2.0 probes^68,69^ (Extended Data Fig. 7a). Reconstruction of individual shank trajectories confirmed that we were able to precisely target CEA (Fig. 4c; Extended Data Fig. 7b,c; 138 ± 30 CEA neurons/animal; mean ± s.e.m.). We initially trained mice to consume water at an equal rate from two port locations (Extended Data Fig. 8a; see Methods). On conditioning day, we replaced one location with a novel flavor, while water remained in the other as an internal control. Immediately after the consumption period ended, mice were transferred to a distinct second context where they would experience postingestive malaise, to ensure that any neural correlates of flavor consumption that we might subsequently observe were not due to features of the original context where consumption occurred. After a 30-min delay period in this second context, we induced postingestive malaise using either (1) optogenetic CGRP neuron cell-body stimulation (Fig. 4b–l), (2) optogenetic CGRP^CEA^ projection stimulation (Extended Data Fig. 9a–e), or (3) LiCl injection (Fig. 4m–r), across three cohorts of mice. While all three cohorts gave similar results, we will begin by describing the data from the first cohort.

During consumption, 34% of CEA neurons were significantly activated by the novel flavor, compared to only 11% for water (Fig. 4d; Extended Data Fig. 8b) and, in a separate experiment described later, 17% for a familiar flavor. These observations are consistent with our Fos data at the consumption timepoint (Fig. 1e). Almost all CEA neurons, including novel flavor-coding neurons, were significantly less active after consumption ended (Fig. 4d,e), which suggests that persistent activation (Hypothesis 1; Fig. 4a) is not the mechanism that the amygdala network uses to associate flavors with delayed malaise signals.

We next investigated how delayed CGRP neuron stimulation impacted CEA neuron activity. Remarkably, CGRP neuron stimulation potently and specifically reactivated novel flavor-coding CEA neurons, with only limited effects on water-coding and non-selective neurons (Fig. 4d,e; consistent only with Hypothesis 2; Fig. 4a). This reactivation was precisely time-locked to individual bouts of CGRP neuron stimulation (Fig. 4f), which suggests that it is directly driven by the release of glutamate from CGRP neuron inputs^59,60^ (Fig. 3c; Extended Data Fig. 5c) rather than by a slow change in affective or physiological internal state. Similar to CGRP neuron cell-body stimulation, delayed CGRP^CEA^ projection stimulation robustly reactivated novel flavor-coding CEA neurons (Extended Data Fig. 9a–d).

Reactivation of novel flavor-coding neurons was similarly present in a separate cohort of mice that experienced delayed LiCl-induced malaise rather than CGRP neuron stimulation (Fig. 4m,n). Moreover, genetic ablation^70^ of CGRP neurons abolished the preferential reactivation of novel flavor-coding neurons by delayed malaise (Fig. 4p,q) and impaired learning (Extended Data Fig. 9f,g). This indicates that the effects of postingestive malaise on CTA — and on CEA dynamics — are mediated by CGRP neurons.

Together, these observations suggest that neurons encoding a recently consumed novel flavor in CEA are specifically reactivated by delayed malaise signals encoded by CGRP neuron inputs, providing a potential mechanism for temporal credit assignment during CTA learning (Hypothesis 2; Fig. 4a).

### Postingestive CGRP activity specifically reactivates population-level novel flavor representations in the amygdala

Population-level analyses corroborated the conclusion that CGRP neuron activity after consumption preferentially reactivates the neural representation of the recently consumed novel flavor. First, we trained a multinomial logistic regression decoder using population activity during the consumption period to discriminate novel flavor or water consumption from baseline activity (Fig. 4g; Extended Data Fig. 8c,d). Cross-validated decoding accuracy was nearly perfect (Extended Data Fig. 8e). We then evaluated this decoder using population activity during the delay and CGRP neuron stimulation periods, and investigated the probability of decoding the novel flavor or water representation on a moment-by-moment basis (decoder output, *P*(novel flavor|population activity)). Strikingly, this decoding analysis showed that individual bouts of CGRP neuron stimulation reliably reactivated population-level novel flavor representations (Fig. 4g–i). In contrast, water representations were rarely reactivated by CGRP neuron stimulation (Fig. 4g–i). Consistent with these results from CGRP neuron cell-body stimulation, we found that LiCl-induced malaise also strongly reactivated population-level novel flavor representations in CEA (Fig. 4o) and that this required functional CGRP neurons (Fig. 4r).

We next compared population activity trajectories during consumption and during CGRP neuron stimulation by performing PCA on the pooled (across mice) trial-averaged population activity during novel flavor and water consumption. Neural activity during consumption was low-dimensional, with the first two PCs explaining >70% of variance in trial-averaged population activity (Fig. 4j). Plotting neural trajectories during novel flavor and water consumption on the PC1–PC2 axis revealed that PC2 perfectly discriminated these two flavors (Fig. 4k). Remarkably, using these PCA loadings to project the population activity during CGRP neuron stimulation onto the same PC1–PC2 axis showed that this experience closely mirrored the neural trajectory of novel flavor consumption (Fig. 4k). This strong effect was also apparent in the population activity of individual mice (Fig. 4l) and in population activity from mice that received delayed CGRP^CEA^ projection stimulation (Extended Data Fig. 9e). Thus, population analyses confirmed that CGRP neuron activity specifically reactivates population-level novel flavor representations in the CEA during delayed postingestive malaise (Hypothesis 2; Fig. 4a).

### Postingestive CGRP activity induces plasticity to stabilize flavor representations in the amygdala

Tracking neural activity across flavor consumption and malaise revealed that malaise signals preferentially reactivate novel flavor representations in the CEA (Fig. 4), providing a potential mechanism for the brain to link flavors experienced during a meal with delayed postingestive feedback. If this mechanism contributes to learning, then postingestive CGRP neuron activity would be expected to trigger functional plasticity in amygdala flavor representations that could underlie the CTA memory. To test this hypothesis, we next examined whether postingestive reactivation of flavor-coding CEA neurons is predictive of stronger flavor responses when the CTA memory is retrieved. To accomplish this, we took advantage of the high stability of our chronic recordings to track the same CEA neurons across days and analyze their responses to flavor consumption before (conditioning day) and after (retrieval day) pairing with CGRP neuron stimulation (Fig. 5a,b; see Methods).

**Fig. 5.**
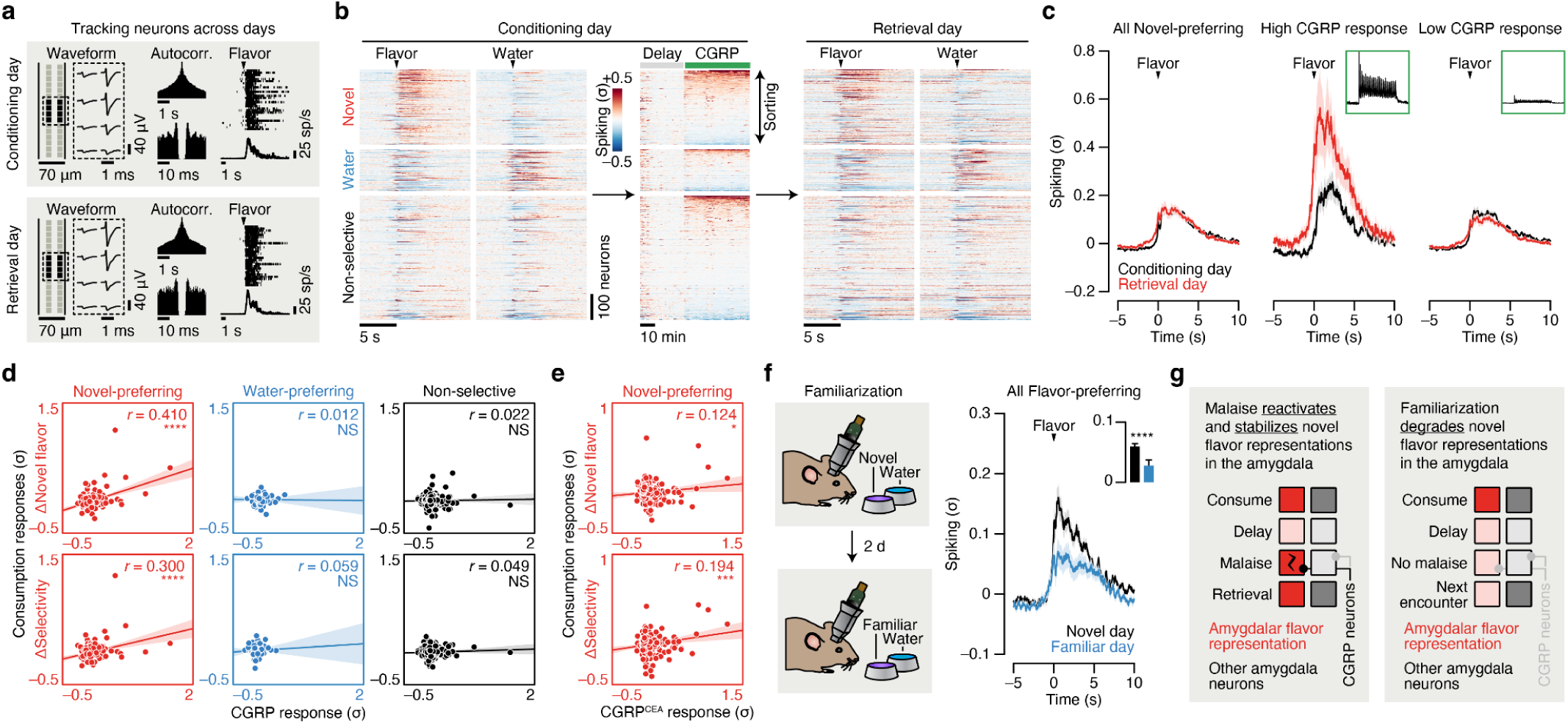
Postingestive CGRP neuron activity induces plasticity to stabilize novel flavor representations in the amygdala upon memory retrieval. **a,** Spike waveforms, autocorrelograms, and flavor response rasters for one example neuron tracked across conditioning and retrieval days. **b,** Heatmap showing the average spiking of all recorded neurons (*n* = 939 neurons from 8 mice) to novel flavor and water consumption during the consumption period (left) and during the delay and CGRP neuron stimulation periods (middle) on conditioning day, and the responses of the same neurons to flavor and water consumption on retrieval day. Neurons are grouped by their novel flavor/water preference on pairing day and then sorted by CGRP response magnitude. **c,** Left: Trial-average spiking of the novel flavor-preferring population (*n* = 265 neurons) during flavor consumption on conditioning day (black) and retrieval day (red). Middle/Right: Trial-average spiking of the novel-preferring neurons with the highest 10% of CGRP response magnitudes (High CGRP response; middle) and of the remaining novel-preferring neurons (Low CGRP response; right) on conditioning day and retrieval day. The insets show the average CGRP neuron stimulation response profile of each subpopulation. **d,** Correlation between each neuron’s change (Retrieval – Conditioning) in flavor response (top) or selectivity (bottom) during the consumption period to its average response during the CGRP neuron stimulation period. The novel flavor-preferring (left; *n* = 265 neurons), water-preferring (middle; *n* = 123 neurons), and non-selective (right; *n* = 551 neurons) populations are shown separately. **e**, Analogous to **d**, but for mice with CGRP^CEA^ projection stimulation rather than cell-body stimulation (*n* = 286 novel flavor-preferring neurons from 8 mice). See Extended Data Fig. 10c–e for additional analysis. **f,** Left: Schematic for the flavor familiarization experiment. Right: Trial-average spiking of the initially flavor-preferring population (*n* = 201 neurons from 7 mice; classified on novel day) during flavor consumption on novel day (black) and familiar day (blue). The inset quantifies the average response on each day. See Extended Data Fig. 10h–k for additional analysis. **g,** Illustration of a neural mechanism for learning from delayed postingestive feedback using malaise-driven reactivation and stabilization of amygdala novel flavor representations. *P*-values in **d,e** are from Pearson correlation *t*-tests. *P*-value in **f** is from a Wilcoxon signed-rank test. Error bars represent mean ± s.e.m. Shaded areas in **c,f** represent mean ± s.e.m. and in **d,e** represent 95% confidence interval for linear fit. \**P* ≤ 0.05, \*\*\**P* ≤ 0.001, \*\*\*\**P* ≤ 0.0001, NS, not significant (*P* > 0.05).

We found that the trial-averaged response of novel flavor-coding CEA neurons (classified on conditioning day) was largely stable across days (Fig. 5c). However, sorting these neurons based on the magnitude of their response to CGRP neuron stimulation revealed a striking effect: novel flavor-coding CEA neurons with the greatest CGRP neuron input responded much more strongly to the flavor during memory retrieval, whereas the responses of novel flavor-coding neurons with weak or no CGRP neuron input remained relatively unchanged (Fig. 5c,d). In contrast, we did not observe a similar correlation for water-coding or non-selective CEA neurons (Fig. 5d), which further suggests that postingestive malaise does not recruit additional neurons into the initial novel flavor representation (as in Hypothesis 3; Fig. 4a). Together, these observations indicate that CGRP neurons induce functional plasticity that stabilizes the amygdala’s response to the conditioned flavor after learning. Consistent with this conclusion, we found that the population-level flavor representation, as visualized using PCA, was remarkably stable across conditioning and retrieval days (Extended Data Fig. 10b,l).

CGRP^CEA^ projection stimulation (Fig. 5e; Extended Data Fig. 10c–e) and LiCl-induced malaise (Extended Data Fig. 10f) had similar stabilizing effects on novel flavor-coding neuron responses during memory retrieval as CGRP neuron cell-body stimulation. In contrast, the responses of novel flavor-coding neurons significantly decreased during memory retrieval in mice lacking CGRP neurons (Extended Data Fig. 10g). Thus, CGRP neuron activity is necessary and sufficient for both the reactivation and stabilization of amygdala flavor representations by delayed malaise signals.

For comparison, we next asked how amygdala activity evolves after familiarization (that is, experience with a flavor without any aversive postingestive consequences, as in the “Familiar” condition in Figs. 1–3). Consistent with our initial Fos data comparing novel versus familiar flavor activation patterns (Fig. 1e; Fig. 2f,g; Fig. 3h,k), tracking CEA neuron activity before and after familiarization revealed that the proportion of flavor-coding neurons significantly decreased after familiarization (Extended Data Fig. 10h,i). Similarly, we found that the trial-averaged response of individual flavor-coding neurons (classified on novel day) significantly decreased after familiarization (Fig. 5f), in contrast to the stability we observed following conditioning with CGRP neuron stimulation (Fig. 5c, left) and LiCl-induced malaise (Extended Data Fig. 10f). Furthermore, we found that initially water-preferring neurons increased their response to the flavor after it became familiar (Extended Data Fig. 10j). Together, these observations suggest that familiarization degrades amygdala flavor representations such that the representation of a flavor moves closer to the representation of pure water. Consistent with this conclusion, we found that the population-level representation of the flavor along the PC2 dimension that discriminates flavor from water was almost completely abolished following familiarization (Extended Data Fig. 10k,l).

Thus, CGRP neurons convey malaise signals that preferentially reactivate novel flavor representations in the amygdala, thereby enabling the brain to bridge the delay between a meal and postingestive feedback during CTA. Moreover, these postingestive signals induce plasticity to stabilize novel flavor representations after conditioning, whereas flavor representations rapidly degrade in the absence of malaise signals as flavors become familiar and safe (Fig. 5g).

### Novel flavor consumption triggers PKA activity in the amygdala, providing a potential biochemical eligibility trace for reactivation by postingestive CGRP activity

While so far we have focused on the role of neural activity in supporting CTA, previous work in the field has taken a complementary approach to examine the role of biochemical signals^71–79^. For example, many studies have shown an important role in CTA for cAMP response element-binding protein^26,34,76–79^ (CREB), a transcription factor that regulates neural excitability^80–82^, and protein kinase A^83,84^ (PKA), which phosphorylates and activates CREB. Expression of CREB at the time of conditioning is thought to bias neurons to become part of the CTA “memory engram”^85,86^: the ensemble of cells that are activated during retrieval of the CTA memory (Fig. 6a).

**Fig. 6.**
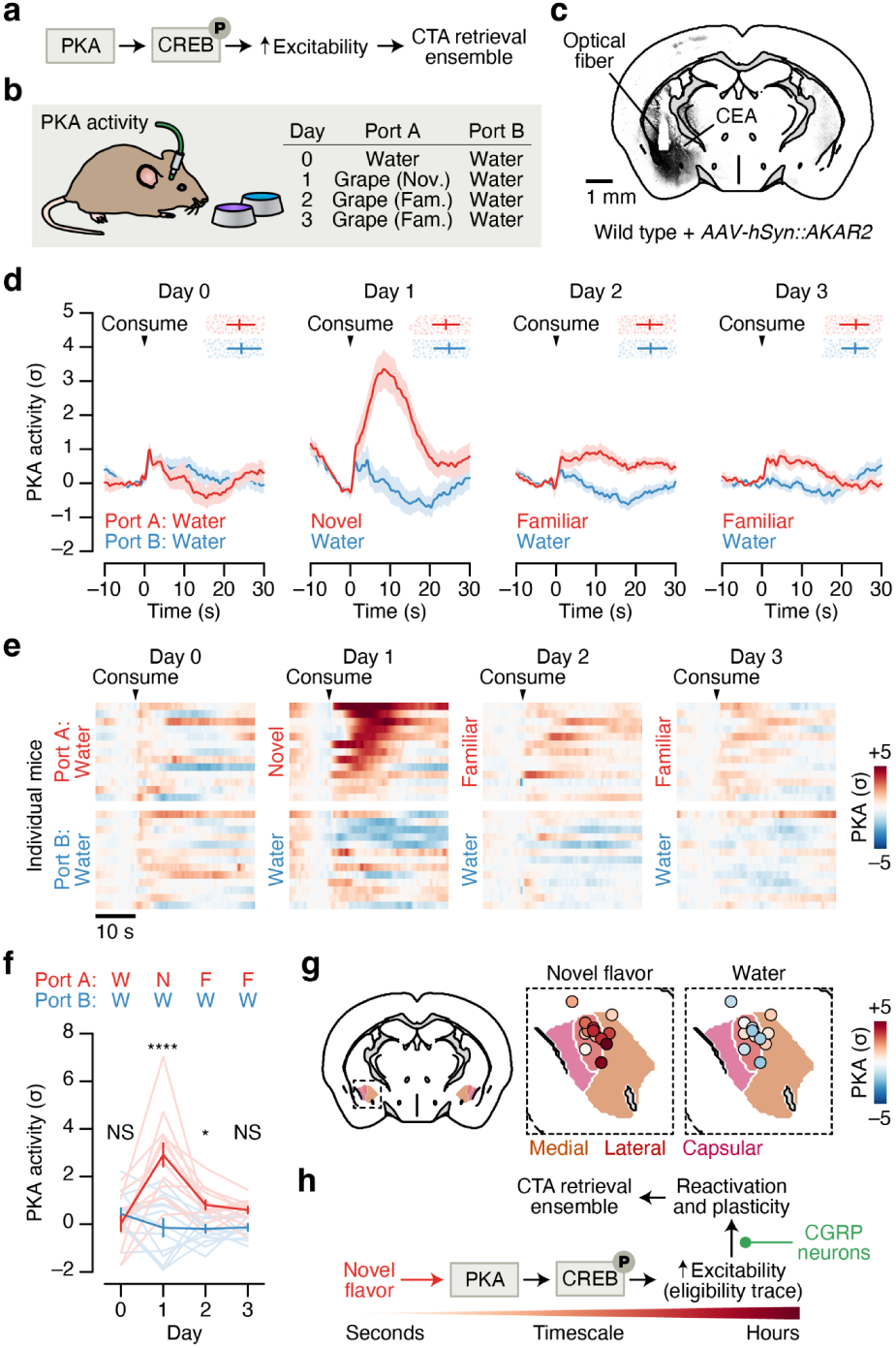
Novel flavor consumption triggers PKA activity in the amygdala, providing a potential biochemical eligibility trace for reactivation by postingestive CGRP neuron activity. **a,** Simplified schematic of the biochemical pathway that has been proposed to recruit a subset of amygdala neurons into the CTA memory retrieval ensemble (“memory engram”)^76–79,83–86^. **b**, Schematic for recording PKA activity in the CEA across familiarization using the AKAR2 sensor^87^. **c**, Example AKAR2 expression in the CEA. **d**, PKA activity in the CEA in response to consumption of a novel or familiar flavor (Port A; red) and to water (Port B; blue) across four consecutive days (*n* = 13 mice). Points and error bars at the top of each plot indicate the timing of the next reward consumption. **e**, PKA activity for individual mice in response to novel or familiar flavor and to water consumption. Mice are sorted by novel flavor response (Day 1). **f**, Summary of PKA activity in response to novel or familiar flavor and to water consumption (*n* = 13 mice). **g**, PKA activity in response to novel flavor (left) and water (right) consumption on Day 1 for individual mice aligned to the Allen CCF (*n* = 13 mice). **h**, Schematic of a putative hypothesis linking biochemistry with neural activity during CTA, with novel flavor-dependent increases in PKA in amygdala neurons leading to increased reactivation by delayed CGRP neuron inputs and recruitment into the CTA memory retrieval ensemble. *P*-values in **f** are from GLMM marginal effect *z*-tests corrected for multiple comparisons across days. Error bars and shaded areas represent mean ± s.e.m. \**P* ≤ 0.05, \*\*\*\**P* ≤ 0.0001, NS, not significant (*P* > 0.05).

How might the PKA → CREB pathway relate to the malaise-driven reactivation and stabilization of novel flavor representations that we describe here? One possibility is that this pathway could be preferentially triggered in the amygdala by novel flavor consumption, which may in turn contribute to increased excitability or responsiveness of novel flavor-coding neurons to CGRP neuron inputs during malaise.

To test the first part of this hypothesis — that the PKA → CREB pathway is preferentially activated by novel flavors — we recorded in vivo PKA activity in the CEA using fiber photometry of the new sensor AKAR2^87^ (Fig. 6b,c). This showed that novel flavors drive a robust increase in PKA activity in the CEA, whereas familiar flavors have little impact on PKA (Fig. 6d–g). This increase in PKA activity (tens of seconds) is substantially longer in duration than the increase in spiking for each bout of consumption (Fig. 4d) — and downstream effects on CREB, gene expression, and neural excitability are presumably far longer-lasting. Thus, novelty-dependent gating of PKA could serve as a biochemical eligibility trace that increases the responsiveness of novel flavor-coding neurons to delayed malaise signals, thereby permitting the selective reactivation and stabilization of amygdala novel flavor representations (Fig. 6h).

## Discussion

The major reason that learning is challenging is because of delays between a stimulus or action and its outcome. How does the brain assign credit to the correct prior event? Most work on credit assignment has been limited to examining learning in the case of relatively short delays (on the order of seconds)^88–97^. Postingestive learning paradigms, such as in CTA, offer an opportunity to study how the brain assigns credit across much longer delays.

Here we have described a neural mechanism that may contribute to solving the credit assignment problem inherent to CTA: postingestive malaise signals specifically reactivate the neural representations of novel flavors experienced during a recent meal, and this reactivation serves to stabilize the flavor representation upon memory retrieval.

While previous work, mostly in the hippocampus^98–100^ and cortex^101–103^, has suggested a role for neural reactivations in memory^104–106^, our study advances this idea in multiple important ways. First, we bring this concept to an entirely new paradigm (postingestive learning and CTA). Second, we discover a role for the outcome signal (unconditioned stimulus) in directly triggering reactivations and find that a cell-type-specific malaise pathway mediates this effect. Third, we discover a strong relationship between outcome-driven flavor reactivations and strengthened flavor representations during memory retrieval. Finally, we describe a potential biochemical eligibility trace (PKA → CREB activity) that may facilitate preferential reactivation and stabilization of flavor representations by delayed outcome signals.

Our entry point into this problem was the fact that novel flavors more easily support postingestive learning than familiar flavors^15–18^. By comparing brainwide neural activation patterns in animals that consumed the same flavor when it was novel versus familiar, we identified an amygdala network that was unique in preferentially responding to novel flavors across every stage of learning. While we focused here on how novel flavors become associated with aversive postingestive feedback in this amygdala network, it is possible that these same ideas generalize to the processes of familiarization or postingestive nutrient learning as well. Specifically, a parallel and mechanistically similar process may be at work when learning that a food is safe or nutritious. In that case, “safety” (or “reward”^7–14,107–109)^ signals may reactivate recently consumed novel flavor representations, rather than the aversive CGRP neuron-mediated reactivations we report here. This process may mediate the weakening of flavor representations in the amygdala network and/or the strengthening of flavor representations in the LS and other limbic regions (Fig. 1f).

Much previous research on CTA has concentrated on the role of the gustatory insular cortex^29,30,110–112^. A consistent finding is that CTA amplifies the cortical representation of the conditioned taste and shifts it to be more similar to innately aversive tastants. Recent work has further shown that homeostatic synaptic plasticity contributes to a transition, over the course of hours or days, from the initial formation of a more generalized taste aversion to a taste-specific CTA memory^113,114^. How the malaise-driven reactivation of amygdala flavor representations we report here relates to such mechanisms in the insula is an open question. One possibility is that the reactivation of novel flavor representations by delayed malaise signals that we report contributes to an initial memory, which is then further refined through homeostatic mechanisms in the insula and its reciprocal connections with the amygdala^29,31,34,115–117^.

Overall, our results demonstrate how dedicated novelty detection circuitry and built-in priors (preferential reactivation of recent flavor representations during malaise) work together to enable the brain to properly link actions and outcomes despite long delays.

## Methods

### Animals and surgery

All experimental procedures were approved by the Princeton University Institutional Animal Care and Use Committee following the NIH Guide for the Care and Use of Laboratory Animals. Wild type mice (JAX 000664) and *Calca::Cre* mice^59^ (JAX 033168) were obtained from the Jackson Laboratory. Adult mice (>8 weeks) of both sexes were used for all experiments. Mice were housed under a 12-hr/12-hr light/dark cycle, and experiments were conducted during the dark cycle. Stereotaxic surgeries were performed under isoflurane anesthesia (3–4% for induction, 0.75–1.5% for maintenance). Mice received preoperative antibiotics (5 mg/kg Baytril subcutaneous, sc.) and pre- and post-operative analgesia (10 mg/kg Ketofen sc.; three daily injections). Post-operative health (evidence of pain, incision healing, activity, posture) was monitored for at least five days. For all CTA experiments, mice were water-restricted and maintained at >80% body weight for the duration of the experiment.

### Viral injections

For CGRP neuron stimulation experiments (Figs. 3–5), we bilaterally injected 400 nl of AAV5-EF1a-DIO-hChR2(H134R)-eYFP (titer: 1.2e13 genome copies (GC)/ml; manufacturer: Princeton Neuroscience Institute (PNI) Viral Core Facility) at –5.00 mm AP, ±1.40 mm ML, –3.50 mm DV into *Calca::Cre* mice. We used this stereotaxic coordinate to target the parabrachial nucleus (PB) in all subsequent experiments. For CGRP neuron fiber photometry experiments (Fig. 3b), we unilaterally injected 400 nl of AAV9-hSyn-FLEX-GCaMP6s (titer: 1.0e13 GC/ml; manufacturer: PNI Viral Core Facility) into the PB of *Calca::Cre* mice. For CGRP neuron → CEA (CGRP^CEA^) projection stimulation experiments (Figs. 3–5), we bilaterally injected 350 nl of AAV5-EF1a-DIO-hChR2(H134R)-eYFP (titer: 1.2e13 GC/ml; manufacturer: PNI Viral Core Facility; RNAscope FISH experiment), AAV5-EF1a-DIO-ChRmine-mScarlet (titer: 9.0e12 GC/ml; manufacturer: PNI Viral Core Facility; all other experiments), or AAV5-EF1a-DIO-eYFP (titer: 1.5e13 GC/ml; manufacturer: PNI Viral Core Facility) into the PB of *Calca::Cre* mice. For CGRP^CEA^ projection inhibition experiments (Fig. 3f), we bilaterally injected 350 nl of AAV5-hSyn-SIO-eOPN3-mScarlet (titer: 9.0e12 GC/ml; manufacturer: Addgene) or AAV5-EF1a-DIO-eYFP (titer: 1.5e13 GC/ml; manufacturer: PNI Viral Core Facility) into the PB of *Calca::Cre* mice. For CGRP neuron ablation experiments (Fig.4p–r; Extended Data Fig. 9g,h), we bilaterally injected 350 nl of AAV5-EF1a-FLEX-taCasp3-TEVp (titer: 1.6e13 GC/ml; manufacturer: Addgene) into the PB of *Calca::Cre* mice. For control LiCl conditioning Neuropixels mice (Fig. 4m–o), we bilaterally injected 350 nl of AAV5-Camk2a-eYFP (titer: 7.5e11 GC/ml; manufacturer: University of North Carolina (UNC) Vector Core) into the PB of wild type mice. For CEA PKA recording experiments (Fig. 6), we unilaterally injected 300 nl of AAV5-hSyn-ExRai-AKAR2 (2.4e13 GC/ml; manufacturer: PNI Viral Core Facility) at –1.15 mm AP, –2.65 mm ML, –4.85 mm DV into wild type mice. For LS activation experiments (Extended Data Fig. 2), we bilaterally injected AAV5-hSyn-hM3D(G_q_)-mCherry (titer: 3.8e12 GC/ml; manufacturer: Addgene) or AAV5-Camk2a-eYFP (titer: 7.5e11 GC/ml; manufacturer: UNC Vector Core) at one (500 nl at +0.55 mm anterior-posterior (AP), ±0.35 mm medial-lateral (ML), 4.00 mm dorsal-ventral (DV)) or two (150 nl each at +0.85 mm/+0.25 mm AP, ±0.60 mm ML, 3.75 mm DV) coordinates into wild type mice. Virus was infused at 100 nl/min. All coordinates are given relative to bregma. We allowed three weeks for AKAR2 and GCaMP expression, at least four weeks for ChR2 and hM3D expression, five weeks for CGRP neuron ablation by taCasp3-TEVp, and eight weeks for CGRP^CEA^ terminal expression of ChRmine, ChR2, and eOPN3.

### Optical fiber implantations

Optical fibers encased in stainless steel ferrules were implanted in the brain for optogenetic and fiber photometry experiments. For bilateral optogenetic stimulation of CGRP neurons (Fig. 3), we implanted ⌀300-µm core/0.39-NA fibers (Thorlabs FT300EMT) above the PB at a 10° angle with the fiber tips terminating 300–400 µm above the viral injection coordinate. For unilateral stimulation of CGRP neurons (Figs. 4,5), we implanted a ⌀300-µm core/0.39-NA fiber above the left PB at a 25–30° angle with the fiber tip terminating 300–400 µm above the viral injection coordinate. For bilateral optogenetic manipulation of CGRP^CEA^ projections (Fig. 3), we implanted ⌀300-µm core/0.39-NA fibers (Thorlabs FT300EMT) above the CEA with the fiber tips terminating at –1.15 mm AP, ±2.85 mm ML, –4.25 mm DV. For unilateral optogenetic stimulation of CGRP^CEA^ projections (Figs. 4,5), we implanted a ⌀300-µm core/0.37-NA fiber (MFC_300/360-0.37_10mm_MF2.5_FLT) above the left CEA at a +55° angle with the fiber tip terminating at –1.20 mm AP, +2.25 mm ML, –3.55 mm DV. For fiber photometry recording of CGRP neurons (Fig. 3b), we implanted a ⌀400-µm core/0.48-NA fiber (Doric MFC_400/430-0.48_5.0mm_MF2.5_FLT) above the left PB at a –10° to –30° angle with the fiber tip terminating approximately at the viral injection coordinate. For fiber photometry recording of CEA PKA activity (Fig. 6), we implanted a ⌀400-µm core/0.48-NA fiber (Doric MFC_400/430-0.48_6.0mm_MF2.5_FLT) above the left CEA with the fiber tip terminating approximately at the viral injection coordinate. Optical fibers were affixed to the skull with Metabond (Parkell S380) which was then covered in dental cement.

### Chronic Neuropixels assembly

We used Neuropixels 2.0 probes^68^ (test-phase, four-shank; Imec), as they were miniaturized to make them more suitable for chronic implantation in small rodents. In order to avoid directly cementing the probes such that they could be reused, we designed a chronic implant assembly (Extended Data Fig. 7a) based on the design for Neuropixels 1.0 probes previously validated in rats^69^. Like that design, the assembly was printed on Formlabs SLA 3D printers and consisted of four discrete parts: (1) a *dovetail adapter* permanently glued to the probe base; (2) an *internal holder* that mated with the dovetail adapter and allowed stereotaxic manipulation of the probe; and (3–4) an *external chassis*, printed in two separated parts, that encased and protected the entire assembly. The external chassis and internal holder were attached using screws that could be removed at the end of the experiment to allow explantation and reuse. The external chassis of the final implant assembly was coated with Metabond before implantation. After explantation, probes were cleaned with consecutive overnight washes in enzyme-active detergent (Alconox Tergazyme) and silicone cleaning solvent (Dowsil DS-2025) prior to reuse. The dimensions of the Neuropixels 2.0 implant assembly were significantly smaller, primarily because of the smaller size of the probe and headstage. The maximum dimensions were 24.7 mm (height), 12.2 mm (width), and 11.2 mm (depth), with a weight of 1.5 g (not including the headstage). Space was made for the headstage to be permanently housed in the implant, as opposed to the previous design in which the headstage was connected only during recording and was secured to a tether attached to the animal. This made connecting the animal for a recording significantly easier and obviated the need for a bulky tether that limits the animal’s movements. This change was made possible due to improvements in Neuropixels cable design, requiring fewer cables per probe and requiring less reinforcement of the cables during free movement. Design files and instructions for printing and assembling the chronic Neuropixels 2.0 implant are available at: https://github.com/agbondy/neuropixels_2.0_implant_assembly.

### Chronic Neuropixels surgery

This surgery was performed 3–4 weeks after AAV injection to allow time for viral expression and behavioral training. First, three craniotomies were drilled: one small craniotomy (⌀500 µm) above the left PB (approached at a –10° to –30° angle) or left CEA (approached at a +55° angle) for the optical fiber, another small craniotomy above the cerebellum for the ground wire, and one large craniotomy (1 mm × 2 mm) above the left CEA for the Neuropixels probe. Next, a single optical fiber was placed above the left PB or left CEA as described above. At this point, the optical fiber was affixed to the skull with Metabond and the exposed skull was covered with Metabond. Next, a prefabricated chronic Neuropixels assembly was lowered at 2.5 µm/s into the CEA using a µMp micromanipulator (Sensapex). The probe shanks were aligned with the skull’s AP axis with the most anterior shank tip terminating at –0.95 mm AP, –2.95 mm ML, –6.50 mm DV. Once the probe was fully lowered, the ground wire was inserted 1–2 mm into the cerebellum and affixed with Metabond. The CEA craniotomy and probe shanks were then covered with medical grade petroleum jelly, and Dentin (Parkell S301) was used to affix the chronic Neuropixels probe to the Metabond on the skull. The PB optical fiber and CEA Neuropixels probe were both placed in the left hemisphere because CGRP neuron projections are primarily ipsilateral^37,61^.

### One-reward CTA paradigm

In Figs. 1–3 we used a one-reward CTA paradigm using either a novel or familiar flavor. Experiments were performed in operant boxes (Med-associates) equipped with a single nosepoke port and light and situated in sound-attenuating chambers. The nosepoke port contained a reward delivery tube that was calibrated to deliver 20-µl rewards via a solenoid valve (Lee Technologies LHDA2433315H). Every behavioral session (training and conditioning) had the following basic structure: First, the mouse was allowed to acclimate to the chamber for 5 min. Then, the consumption period began and the light turned on to indicate that rewards were available. During this period, each nosepoke, detected by an infrared beam break with a 1-s timeout period, triggered the delivery of a single reward, and the period ended when 1.2 ml of reward was consumed or 10-min had passed. Then, the delay period began and lasted until 30-min after the beginning of the consumption period. During training sessions, mice were returned to the homecage after the end of the delay period.

Novel flavor condition mice first received four training days in the paradigm above with water as the reward and no LiCl or CGRP neuron stimulation. On conditioning day, sweetened grape Kool-Aid (0.06% grape + 0.3% saccharin sodium salt; Sigma S1002) was the reward. Familiar flavor condition mice had sweetened grape Kool-Aid as the reward for all four training days as well as the conditioning day.

On the LiCl conditioning day (Figs. 1,2), mice received an intraperitoneal (ip.) injection of LiCl (125 mg/kg; Fisher Scientific L121) or normal saline after the 30-min delay following the end of the consumption period. For CGRP neuron stimulation (Fig. 3d) and CGRP^CEA^ projection stimulation (Fig. 3e) experiments, mice then received 45-min of intermittent stimulation beginning after the 30-min delay. Blue light was generated using a 447-nm laser for ChR2 experiments. Green light was generated using a 532-nm laser for ChRmine experiments. The light was split through a rotary joint and delivered to the animal using ⌀200-µm core patch cables. Light power was calibrated to approximately 10 mW at the patch cable tip for ChR2 experiments and 3 mW for ChRmine experiments. During the experiment, the laser was controlled with a Pulse Pal signal generator (Sanworks 1102) programmed to deliver 5-ms laser pulses at 10 Hz. For the duration of the stimulation period, the laser was pulsed for 1.5–15-s intervals (randomly chosen from a uniform distribution with 1.5-s step size) and then off for 1–10-s intervals (randomly chosen from a uniform distribution with 1-s step size). For the eOPN3 experiment (Fig. 3f), photoinhibition began 1-min before the LiCl injection and then continued for 90 min (532-nm laser; 10-mW power 500-ms laser pulses at 0.4 Hz). Mice were then returned to the homecage. For the LS activation experiments (Extended Data Fig. 2), mice received an ip. injection injection of 3 mg/kg clozapine *N*-oxide (CNO; Hellobio 6149) 45-min before the experiment began.

We assessed learning using a two-bottle memory retrieval test. Two bottles were affixed to the side of mouse cages (Animal Care Systems Optimice) such that the sipper tube openings were located approximately 1-cm apart. One day after conditioning, mice were given 30-min access with water in both bottles. We calculated a preference for each mouse for this session, and then counterbalanced the location of the test bottle for the retrieval test such that the average water day preference for the two bottle locations was as close to 50% as possible for each group. The next day, mice were given 30-min access with water in one bottle and sweetened grape Kool-Aid in the other bottle. Flavor preference was then calculated using the weight consumed from each bottle during this retrieval test, Flavor/(Flavor+Water).

To initially characterize behavior in our CTA paradigm (Fig. 1b), retrieval tests were conducted on three consecutive days with the flavor bottle in the same location each day. We then fit a GLMM to this dataset using the glmmTMB R package^119^ (https://github.com/glmmTMB/glmmTMB) with a gaussian link function and the formula:

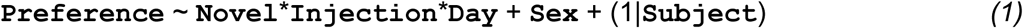

where **`Preference`** is the retrieval test result, **`Novel`** (Novel, Familiar), **`Injection`** (LiCl, Saline), **`Day`** (Day 1, Day 2, Day 3), and **`Sex`** (Female, Male) are fixed effect categorical variables, (1|**`Subject`**) is a random effect for each mouse, and * represents the main effects and interactions. This GLMM showed a strong **`Novel`**:**`Injection`** interaction effect (*P* = 2.22e-6, coefficient estimate *z*-test, *n* = 32 mice) and a weak effect of **`Novel`** alone (*P* = 0.025), but no effect of **`Sex`** (*P* = 0.137) or **`Injection`** (*P* = 0.574) alone or for any other effects. Using the coefficients from this GLMM, we then used the marginaleffects R package (https://github.com/vincentarelbundock/marginaleffects) to calculate the marginal effect of **`Novel`** condition (Novel – Familiar) on each **`Day`** independently for each **`Injection`** group. We used the marginal effect estimates and standard errors to calculate a *P-*value for each **`Injection`**/**`Day`** combination with a *z*-test, and then corrected for multiple comparisons within each **`Injection`** group using the Hochberg-Bonferroni step-up procedure^120^.

For subsequent experiments (Fig. 3d–f; Extended Data Fig. 2b), we performed a single retrieval test per animal and tested for significant differences across groups using Wilcoxon rank-sum tests.

### Histology

We visualized mCherry, mScarlet, and YFP signals to validate transgene expression in our LS chemogenetics (Extended Data Fig. 2a,b) and CGRP neuron optogenetics (Fig. 3d–f) experiments. Mice were deeply anesthetized (2 mg/kg Euthasol ip.) and then transcardially perfused with phosphate buffered saline (PBS) followed by 4% paraformaldehyde (PFA) in PBS. Brains were then extracted and post-fixed overnight in 4% PFA at 4°C, and then cryo-protected overnight in 30% sucrose in PBS at 4°C. Free-floating sections (40 μm) were prepared with a cryostat, mounted with DAPI Fluoromount-G (Southern Biotech 0100), and imaged with a slide scanner (Hamamatsu NanoZoomer S60).

To visualize ExRai-AKAR2 signal (Fig. 6c), we stained for GFP immunoreactivity in the CEA. To validate CGRP neuron ablation following taCasp3-TEVp injection (Fig. 4p; Extended Data Fig. 9f), we stained for CGRP immunoreactivity in the PB. Briefly, sections were washed, blocked (3% NDS and 0.3% Triton-X in PBS for 90 min), and then incubated with primary antibody (rabbit anti-GFP, Novus NB600-308, 1:1,000; mouse anti-CGRP, Abcam ab81887, 1:250) in blocking buffer overnight at 4 °C. Sections were then washed, incubated with secondary antibody (Alexa Fluor 647 donkey anti-rabbit, Invitrogen A31573, 1:500; Alexa Fluor 568 donkey anti-mouse, Life Technologies A10037, 1:500) in blocking buffer for 90-min at room temperature, washed again, mounted with DAPI Fluoromount-G (Southern Biotech), and imaged with a slide scanner (Hamamatsu NanoZoomer S60).

### Modified Allen CCF atlas

The reference atlas we used is based on the 25-μm resolution Allen CCFv3^43^ (https://atlas.brain-map.org). For Fos experiments, we considered every brain region in the atlas that met the following criteria: (1) total volume ≥0.1 mm^3^; (2) lowest level of its branch of the ontology tree (cortical layers/zones not included). We made two modifications to the standard atlas for this study.

First, we reassigned brain region IDs in order to increase the clarity of our Fos visualizations incorporating the atlas and to accurately represent the full functional extent of the CEA. We merged several small region subdivisions together into the larger LG, PVH, PV, MRN, PRN, and SPV regions (see Extended Data Table 1 for list of brain region abbreviations). We reassigned all cortical layers and zones to their immediate parent regions (for example, *Gustatory areas, layers 1–6b* (111–117) were reassigned to *Gustatory areas* (110)). We merged all unassigned regions (*-un* suffix in Allen CCF) into relevant parent regions (for example, *HPF-un* (563) was reassigned to *Hippocampal formation* (462)). We reassigned the voxels immediately surrounding the CEA that were assigned to *Striatum* (581) to *Central amygdalar nucleus* (605), because we found that cells localized to these CEA-adjacent voxels had very similar Fos and Neuropixels responses compared to cells localized strictly within CEA. These atlas changes were used throughout the paper (Fos experiments and Neuropixels experiments). Summaries across the entire CEA (for example, Fig. 2g, Fig. 3h, and Figs. 4,5) included all atlas voxels assigned to the parent CEA region (605) and the CEAc (606), CEAl (607), and CEAc (608) subdivisions.

Second, we made the left and right hemispheres symmetric in order to facilitate pooling data from both hemispheres for our Fos visualizations. To ensure that the hemispheres of the Allen CCF were perfectly symmetric, we replaced the left hemisphere with a mirrored version of the right hemisphere. This atlas change was used only for Fos experiments.

### Brainwide Fos timepoints

All mice whose data is shown in Figs. 1–3 were trained in the one-reward CTA paradigm as described above. Consumption timepoint (Fig. 1): Mice were euthanized 60-min after the end of the consumption period on conditioning day (no LiCl injection was given). Malaise timepoint (Fig. 2): Mice were euthanized 60-min after the LiCl injection on conditioning day. Retrieval timepoint (Fig. 2): Mice received the LiCl conditioning described above, and then were returned to the operant box two days later for another consumption of the paired flavor using the same task structure as above. Mice were euthanized 60-min after the end of the consumption period of the retrieval session (no LiCl was given during retrieval session). CGRP stim timepoint (Fig. 3): Mice were euthanized 60-min after the onset of CGRP neuron stimulation on conditioning day, and stimulation continued for the full 60-min. LS activation timepoint (Extended Data Fig. 2): Mice received an ip. injection of 3 mg/kg CNO 45-min before consumption and were then euthanized 60-min after the LiCl injection on conditioning day.

Mice were deeply anesthetized (2 mg/kg Euthasol ip.) and then transcardially perfused with ice-cold PBS + heparin (20 U/ml; Sigma H3149) followed by ice-cold 4% PFA in PBS. Brains were then extracted and post-fixed overnight in 4% PFA at 4°C.

### Tissue clearing and immunolabeling

Brains were cleared and immunolabeled using an iDISCO+ protocol as previously described^41,42,121^. All incubations were performed at room temperature unless otherwise noted.

Clearing. Brains were serially dehydrated in increasing concentrations of methanol (Carolina Biological Supply 874195; 20%, 40%, 60%, 80%, 100% in doubly distilled water (ddH_2_O); 45 min–1 h each), bleached in 5% hydrogen peroxide (Sigma H1009) in methanol overnight, and then serially rehydrated in decreasing concentrations of methanol (100%, 80%, 60%, 40%, 20% in ddH_2_O; 45 min–1 h each).

Immunolabeling. Brains were washed in 0.2% Triton X-100 (Sigma T8787) in PBS, followed by 20% DMSO (Fisher Scientific D128) + 0.3 M glycine (Sigma 410225) + 0.2% Triton X-100 in PBS at 37°C for 2 d. Brains were then washed in 10% DMSO + 6% normal donkey serum (NDS; EMD Millipore S30) + 0.2% Triton X-100 in PBS at 37°C for 2–3 d to block non-specific antibody binding. Brains were then twice washed for 1 h at 37°C in 0.2% Tween-20 (Sigma P9416) + 10 mg/ml heparin in PBS (PTwH solution) followed by incubation with primary antibody solution (rabbit anti-Fos, 1:1000; Synaptic Systems 226008) in 5% DMSO + 3% NDS + PTwH at 37°C for 7 d. Brains were then washed in PTwH 6× for increasing durations (10 min, 15 min, 30 min, 1 h, 2 h, overnight) followed by incubation with secondary antibody solution (Alexa Fluor 647 donkey anti-rabbit, 1:200; Abcam ab150075) in 3% NDS + PTwH at 37°C for 7 d. Brains were then washed in PTwH 6× for increasing durations again (10 min, 15 min, 30 min, 1 h, 2 h, overnight).

CGRP stim timepoint samples also received primary (chicken anti-GFP, 1:500; Aves GFP-1020) and secondary (Alexa Fluor 594 donkey anti-chicken, 1:500; Jackson Immuno 703-585-155) antibodies for ChR2-YFP immunolabeling during the above protocol.

Final storage and imaging: Brains were serially dehydrated in increasing concentrations of methanol (20%, 40%, 60%, 80%, 100% in ddH_2_O; 45 min–1 h each), then incubated in a 2:1 solution of dichloromethane (DCM; Sigma 270997) and methanol for 3 h followed by 2× 15-min washes 100% DCM. Before imaging, brains were stored in the refractive index-matching solution dibenzyl ether (DBE; Sigma 108014).

### Fos light sheet imaging

Cleared and immunolabeled brains were glued (Loctite 234796) ventral side-down to a 3D-printed holder and imaged in DBE using a dynamic axial sweeping light sheet fluorescence microscope^122^ (Life Canvas SmartSPIM). Images were acquired using a 3.6×/0.2 NA objective with a 3,650 µm × 3,650 µm field-of-view onto a 2,048 px × 2,048 px sCMOS camera (pixel size: 1.78 µm × 1.78 µm) with a spacing of 2 µm between horizontal planes (nominal z-dimension point spread function: 3.2–4.0 µm). Imaging the entire brain required 4 × 6 tiling across the horizontal plane and 3,300–3,900 total horizontal planes. Autofluorescence channel images were acquired using 488-nm excitation light at 20% power (maximum output: 150 mW) and 2-ms exposure time, and Fos channel images were acquired using 639-nm excitation light at 90% power (maximum output: 160 mW) and 2-ms exposure time. For CGRP stim timepoint samples, a bilateral volume encompassing both PB was imaged separately using 561-nm excitation light at 20% power (maximum output: 150 mW) and 2-ms exposure time to confirm ChR2-YFP expression.

After acquisition, tiled images for the Fos channel were first stitched into a single imaging volume using the TeraStitcher C++ package^123^ (https://github.com/abria/TeraStitcher). These stitching parameters were then directly applied to the tiled autofluorescence channel images, yielding two aligned 3D imaging volumes with the same final dimensions. After tile stitching, striping artifacts were removed from each channel using the Pystripe Python package^124^ (https://github.com/chunglabmit/pystripe).

We registered the final Fos imaging volume to the Allen CCF using the autofluorescence imaging volume as an intermediary^41^. We first downsampled both imaging volumes by a factor of 5 for computational efficiency. Autofluorescence→atlas alignment was done by applying an affine transformation to obtain general alignment using only translation, rotation, shearing, and scaling, followed by applying a b-spline transformation to account for local nonlinear variability among individual brains. Fos→autofluorescence alignment was done by applying only affine transformations to account for brain movement during imaging and wavelength-dependent aberrations. Alignment transformations were computed using the Elastix C++ package^125,126^ (https://github.com/SuperElastix/elastix). These transformations allowed us to transform Fos^+^ cell coordinates first from their native space to the autofluorescence space and then to Allen CCF space.

### Deep learning-assisted cell detection pipeline

We first use standard machine vision approaches to identify candidate Fos^+^ cells based on peak intensity and then use a convolutional neural network to remove artifacts. Our pipeline builds upon the ClearMap Python package^42,127^ (https://github.com/ChristophKirst/ClearMap2) for identifying candidate cells and the Cellfinder Python package^128^ (https://github.com/brainglobe/cellfinder) for artifact removal.

Cell detection. ClearMap operates through a series of simple image processing steps. First, the Fos imaging volume is background-subtracted using a morphological opening (disk size: 21 px). Second, potential cell centers are found as local maxima in the background-subtracted imaging volume (structural element shape: 11 px). Third, cell size is determined for each potential cell center using a watershed algorithm (see below for details on watershed detection threshold). Fourth, a final list of candidate cells is generated by removing all potential cells that are smaller than a preset size (size threshold: 350 px). We confirmed that our findings were consistent across a wide range of potential size thresholds.

We implemented three changes to the standard ClearMap algorithm. First, we de-noised the Fos imaging volume using a median filter^129^ (function: *scipy.ndimage.median_filter*; size: 3 px) before the background subtraction step. Second, we dynamically adjusted the watershed detection threshold for each sample based on its fluorescence intensity. This step was important for achieving consistent cell detection performance despite changes in background and signal intensity across batches and samples due to technical variation in clearing, immunolabeling, and imaging. Briefly, we selected a 1,000 px × 1,000 px × 200 px subvolume at the center of each sample’s Fos imaging volume. We then median filtered and background subtracted this subvolume as described above. We then used sigma clipping^130^ (function: *astropy.stats.sigma_clipped_stats*; *sigma*=3.0, *maxiters*=10, *cenfunc*=’median’, *stdfunc*=’mad_std’) to estimate the mean background (non-cell) signal level for this subvolume, µ_bg_, and set each sample’s watershed detection threshold to 10*µ_bg_. Third, we removed from further analysis all cell candidates that were located outside the brain, in the anterior olfactory areas or cerebellum (which were often damaged during dissection), or in the ventricles, fiber tracts, and grooves following registration to the Allen CCF.

Cell classification. One limitation of the watershed algorithm implemented by ClearMap is that it identifies any high-contrast feature as a candidate cell, including exterior and ventricle brain edges, tissue tears, bubbles, and other aberrations. To overcome this limitation, we re-trained the 50-layer ResNet^131^ implemented in Keras (https://keras.io) for TensorFlow (https://www.tensorflow.org) from the Cellfinder^128^ Python package to classify candidate Fos^+^ cells in our high-resolution light sheet imaging dataset as true Fos^+^ cells or artifacts. This network uses both the autofluorescence and Fos channels during classification because the autofluorescence channel has significant information about high-contrast anatomical features and imaging aberrations. We first manually annotated 2,000 true Fos^+^ cells and 1,000 artifacts from each of four brains across two technical batches using the Cellfinder Napari plugin, for a total training dataset of 12,000 examples. We then re-trained the Cellfinder network (which had already been trained on ∼100,000 examples from serial two-photon images of GFP-labeled neurons) using TensorFlow over 100 epochs with a learning rate of 0.0001 and 1,200 examples (10% of the training dataset) held out for validation. Re-training took 4 d 16 min 41 s on a high performance computing cluster using 1 GPU and 12 CPU threads. We achieved a final validation accuracy of 98.33%. Across all samples in our brainwide Fos dataset, our trained convolutional neural network removed 15.99% ± 0.58% (mean ± s.e.m.; range: 2.96% – 32.71%; *n* = 99 brains across all timepoints) of cell candidates from ClearMap as artifacts.

Atlas registration. We used the ClearMap interface with Elastix to transform the coordinates of each true Fos^+^ cell to Allen CCF space using the transformations described above. We then used these coordinates to assign each Fos^+^ cell to an Allen CCF brain region. For each sample, we generated a final data structure containing the Allen CCF coordinates (*x*,*y*,*z*), size, and brain region for each true Fos^+^ cell.

### Fos density maps

We generated 3D maps of Fos^+^ cell density by applying a gaussian kernel-density estimate (KDE)^129^ (function: *scipy.stats.gaussian_kde*) in Python to all Fos^+^ cells across all animals within a given experimental condition (for example, Novel flavor + Consumption timepoint). These maps are visualized in Fig. 1e,f, Fig.2f, Fig. 3k, Extended Data Fig. 1a–f, Extended Data Fig. 2g, and Extended Data Fig. 6e,f.

We first generated a table containing the Allen CCF coordinates (*x*,*y*,*z*) for every Fos^+^ cell in every animal within an experimental condition. At this stage, we listed each cell twice (once with its original coordinates and once with its ML (*z*) coordinate flipped to the opposite hemisphere) in order to pool data from both hemispheres. We then assigned each cell a weight equal to the inverse of the total number of Fos^+^ cells in that animal to ensure that each animal within an experimental condition would be weighted equally. We then fit a 3D gaussian KDE for each experimental condition using the *scipy.stats.gaussian_kde* function, and manually set the kernel bandwidth for every experimental condition to be equal at 0.04. We then evaluated this KDE at every voxel in the Allen CCF (excluding voxels outside the brain or in anterior olfactory areas, cerebellum, ventricles, fiber tracts, grooves) to obtain a 3D map of Fos^+^ density for each condition. Lastly, we normalized the KDE for each experimental condition by dividing by its sum as well as the voxel size of the atlas, (0.025 mm)^3^, to generate a final 3D map with units of “% Fos^+^ cells per mm^3^”. For the CGRP stim timepoint, we assigned each cell a weight equal to the inverse of the number of Fos^+^ cells in the PB of that animal, rather than the total number Fos^+^ cells, to account for variations in ChR2-induced activation across mice and flavor conditions.

To examine the difference in Fos^+^ cell density across flavor conditions (for example, in Fig. 1e,f and Extended Data Fig. 1b for the Consumption timepoint) we simply subtracted the 3D KDE volumes for the two conditions, Novel – Familiar, and then plotted coronal sections through this subtracted volume with Allen CCF boundaries overlaid. The colorbar limits for all Novel – Familiar ΔFos KDE figures are ±0.5% Fos^+^ cells per mm^3^ and for all Average Fos KDE figures are 0–1% Fos^+^ cells per mm^3^.

We used the WebGL-based Neuroglancer tool (https://github.com/google/neuroglancer) to generate interactive 3D visualizations of the Fos^+^ cell density maps for each experimental timepoint (https://www.brainsharer.org/ng/?id=872). To achieve this, we used the cloudvolume Python package (https://github.com/seung-lab/cloud-volume) to convert our 3D KDE volumes from numpy format to precomputed layers compatible with Neuroglancer and then loaded these layers into the Brainsharer web portal (https://brainsharer.org) to create the final visualization.

### Fos GLMMs

We adopted a GLMM to analyze the brainwide Fos data in Figs. 1–3. This allowed us to model the contribution of flavor and experimental timepoint to neural activation in each brain region, while also accounting for (1) the overdispersed, discrete nature of the data by employing a negative binomial link function, (2) the contribution of batch-to-batch technical variation in tissue clearing, immunolabeling, and imaging by modeling this as a random effect, and (3) the potential contribution of sex as a biological variable by modeling this as a fixed effect.

The first step was to determine if there was any effect of novel/familiar, experimental timepoint, or their interaction for each brain region, while controlling the false discovery rate (FDR) across all regions. To accomplish this, we fit a full GLMM for each brain region using the glmmTMB R package^119^ (https://github.com/glmmTMB/glmmTMB) with a negative binomial link function (*nbinom2*) and the formula:

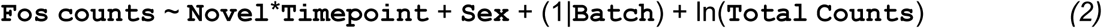

where **`Fos counts`** is the number of Fos^+^ cells in a brain region, **`Novel`** (Novel, Familiar), **`Timepoint`** (Consumption, Malaise, Retrieval), and **`Sex`** (Female, Male) are fixed effect categorical variables, (1|**`Batch`**) is a random effect for each technical batch (that is, each set of samples that underwent tissue clearing, immunolabeling, and light sheet imaging together), ln(**`Total Counts`**) is an offset term for the total number of Fos^+^ cells in each sample, and * represents all possible main effects and interactions (Fig. 2b). We then fit a reduced GLMM for each brain region, which was the same as the full model *(2)* but with the **`Novel`**\***`Timepoint`** terms (that is, all main effects and interactions related to flavor novelty and experimental timepoint) removed. We compared these two models for each brain region using likelihood-ratio χ^2^-tests and then adjusted the resultant *P*-values using the Benjamini-Krieger-Yekutieli two-step procedure^132^ to permit a 10% FDR across all brain regions. The 10% FDR threshold used here is standard for brainwide Fos studies^133–136^. Of 200 brain regions tested, 130 met this criterion and were included for downstream analysis.

We next specifically tested the effect of flavor novelty on Fos counts separately at each experimental timepoint for the 130 brain regions that passed that FDR threshold above. To calculate the marginal effect of **`Novel`** condition (Novel – Familiar) at each **`Timepoint`** for each brain region, we used the marginaleffects R package (https://github.com/vincentarelbundock/marginaleffects) to do post-hoc testing of the full GLMM. We used the marginal effect estimates and standard errors to calculate a *P-*value for each **`Timepoint`** with a *z*-test, and then corrected for multiple comparisons across timepoints within each brain region using the Hochberg-Bonferroni procedure^120^ We also used the ratio of these marginal effect estimates and standard errors to compute the standardized average difference in Fos^+^ cell counts across **`Novel`** conditions for each brain region at each timepoint (*Z* = estimate/standard error; Fig. 2c–e; Extended Data Table 1). The advantage of this metric is that it explicitly accounts for variance within and across groups, for effects of sex and technical batch, and is independent of brain region size.

When displaying Fos^+^ cell counts for individual samples (Fig. 1e,f; Fig. 2g; Extended Data Fig. 6b), we divided the number of Fos^+^ cells for each animal/brain region by the total number of Fos^+^ cells in that animal and by the Allen CCF volume of that brain region, so that each region’s data are presented as “% Fos^+^ cells per mm^3^”. We used the *P*-values from the GLMM marginal effect *z*-tests described above to assess significance.

To examine the brainwide shift in Novel – Familiar coding across timepoints (Fig. 2c; Extended Data Fig. 3a–c), we used the Violinplot MATLAB package (https://github.com/bastibe/Violinplot-Matlab) to plot the distribution of standardized average difference *Z* values at each timepoint for the brain regions that passed our FDR threshold and then used Kolmogorov-Smirnov tests to assess whether these distributions were significantly different from each other, correcting for multiple comparisons across timepoints using the Hochberg-Bonferroni procedure^120^.

To identify structure in Novel – Familiar coding across timepoints (Fig. 2d; Extended Data Fig. 3d–n), we used the built-in MATLAB *linkage* function (*method*=’ward’, *metric*=’chebychev’) to create a hierarchical tree using the standardized average difference *Z* values at each timepoint for the brain regions that passed our FDR threshold. The input matrix was 130 brain regions × 3 timepoints. We then used the built-in MATLAB *dendrogram* function to plot this hierarchical tree and used a distance threshold of 4.7 for clustering.

We followed an analogous procedure to analyze brainwide Fos data for the CGRP stim timepoint (Fig. 3; Extended Data Fig. 6). To account for variations in ChR2 expression across mice and flavor conditions, we weighted Fos^+^ cell counts by the number of Fos*^+^* cells in the PB in these analyses. Specifically, to compare the effects of CGRP neuron stimulation and LiCl-induced malaise on overall Fos levels (Extended Data Fig. 6c), we fit a GLMM for each brain region with the negative binomial link function and the formula:

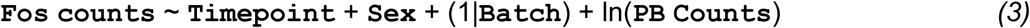

where **`Fos counts`** is the number of Fos^+^ cells in a brain region, **`Timepoint`** (Consumption, Malaise, Retrieval, CGRP stim), and **`Sex`** (Female, Male) are fixed effect categorical variables, (1|**`Batch`**) is a random effect for each technical batch, ln(**`PB Counts`**) is an offset term for the total number of Fos^+^ cells in the PB of each sample, and * represents all possible main effects and interactions. For this model, we did not include any terms related to flavor novelty because we were specifically investigating changes in overall Fos levels. We then plotted the coefficient estimate *Z*-values from this GLMM (Extended Data Fig. 6c). To compare the effects of CGRP neuron stimulation and LiCl-induced malaise on Fos levels in the novel versus familiar flavor condition (Extended Data Fig. 6d), we fit a GLMM for each brain region with the formula:

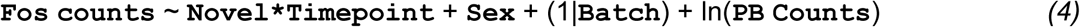

where **`Fos counts`** is the number of Fos^+^ cells in a brain region, **`Novel`** (Novel, Familiar), **`Timepoint`** (Malaise, CGRP stim), and **`Sex`** (Female, Male) are fixed effect categorical variables, (1|**`Batch`**) is a random effect for each technical batch, ln(**`PB Counts`**) is an offset term for the total number of Fos^+^ cells in the PB of each sample, and * represents all possible main effects and interactions. For this model, we only included the experimental timepoints in which parabrachial CGRP neurons were activated either optogenetically (CGRP stim) or pharmacologically (Malaise); see Extended Data Fig. 6b for quantification of PB activation. We then calculated and plotted the marginal effect of **`Novel`** condition (Novel – Familiar) separately for each timepoint and brain region (Extended Data Fig. 6d). We also used the GLMM from Equation 4 to calculate the *P*-value for Fig. 3h. When displaying Fos^+^ cell counts for individual animals (Fig. 3h, CGRP stim timepoint) or brain regions (Fig. 3i,j; Malaise and CGRP stim timepoints), we first divided the number of Fos^+^ cells for each brain region in each animal by the number of Fos^+^ cells in the PB for that animal and by the Allen CCF volume of that brain region. We then divided this number by the average ratio of total Fos^+^ cells to PB Fos^+^ cells across every sample in that timepoint (Malaise or CGRP stim), which yielded a final measure of each animal/region’s Fos^+^ cells as a percentage of the entire brain’s Fos^+^ cells weighted by the relative count of PB Fos^+^ cells for that animal. We obtained consistent results by instead subsampling the animals in the CGRP stim timepoint to have approximately equal Fos^+^ cell counts in both flavor conditions and then weighting by total Fos^+^ cell count of each animal.

### Fos correlation analysis

To quantify Fos correlations across individual mice (Extended Data Fig. 4), we considered each experimental timepoint (Consumption, Malaise, Retrieval) separately. We first assembled the relative Fos^+^ cell counts (% per mm^3^) for every brain region that passed our FDR threshold and then sorted these regions using the hierarchical tree fit above, which resulted in a 130 brain regions × 24 animals input matrix for each experimental timepoint. We then used the built-in MATLAB *corr* function to calculate and visualize pairwise correlations among all brain regions (Extended Data Fig. 4a). To estimate the correlation among individual brain regions in the amygdala cluster at each timepoint (Extended Data Fig. 4b), we averaged each brain region’s pairwise correlations with all other amygdala cluster regions in the correlation matrices above. We tested whether the correlation among individual amygdala cluster brain regions was significant at each timepoint using Wilcoxon signed-rank tests, correcting for multiple comparisons across timepoints using the Hochberg-Bonferroni procedure^120^. To estimate the correlation between the amygdala cluster and every other cluster at each timepoint (Extended Data Fig. 4c), we averaged the pairwise correlations for all brain region pairs across the two clusters.

### LS activation Fos analysis

To compare the effects of LS activation on Fos levels in the LS and CEA (Extended Data Fig. 2d,e), we calculated *P*-values for these two regions using a GLMM with the formula:

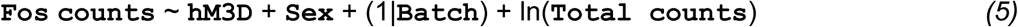

where **`Fos counts`** is the number of Fos^+^ cells in a brain region, **`hM3D`** (hM3D, YFP), and **`Sex`** (Female, Male) are fixed effect categorical variables, (1|**`Batch`**) is a random effect for each technical batch, ln(**`Total Counts`**) is an offset term for the total number of Fos^+^ cells in each sample, and * represents all possible main effects and interactions. The *P*-values in the figure are from the **`hM3D`** coefficient estimates.

To compare the effects of LS activation of Fos levels across the brain (Extended Data Fig. 2f), we plotted the average Fos level (% Fos^+^ cells per mm^3^) across all mice in each condition (hM3D, YFP) separately for three groups of brain regions: the amygdala network, the septal complex, and other regions. We then used the built-in MATLAB *aoctool* function to fit a one-way analysis of covariance model for these three groups of brain regions, correcting for multiple comparisons using the Bonferroni procedure.

### RNAscope fluorescence in situ hybridization

We sliced 18–25-µm thick tissue sections from perfused tissue. Multiplex FISH (Fig. 3l–n; Extended Data Fig. 6g,h) was performed using the RNAscope Multiplex Fluorescent Assay v2 (ACD, no. 323120) with the following probes: Mm-Calcrl (452281), Mm-Sst-C2 (404631-C2, 1:50 dilution in C1 solution), Mm-Prkcd-C3 (441791-C3, 1:50 dilution in C1 solution), and Mm-Fos-C4 (316921-C4, 1:50 dilution in C1 solution). The *Calcrl*, *Sst*, *Prkcd*, and *Fos* probes were linked to Opal 690, Opal 520, Opal 620, and Opal 570 fluorophores, respectively (Akoya Biosciences). All fluorophores were reconstituted in dimethylsulfoxide according to Akoya Biosciences instructions and diluted 1:1,200 in tyramide signal amplification (TSA) buffer included in the RNAscope kit. After in situ hybridization, slides were coverslipped using Fluoromount-G containing DAPI (Southern Biotech).

We obtained 20× z-stacks from the CEA using a confocal microscope (Leica TCS SP8 X) and converted these into maximum intensity projections for each labeled RNA. We trained a Cellpose^137,138^ (https://cellpose.readthedocs.io) model to identify *Fos*^+^ cells in the maximum intensity projections using eight manually corrected examples, and then used this model to identify *Fos*^+^ cells in the remaining images. We manually annotated whether every *Fos*^+^ cell identified by Cellpose also expressed *Sst*, *Prkcd*, and/or *Calcrl*. We used the full *z*-stacks for each labeled RNA for this process in order to ensure that potentially overlapping cells were labeled separately. We then imaged each section with a slide scanner (Hamamatsu Nanozoomer S60), registered them to the Allen CCF using ABBA (https://abba-documentation.readthedocs.io). To remove *Fos*^+^ cells outside the CEA from analysis, we manually aligned the confocal and nanozoomer images using the *Fos* channel in each image as a guide and then manually transferred the CEA boundaries to the confocal images.

### Slice electrophysiology

All electrophysiology experiments (Fig. 3c) were performed on brain slices collected at approximately the same time of day. *Calca::Cre* mice were first injected with 400 nl of AAV5-EF1a-DIO-hChR2(H134R)-eYFP (titer: 1.2e13 GC/ml; manufacturer: PNI Viral Core Facility) bilaterally into the PB 6+ weeks before the experiment. On the day of the experiment, mice were anesthetized with isoflurane and decapitated to remove the brain. After extraction, the brain was immersed in ice-cold NMDG ACSF (92 mM NMDG, 2.5 mM KCl, 1.25 mM NaH_2_PO_4_, 30 mM NaHCO_3_, 20 mM HEPES, 25 mM glucose, 2 mM thiourea, 5 mM Na-ascorbate, 3 mM Na-pyruvate, 0.5 mM CaCl_2_·4H_2_O, 10 mM MgSO_4_·7H_2_O, and 12 mM *N*-Acetyl-L-cysteine; pH adjusted to 7.3–7.4) for 2 min. Afterward, coronal slices (300 μm) were sectioned using a vibratome (VT1200s, Leica, Germany) and then incubated in NMDG ACSF at 34 °C for approximately 15 min. Slices were then transferred into a holding solution of HEPES ACSF (92 mM NaCl, 2.5 mM KCl, 1.25 mM NaH_2_PO_4_, 30 mM NaHCO_3_, 20 mM HEPES, 25 mM glucose, 2 mM thiourea, 5 mM Na-ascorbate, 3 mM Na-pyruvate, 2 mM CaCl_2_·4H_2_O, 2 mM MgSO_4_·7H_2_O and 12 mM *N*-Acetyl-L-cysteine, bubbled at room temperature with 95% O_2_/5% CO_2_) for at least 60 min until recordings were performed.

Whole-cell recordings were performed using a Multiclamp 700B (Molecular Devices) using pipettes with a resistance of 4–7 MΩ filled with an internal solution containing 120 mM potassium gluconate, 0.2 mM EGTA, 10 mM HEPES, 5 mM NaCl, 1 mM MgCl_2_, 2 mM Mg-ATP and 0.3 mM NA-GTP, with the pH adjusted to 7.2 with KOH and the osmolarity adjusted to around 289 mmol kg^−1^ with sucrose. During recordings, slices were perfused with a recording ACSF solution (100 μM picrotoxin, 120 mM NaCl, 3.5 mM KCl, 1.25 mM NaH_2_PO_4_, 26 mM NaHCO_3_, 1.3 mM MgCl_2_, 2 mM CaCl_2_, and 11 mM D-(+)-glucose) with 1 µM TTX and 100 µM 4AP that was continuously bubbled with 95% O_2_/5% CO_2_. Infrared differential interference contrast-enhanced visual guidance was used to select neurons that were 3–4 cell layers below the surface of the slices. All CEAc/l recordings were made where eYFP-expressing CGRP axons were visible, and all CEAm recordings were made more medial to this location using the Allen CCF as a guide The recording solution was delivered to slices via superfusion driven by peristaltic pump (flow rate of 4–5 ml/min) and was held at room temperature. The neurons were held at −70 mV (voltage clamp), and the pipette series resistance was monitored throughout the experiments by hyperpolarizing steps of 1 mV with each sweep. If the series resistance changed by >20% during the recording, the data were discarded. Whole-cell currents were filtered at 4 kHz online and digitized and stored at 10 KHz (Clampex 10; MDS Analytical Technologies). Currents were then filtered at 1 kHz offline prior to analysis. During the experiment, we measured light-evoked oEPSCs every 30 s with light stimulation (0.074 mW/mm^2^) delivered for a duration of 5 ms. Twenty repetitions of the stimulation protocol were recorded per cell after stable evoked EPSCs were achieved. All experiments were completed within 4 h after slices were made to maximize cell viability and consistency.

### Two-reward CTA paradigm

For Neuropixels recording experiments (Figs. 4–6), we used a two-reward CTA paradigm (rather than the one-reward paradigm used for Fos experiments). This allowed us to compare neural correlates of the flavored novel reward, which was delivered from one port, with responses related to a control port that delivered water. Experiments were performed in an operant box situated in a sound-attenuating chamber and equipped with a single speaker and with two custom 3D-printed nosepoke ports with built-in lights and infrared beam breaks within each port. The nosepoke ports each contained a reward delivery tube that was calibrated to deliver 20-µl rewards via a solenoid valve. The ports were located on either side of the same wall of the operant box.

Basic task structure and training. Mice first underwent a basic shaping procedure to train them to drink from two reward ports at a relatively equal rate in a cued manner (Extended Data Fig. 8a). Each behavioral session had the following structure: First, the mouse was allowed to acclimate to the chamber for 5 min. Then, the consumption period began and rewards were made available in a trial-based manner that forced mice to drink from the two ports at a relatively equal rate throughout the session. At the beginning of each trial, one port was randomly selected and made available to the mouse. This was cued via the port light turning on and a distinct tone (2.5 kHz or 7.5 kHz; 70 dB) playing. The mouse had 10 s to enter the port and receive a reward, which was detected by the infrared beam break. The end of the 10-s reward availability period or entering either port ended the trial; at this point, the cueing light and sound were terminated and an inter-trial interval was initiated (randomly selected from a uniform distribution of 10–20 s; 1-s step size). At the end of the inter-trial interval, a new trial would begin as long as the mouse had not entered either reward port in the previous 2 s; otherwise, the next trial was delayed until this criterion was satisfied. To ensure that mice drank from the two ports at a relatively equal rate, we required that each consecutive block of 10 successful rewarded trials must be evenly split between the two ports. The consumption period ended when 1.2 ml (60 rewards) was consumed. Mice learned to perform this task nearly perfectly (<5 unsuccessful trials per session) in approximately one week. During initial training, both ports delivered water. After mice were trained in the task, they underwent the chronic Neuropixels surgery described above and were allowed to recover for at least five days. Mice were then returned to daily training, with the addition of a delay period following the consumption period. At the beginning of the delay period (immediately after the final reward), mice were transferred to a distinct second context, which was triangular in shape with smooth white acrylic walls. Mice remained in this second context for at least 30-min before returning to the homecage. Mice were acclimated to the delay period and second context, and to tethering of the Neuropixels probe and optical fiber, for at least four days before proceeding to conditioning experiments. Variations to this basic task structure for specific experiments that were used following training and surgery are described below.

CGRP neuron stimulation conditioning experiment (Figs. 4,5). On conditioning day, the same behavioral session structure was followed, but now one port delivered water and the other port delivered sweetened grape Kool-Aid (0.06% grape + 0.3% saccharin sodium salt). The novel flavor port was counterbalanced across mice. Mice (*n* = 8) were run in two separate cohorts separated by approximately two months. After the 30-min delay period in the second context, the CGRP neuron stimulation period began and lasted for 45-min in the same second context. Blue light was generated using a 447-nm laser and delivered to the animal using a ⌀200-µm patch cable. Light power was calibrated to approximately 10 mW at the patch cable tip. The laser was controlled with a Pulse Pal signal generator programmed to deliver 5-ms laser pulses at 10 Hz. For the duration of the CGRP neuron stimulation period, the laser was pulsed for 3-s bouts and then off for random intervals chosen from an exponential distribution (minimum: 1 s; mean: 3 s; maximum: 7.8 s). Following the 45-min neuron stimulation period, mice were returned to the homecage overnight. The following day, mice underwent a forced retrieval session that followed the same trial structure as previous sessions, and the flavor and water were delivered from the same ports as on conditioning day.

CGRP^CEA^ projection stimulation conditioning experiment (Figs. 4–5). This experiment was performed using the same strategy as the cell-body stimulation experiment described above in a separate cohort of mice (*n* = 8) with the following changes. Green light was generated using a 532-nm laser and calibrated to approximately 3 mW at the patch cable tip. To minimize potential photoelectric artifacts in our recordings, we (1) positioned the tip of the optical fiber 1.5 mm from the Neuropixels shanks in CEA for an irradiance at the electrodes of approximately 0.1 mW/mm^2^ and (2) reduced the laser pulse width to 2 ms. These stimulation parameters are sufficient to activate the ultra-sensitive opsin ChRmine^139^.

LiCl conditioning experiment (Fig. 4m–r; Extended Data Fig. 9f,g). This experiment was performed in a separate cohort of mice (*n* = 4 control mice and 4 CGRP neuron ablation mice). It followed the same structure as above, except that LiCl (125 mg/kg ip.) was injected to induce visceral malaise after 30-min in the second context (delay period) instead of CGRP neuron stimulation. For behavioral validation of CGRP neuron ablation (Extended Data Fig. 9g), we included five mice that were not used for recordings but either received taCasp3 virus (*n* = 2 ablation mice) or did not undergo surgery (*n* = 3 control mice).

Familiarization experiment (Fig. 5f; Extended Data Fig. 10f–k). A different flavor, sweetened cherry Kool-Aid (0.06% cherry + 0.3% saccharin sodium salt), was used for this experiment. The experiment followed the same basic task structure as during initial training for the two-reward CTA paradigm, without any aversive conditioning experiences (LiCl injection or CGRP neuron stimulation). The experiment was run on three consecutive days. On the first day (Novel day), one port contained the novel sweetened cherry Kool-Aid flavor and the other port contained water. On the second day, the port locations were switched. On the third day (Familiar day), the port locations were switched again (that is, back to the initial locations from Novel day).

CGRP neuron stimulation→LiCl injection experiment (Extended Data Fig. 9h–j). This experiment did not involve rewards or use the task structure described above. First, mice were allowed to acclimate to the operant box recording chamber for 5 min. Then, CGRP neurons were photostimulated using the same protocol as for the acute Neuropixels recording experiment described below. Briefly, mice received 1 s of 10-Hz CGRP neuron stimulation followed by a 9-s inter-trial interval for a total of 10 min/60 trains. After a 5-min recovery period, LiCl (125 mg/kg ip.) was then injected to induce visceral malaise. Mice remained in the recording chamber for at least 15 min before being returned to the homecage.

We used 27 mice for chronic Neuropixels recording experiments (Extended Data Fig. 7c). Animals 1–4 were used for the CGRP neuron stimulation conditioning experiment. Animals 5–8 were used for multiple experiments with the following timeline: (1) CGRP neuron stimulation conditioning experiment, (2) familiarization experiment, (3) CGRP neuron stimulation→LiCl injection experiment. Animals 9–12 were control mice used for the LiCl conditioning experiment. Animals 13–16 were CGRP neuron ablation mice used for the LiCl conditioning experiment. Animals 17–24 were used for the CGRP^CEA^ projection stimulation conditioning experiment. Animals 25–27 were used for the familiarization experiment.

### Chronic Neuropixels recordings

Before beginning experiments, we performed a series of test recordings for each mouse in order to identify the recording sites along each Neuropixels shank that were located in the CEA. We recorded for approximately 10-min from the bottom 384 recording sites of each shank. We found that recording sites properly targeted to the CEA could be identified by a dense band of single-unit and multi-unit activity (examples: Extended Data Fig. 7b). This process allowed us to design custom Imec readout (IMRO) tables (recording site maps; https://billkarsh.github.io/SpikeGLX/help/imroTables) for each mouse that maximized the yield of CEA neurons during subsequent experiments.

### Neuropixels acquisition and processing

We recorded 384 Neuropixels channels per session at 30 kHz with 250/500 LFP/AP gain settings using National Instruments PXI hardware and SpikeGLX software (https://billkarsh.github.io/SpikeGLX). Experimental TTL signals (representing reward cues, port entries, reward deliveries, and laser pulses) were recorded simultaneously using the same system.

Preprocessing. We used CatGT (https://billkarsh.github.io/SpikeGLX/help/catgt_tshift/catgt_tshift) to apply global common average referencing (*-gblcar*) and to isolate the action potential frequency band (*-apfilter=butter,12,300,9000*). We then used the International Brain Laboratory’s (IBL) Python Kilosort 2.5 implementation^140,141^ (https://github.com/int-brain-lab/pykilosort) to correct for sample drift along the length of the probe, detect and remove failing channels, and apply a spatial de-striping filter.

Spike sorting and curation. We also used the IBL’s Python Kilosort 2.5 implementation^140,142,143^ (https://github.com/int-brain-lab/pykilosort) for spike sorting. We then used the Phy Python package (https://github.com/cortex-lab/phy) for interactive visualization and manual curation of spike sorting output. We used Phy to classify clusters from Kilosort as single-unit (“good”) or multi-unit (“MUA”) clusters and to remove noise. We relied on waveform shape, autocorrelogram shape, spike amplitude timecourse, and cluster separation for classification. Following curation with Phy, we used MATLAB to compute three statistics for each cluster. First, we calculated each cluster’s median amplitude using the template scaling amplitudes from Kilosort (stored in *amplitudes.npy*), converted from bits to µV using the gain factor 2.34375. These template scaling amplitudes are calculated after whitening the data and are significantly smaller than the equivalent raw spike amplitudes (µV). Second, we calculated each cluster’s estimated false positive rate based on 2-ms refractory period violations^144^. Third, we calculated each cluster’s firing rate (sp/s). For experiments with multiple epochs (for example, consumption and CGRP neuron stimulation in Fig. 4), we calculated these metrics separately for each epoch and then kept the minimum median template scaling amplitude, maximum estimated false positive rate, and minimum firing rate across epochs for each cluster. We removed clusters with median template scaling amplitude < 20 µV, estimated false positive rate > 100%, or firing rate < 0.05 sp/s as noise. We classified the remaining clusters that were labeled “good” in Phy and had an estimated false positive rate < 10% as single-units, and the rest as multi-units. We included both single-unit and multi-unit clusters throughout the paper. We confirmed that our findings were consistent across a range of amplitude thresholds and for only single-unit clusters. Finally, we binned the spikes for each included neuron into 10-ms bins for downstream analysis.

Atlas alignment. All probes were coated in CellTracker CM-DiI (Invitrogen C7000) before implantation (https://www.protocols.io/view/painting-neuropixels-probes-and-other-silicon-prob-5jyl8neprl2w). After the conclusion of experiments, animals were euthanized and the brains cleared with an abbreviated version of the iDISCO+ protocol described above without immunostaining. The cleared brains were then imaged on a light sheet microscope (Lavision Ultramicroscope II) using 488-nm (autofluorescence channel) and 561-nm (CM-DiI channel) excitation light with 10 µm between horizontal planes and 5.91-µm/px resolution. Atlas alignment then followed the IBL’s pipeline^145^ (https://github.com/int-brain-lab/iblapps/wiki). The autofluorescence volume was registered to the atlas using the Brainreg Python package^146^ (https://github.com/brainglobe/brainreg) and these transformations were then directly applied to the CM-DiI volume. Individual Neuropixels shank trajectories were then manually annotated in the atlas-registered CM-DiI volume using the Brainreg-segment Napari module (https://github.com/brainglobe/brainreg-segment). Every recording site was then localized to an Allen CCF coordinate (*x,y,z*) and brain region using the IBL’s alignment GUI. The alignment process was performed separately for each Neuropixels probe shank. For all analyses, we only included neurons from recording sites that were localized to the CEA.

Multi-day recordings. We made two changes to our processing pipeline in order to track units across two recording sessions (for example, the conditioning and retrieval sessions in Fig. 5b–e and Extended Data Fig. 10a–g, and the novel and familiar sessions in Fig. 5f and Extended Data Fig. 10h–k). First, we concatenated the two recording sessions using CatGT after applying global common average referencing and isolating the action potential frequency band. We then spike sorted using the Python implementation of Kilosort 2.5 as described above. Second, during manual curation in Phy we removed clusters with obvious discontinuities or irregularities across days as noise. We then evaluated our quality metrics in MATLAB as described above. In order to improve Kilosort’s ability to track units in multi-day recordings, pairs of sessions were separated by only 1 (conditioning/retrieval) or 2 (novel/familiar) days. We then used the Spikes MATLAB package (https://github.com/cortex-lab/spikes) to extract spike waveforms and generate autocorrelograms for each recording session (Fig. 5a).

### Chronic Neuropixels analysis

For experiments involving reward delivery, we classified all neurons as novel flavor-preferring, water-preferring, or non-selective (for example, in the heatmaps in Fig. 4d and Fig. 5b). We first *z*-scored each neuron’s 10-ms binned spiking across the entire consumption period. We then calculated the average neural activity in the 10-s following every reward delivery, which was triggered by the animal entering the port. We distinguished non-selective neurons from reward-selective neurons using a Wilcoxon rank-sum test on these average responses for novel flavor and water trials while permitting a 5% FDR across all recorded neurons with the Benjamini-Krieger-Yekutieli procedure^132^. We then classified reward-selective neurons as novel flavor-preferring if their average neural activity in the 10-s following reward delivery was greater for novel flavor trials than for water trials; the remaining neurons were classified as water-preferring. We defined “Novel flavor response” as the average neural activity in the 10-s following novel flavor delivery and “Novel flavor selectivity” as the average neural activity in the 10-s following novel flavor delivery minus the average neural activity in the 10-s following water delivery (Fig. 5d,e). When correlating CGRP response to the change in flavor response and selectivity across days (Fig. 5d,e), we subtracted the baseline activity (–10 s to –5 s before reward delivery) from each trial when calculating reward responses in order to account for potential changes in baseline firing rate across days.

We generated PETHs surrounding reward delivery (–5 s to +10 s) using the *z*-scored traces calculated above. When generating reward PETHs for the second day of multi-day recordings (Retrieval day, Familiar day), we used the mean and standard deviation calculated while *z*-scoring the consumption period trace for the first day in order to ensure that units were comparable across days. We generated PETHs surrounding CGRP neuron stimulation or CGRP^CEA^ projection stimulation trains (–1 s to +4 s) using the mean and standard deviation calculated while *z*-scoring that day’s consumption period period trace, and then subtracted the baseline (–1 s to 0 s) mean of each neuron’s PETH. For plotting reward delivery PETHs as heatmaps and traces, we convolved each neuron’s PETH with an acausal half-gaussian filter with 100-ms standard deviation.

We generated delay→CGRP stim PETHs (Fig. 4d; Fig. 5b), delay→CGRP^CEA^ stim PETHs (Extended Data Fig. 9c; Extended Data Fig. 10c), and delay→LiCl PETHs (Fig. 4n,q) using the 10-ms binned spiking from 30 min before to 45 min after the onset of CGRP neuron stimulation, CGRP^CEA^ projection stimulation or 30 min after LiCl injection. We then *z*-scored these traces using the mean and standard deviation from the final 20 min of the delay period and downsampled the final normalized PETHs to 1 sample per minute for plotting. We defined “CGRP response” as the average neural activity across the entire 45-min CGRP neuron stimulation or CGRP^CEA^ projection stimulation period in the PETHs described above (Fig. 4e; Extended Data Fig. 9c; Fig. 5d,e), and “LiCl response” as the average neural activity from 5–15 min after LiCl injection in the PETHs described above (FIg. 4n,q). We generated whole-experiment PETHs (Fig. 4e,n,q; Extended Data Fig. 9c) by concatenating the 10-ms binned spiking from the final 15-min of the consumption period, the first 30-min of the delay period, and the first 45-min of the CGRP neuron stimulation period, CGRP^CEA^ projection stimulation period, or 30-min of the LiCl-induced malaise period. We then *z*-scored these traces using the mean and standard deviation of the delay period calculated above, and downsampled the final normalized PETHs to 1 sample per minute for plotting. When comparing the CGRP response (Fig. 4e,f; Extended Data Fig. 9c,d) or LiCl response (Fig. 4n,q) of novel flavor-preferring, water-preferring, and non-selective neurons, we corrected for multiple comparisons using the Hochberg-Bonferroni procedure^120^.

For the CGRP neuron stimulation→LiCl injection experiment (Extended Data Fig. 9h–j), we first classified each neuron’s CGRP response type using the strategy from the acute recording experiment described below. We first *z*-scored each neuron’s 10-ms binned spiking across the entire 10-min CGRP neuron stimulation period and generated baseline-subtracted PETHs surrounding CGRP neuron stimulation trains (–1 s to +2 s). We then applied the GMM fit on the acute recording data (Extended Data Fig. 5c) to these PETHs in order to determine each neuron’s CGRP response type and identify CGRP-activated neurons (Extended Data Fig. 9i). We then generated LiCl injection PETHs using the 10-ms binned spiking from 5 min before to 15 min after the LiCl ip. injection. We then *z*-scored these traces using the mean and standard deviation from the 5-min acclimatization period before CGRP neuron stimulation began, and baseline-subtracted the normalized PETHs using the mean activity during the period between CGRP neuron stimulation and LiCl ip. injection (–5 min to –1 min before LiCl). We downsampled the final normalized PETHs to 1 sample per minute for plotting, and then plotted average LiCl injection PETHs separately for CGRP-activated neurons and for other neurons (Extended Data Fig. 9j). We defined “LiCl response” as the average neural activity from 5–15 min after LiCl injection in the PETHs described above (Extended Data Fig. 9j).

### Decoding analysis

To identify the reactivations of neural flavor representations (Fig. 4g–i,o,r), we trained a multinomial logistic regression decoder using the LogisticRegression class from the *scikit-learn* Python package^147^ (https://scikit-learn.org) separately for each mouse using all CEA neurons during the consumption period on the conditioning day, and then evaluated this decoder across the entire conditioning session. We included all mice with > 75 simultaneously recorded CEA neurons (6/8 mice total) for this analysis. The decoder was trained to discriminate behavioral states during the consumption period across 3 categories: novel flavor consumption (which is represented by normalized spike counts within 1 s after novel flavor delivery; the normalization procedure is described in the next paragraph), water consumption (normalized spike counts within 1 s after water delivery), and baseline (normalized spike counts within 1 s before each cue onset). We trained the decoder using Lasso regularization and tested λ from 10^-4^ to 10^4^ (nine logarithmically spaced values; Extended Data Fig. 8c). We chose λ = 1 for the final decoder because it provided a high level of regularization without decreasing log-likelihood in the held-out data during 10-fold cross-validation. Cross-validation also verified that the decoder correctly identified the animal’s behavioral state (novel flavor, water, baseline) during the consumption period (Extended Data Fig. 8e).

We normalized spike counts separately for each neuron and task period. For the consumption period (decoder training), we calculated each neuron’s average spike counts within 1 s before cue onsets and subtracted it from the binned spike counts. We then divided these baseline-subtracted spike counts by the standard deviation during the consumption period. For the delay and CGRP neuron stimulation periods (decoder evaluation), we *z*-scored each neuron’s spike counts based on its mean and standard deviation during the delay period.

We then evaluated the decoder using neural activities across the session. We first used a 1-s sliding window with 150-ms steps to bin the spikes across the start to the end of the session. After obtaining the *N* neurons × *N* time-bins normalized spike counts, we used the decoder to classify the behavioral state for each time-bin based on the corresponding normalized spike counts. To visualize the decoder’s performance (Fig. 4g), we plotted the decoder output along with the simultaneously recorded neural activity (spike trains convolved with a causal half-gaussian filter with 80-ms standard deviation) grouped by flavor preference, defined using criteria described above. For clarity, we only displayed a subset of recorded neurons (50 out of 90) in the example raster: all novel flavor-preferring and water-preferring neurons, along with fifteen randomly chosen non-selective neurons. We focused our analysis on the comparison between the decoded probabilities for the novel flavor and water categories (Fig. 4g, top; Fig. 4h). We detected peaks (local maxima with values > 0.5) of the decoder output as reactivation events, and counted the number of novel and familiar reactivations with a sliding window of 1-min width and 30-s step size (Fig. 4i,o,r).

### Principal components analysis

We used the built-in MATLAB *pca* function (Fig. 4j). We began by taking the novel flavor delivery, water delivery, and CGRP neuron stimulation PETHs described above, all convolved with a causal half-gaussian filter with 100-ms standard deviation, for all reward-selective neurons (*n* = 494 pooled across all mice). We baseline-subtracted each neuron’s reward-delivery PETHs using the mean baseline activity (–5 s to –4 s before reward delivery) averaged across both trial types, and then peak-normalized each neuron’s PETHs using the maximum absolute value across both reward types. We baseline-subtracted each neuron’s CGRP neuron stimulation PETH using the mean baseline activity (–1 s to 0 s) before laser onset, and normalized each neuron’s PETH using the maximum absolute value of the PETH. To identify PC loadings and calculate variance explained, we concatenated each neuron’s novel flavor and water reward PETHs (0 s to +5 s from reward delivery), yielding a final input matrix that was 494 neurons × 1,000 time-bins for PCA. We centered every column of this matrix before performing PCA along the neuron dimension.

To plot neural trajectories during novel flavor consumption and water consumption (Fig. 4k), we used the PC loadings defined above to calculate PC1 and PC2 values for the entire population at each time-bin of the PETH (–5 s to +10 s from reward delivery). We followed an analogous procedure to plot neural trajectories during CGRP neuron stimulation. In both cases, we centered every column (time-bin).

We repeated this entire analysis using only the neurons from individual mice (Fig. 4l), or using all neurons from a separate group of mice that received CGRP^CEA^ projection stimulation (Extended Data Fig. 9e).

When analyzing changes in PC trajectories across days (Retrieval in Extended Data Fig. 10b and Familiarization in Extended Data Fig. 10k), we followed basically the same procedure as above. For these analyses, we identified PC loadings using only the first day’s (Conditioning day or Novel day) reward delivery PETHs and then used this set of PC loadings when plotting the PC trajectories for both days. Similarly, we baseline-subtracted and peak-normalized the second day’s PETHs using values that were calculated using only the first day’s PETHs. These measures ensured that PC trajectories were comparable across multi-day recordings.

### Acute Neuropixels recordings

Surgery. *Calca::Cre* mice were first injected with 400 nl of AAV5-EF1a-DIO-hChR2(H134R)-eYFP (titer: 1.2e13 GC/ml; manufacturer: PNI Viral Core Facility) into the left PB. Four weeks later, in a second surgery, an optical fiber (⌀300-µm core/0.39-NA) was implanted at a –30° angle above the injection site (see *Viral injections* and *Optical fiber implantations* for details), a steel headbar^148^ (∼1 g) was implanted at AP +1.25 mm, and and a ground pin (Newark Electronics) was placed above the right hemisphere of the cerebellum. Finally, a 2 mm^2^ recording chamber was built with Dentin above the left hemisphere extending from AP 0 mm to AP –2.0 mm and ML –2.5 mm to ML –3.5 mm. The exposed skull was removed and the brain covered with a silicone elastomer (Kwik-Cast, World Precision Instruments).

Recordings. Mice were habituated to head fixation (3× 30-min sessions). On the recording day, mice were head-fixed in a custom-built recording rig^140^, the silicone elastomer removed and the exposed brain briefly cleaned with normal saline. Neuropixels 1.0 probes^149^ had a soldered connection to short ground to external reference, which was also connected to the mouse’s ground pin during recording. Immediately prior to the start of the recording session, the probe was covered in CellTracker CM-DiI as described above for posterior histological reconstruction. A single probe was lowered (∼10 μm/s) with a µMp micromanipulator (Sensapex) into the amygdala. To prevent drying, the exposed brain and probe shank were covered with a viscous silicone polymer (Dow-Sil, Corning). Upon reaching the targeted location, the brain tissue was allowed to settle for 15 min before starting the recording. Recordings were acquired at 30 kHz in SpikeGLX. 6× locations/recordings were targeted in each animal. During the recording mice received 1 s of 10-Hz CGRP neuron stimulation followed by a 9-s inter-trial interval for a total of 10 min/60 trains. Blue light was generated using a 447-nm laser and delivered to the animal using a ⌀200-µm core patch cable. Light power was calibrated to approximately 8 mW at the fiber tip. The laser was controlled with a Pulse Pal signal generator (Sanworks 1102) programmed to deliver 5-ms laser pulses at 10 Hz.

Analysis. Spike sorting, manual curation, and atlas alignment were performed as described above for chronic Neuropixels recordings. To precisely map our electrophysiological data to the anatomical subdivisions of the amygdala in this experiment, we used the IBL’s electrophysiology alignment GUI (https://github.com/int-brain-lab/iblapps/wiki) to manually tune the alignment of each recording to the Allen CCF using electrophysiological landmarks. We then *z*-scored each neuron’s 10-ms binned spiking across the entire 10-min CGRP neuron stimulation period. We then generated PETHs surrounding CGRP neuron stimulation trains (–1 s to +2 s), and subtracted the baseline (–1 s to 0 s) mean of each neuron’s PETH (Extended Data Fig. 5c). We then used the built-in MATLAB *fitgmdist* function (*CovarianceType*=’diagonal’, *RegularizationValue*=1e-5, *SharedCovariance*=false, *Replicates*=100) to fit a gaussian mixture model (GMM) with four response types to this dataset. Specifically, we used the time-bins during stimulation (0 s to +1 s) from all amygdala neurons to generate a 3,524 neurons × 100 time-bins input matrix for GMM fitting. This GMM revealed two CGRP neuron stimulation-activated response types (shown in green; Extended Data Fig. 5c, left), one CGRP neuron stimulation-inhibited response type (shown in purple), and one unmodulated response type (shown in gray). We then plotted average CGRP neuron stimulation PETHs for each response type separately (Extended Data Fig. 5c, right), and analyzed the distribution of CGRP neuron stimulation-activated neurons across amygdala regions (Extended Data Fig. 5d,e).

### Fiber photometry (GCaMP)

We recorded CGRP neuron GCaMP signals (Fig. 3b) using a low-autofluorescence patch cable (Doric MFP_400/430/1100-0.57_0.45m_FCM-MF2.5_LAF). Excitation light was supplied at two wavelengths: isosbestic 405-nm (intensity at patch cable tip: 5–10 µW; sinusoidal frequency modulation: 531 Hz) and activity-dependent 488 nm (intensity: 15–25 µW; sinusoidal frequency modulation: 211 Hz) using an LED driver (Thorlabs DC4104). Emission light was collected through the same patch cable using a low-light photoreceiver (Newport Femtowatt 215) and then digitized using a TDT system (RZ5D) that served both as an analog-to-digital converter and a lock-in amplifier. We then low-pass filtered (2 Hz) and downsampled (100 Hz) both the isosbestic 405-nm and activity-dependent 488-nm signals. To control for photobleaching, we applied a linear fit to the isosbestic signal to align it to the activity-dependent signal and then subtracted this fitted isosbestic signal from the activity-dependent signal^150^ to obtain the final de-bleached activity-dependent GCaMP signal.

To assess changes in CGRP neuron activity due to LiCl-induced malaise, we generated peri-event time histograms (PETHs) using the 10-min before and 30-min after LiCl injection (125 mg/kg ip.). We *z*-scored the entire PETH for each mouse using the mean and standard deviation of the full 10-min before LiCl injection and then downsampled the final normalized PETHs to 1 Hz for plotting.

### Fiber photometry (AKAR2)

We recorded CEA PKA signals (Fig. 6) using the same acquisition hardware described above for GCaMP recordings. We low-pass filtered (1 Hz) and downsampled (100 Hz) both the 405-nm and 488-nm signals. Because AKAR2 is a ratiometric indicator of PKA activity^87^, we divided the 488-nm signal by the 405-nm signal to obtain the final de-bleached activity-dependent PKA signal.

This experiment was run on four consecutive days using a version of the two-reward familiarization paradigm described above (Fig. 6b). On Day 0 (Water day), both ports contained water. On Day 1 (Novel day), one port (Port A) contained the novel sweetened grape Kool-Aid flavor and the other port (Port B) contained water. On Days 2 and 3 (Familiar days), the same ports contained the same sweetened grape Kool-Aid flavor and water as on Day 1. The flavor port was counterbalanced across mice.

We generated PETHs surrounding reward delivery (–10 s to +30 s) using the final PKA signal described above. We *z*-scored these PETHs separately for each mouse and day using the following procedure. First, we centered each individual reward event PETH by subtracting the mean baseline signal (–5 to –1 s before reward delivery). Then, we concatenated this baseline epoch from all individual reward events from both ports for that mouse/day and calculated the standard deviation of this vector. Then, we divided each individual reward event PETH for that mouse/day by the calculated standard deviation. Last, we averaged across all rewards of each type (Port A, Port B) for that mouse/day (Fig. 6d,e).

To quantify PKA activity for statistical analysis, we calculated the average response from +5 to +15 s after reward delivery for each mouse/day/port using the final averaged PETHs calculated above. We then fit a GLMM using the glmmTMB R package^119^ with a gaussian link function and the formula:

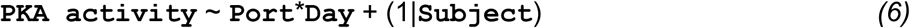

where **`PKA activity`** is as described above, **`Port`** (Port A, Port B) and **`Day`** (Day 0, Day 1, Day 2, Day 3) are fixed effect categorical variables, (1|**`Subject`**) is a random effect for each mouse, and * represents the main effects and interactions. Using the coefficients from this GLMM, we used the marginaleffects R package to calculate the marginal effect of **`Port`** (Port A – Port B) on each **`Day`**. We used the marginal effect estimates and standard errors to calculate a *P-*value for each **`Day`** with a *z*-test, and then corrected for multiple comparisons using the Hochberg-Bonferroni step-up procedure^120^ (Fig. 6f).

The recording location for each animal was determined using the same procedure as for our Neuropixels recordings described above. Briefly, we manually annotated the tip of the optical fiber lesion for each animal in the light sheet imaging data after registration to the Allen CCF and visualized these recording locations on the Allen CCF (Fig. 6g) as 100-µm circles centered on the fiber tip location for each animal.

### Exclusions

We excluded data in three instances. First, we excluded one mouse with no hM3D(Gq)-mCherry expression and one mouse with no YFP expression in the LS from Extended Data Fig. 2b. Second, we excluded one mouse with no eOPN3-mScarlet expression in the CEA from Fig. 3f. Third, we excluded confocal images with poor FISH labeling (defined as <35 total *Fos*^+^ CEA cells, or <25% of *Fos*^+^ cells also *Sst*^+^, or <25% of *Fos*^+^ cells also *Prkcd*^+^, or <25% of *Fos*^+^ cells also *Calcrl*^+^) from Fig. 3m,n (22 out of 109 images).

### Statistics and reproducibility

All data and code used to generate the figures in this paper will be made publicly available upon publication. All resources to run the deep learning-assisted Fos pipeline — including the modified Allen CCF reference atlas, modified ClearMap cell detection algorithm, trained cell classification neural network, and interactive cell density visualization tools — will be made publicly available upon publication. Design files and instructions for printing and assembling the chronic Neuropixels 2.0 implant are available at: https://github.com/agbondy/neuropixels_2.0_implant_assembly. Sample sizes were based on established standards in the field, and no statistical methods were used to predetermine sample sizes. The experiments were not randomized and the investigators were not blinded to allocation during experiments and outcome assessment. Statistical tests were performed in R and MATLAB as described. We controlled the FDR for Fos GLMMs and Neuropixels reward selectivity tests using the Benjamini-Krieger-Yekutieli two-step procedure^132^. We corrected for multiple comparisons elsewhere by controlling the family-wise error rate using the Hochberg-Bonferroni step-up procedure^120^. \**P* ≤ 0.05, \*\**P* ≤ 0.01, \*\*\**P* ≤ 0.001, \*\*\*\**P* ≤ 0.0001 throughout the paper. Individual data points are shown when practical (and always for *n* ≤ 10). Sample sizes, statistical tests, error bars, and shaded areas are defined in the figure captions.

## Acknowledgements

We thank Timothy Harris and Wade Sun for the design, manufacture, and attachment of the Neuropixels probe dovetail adapter; Thomas Luo for help troubleshooting Neuropixels recordings; Esteban Engel, Oliver Huang, Angela Chan, and the PNI Viral Core Facility for AAV production; Jeffrey Stirman and Life Canvas Technologies for light sheet imaging support; the Kleinfeld and Wang labs for collaboration on the Brainsharer web portal; Jonathan Pillow for advice on the decoding analysis; Karl Deisseroth, Douglas Kim, Richard Palmiter, Bryan Roth, Nirao Shah, Jim Wells, Ofer Yizhar, Jin Zhang, and the HHMI Janelia GENIE Project for sharing reagents; Adrian Sirko and the Princeton Laboratory Animal Resources staff for help with animal husbandry; Abigail Kalmbach and Misael Magos for technical assistance; and Peter Dayan, Zachary Knight, Brooke Jarvie, Weston Fleming, and members of the Witten lab for feedback on the manuscript. Funding was provided by the Helen Hay Whitney Foundation (to C.A.Z.), Brain Research Foundation (to I.B.W.), Simons Collaboration on the Global Brain (to I.B.W.), and National Institutes of Health (K99-DA059957 to C.A.Z., U19-NS104648 to I.B.W. and S.S.-H.W., U19-NS123716 to I.B.W., DP1-MH136573 to I.B.W., and RF1-MH128776 to S.S.-H.W.).

## Contributions

C.A.Z. conceived the project with input from I.B.W. C.A.Z. and I.B.W. designed and interpreted the experiments. I.B.W. supervised all aspects of the project. C.A.Z. performed the experiments, analyzed the data, and generated the figures with contributions from all the authors as described below. A.P.-V., with support from C.A.Z., R.N.F., and B.M., performed the acute Neuropixels recordings. B.W., with support from C.A.Z., performed the decoding analysis. E.F.K. performed stereotaxic surgeries; E.F.K. and J.B.M.-G. performed histology. S.S.B. and J.B.M.-G., with support from C.A.Z., performed the RNAscope FISH experiments. E.M.G., with support from J.L. and A.L.F., performed and analyzed the slice electrophysiology recordings. C.A.Z., with support from A.T.H., developed the deep learning-assisted cell detection pipeline. L.A.L., S.N.J., and C.A.Z., with support from S.S.-H.W., performed the tissue clearing and light sheet imaging. A.P.-V. and S.S.B. supported the chronic Neuropixels recordings. J.F.L.L. designed and manufactured behavioral equipment. A.G.B. designed the chronic Neuropixels 2.0 implant assembly. C.A.Z. and I.B.W. wrote the paper with input from all the authors.

---

**Extended Data Fig. 1.**
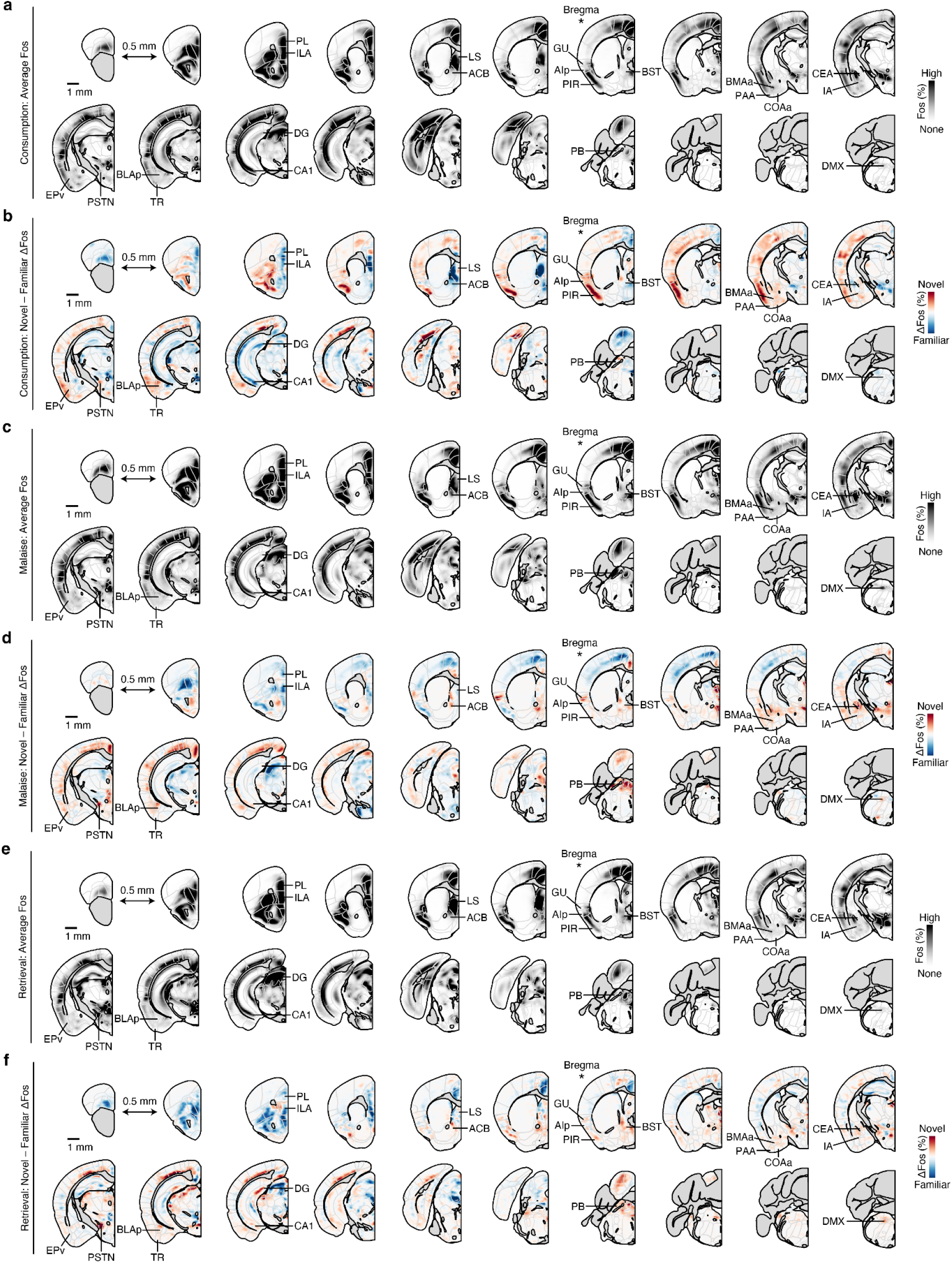
Brainwide novel versus familiar flavor activation patterns at each stage of postingestive learning. **a,** Map of average Fos^+^ cell density across all mice for the Consumption timepoint (*n* = 24 mice). The Allen CCF is overlaid. Coronal sections are spaced by 0.5 mm, the section corresponding to Bregma is marked with a *, and key brain regions are labeled. **b,** Map of the difference in average Fos^+^ cell density across Novel versus Familiar flavor condition mice for the Consumption timepoint (*n* = 12 mice per flavor condition). **c,** Map of average Fos^+^ cell density across all mice for the Malaise timepoint (*n* = 24 mice). **d,** Map of the difference in average Fos^+^ cell density across Novel versus Familiar flavor condition mice for the Malaise timepoint (*n* = 12 mice per flavor condition). **e,** Map of average Fos^+^ cell density across all mice for the Retrieval timepoint (*n* = 24 mice). **f,** Map of the difference in average Fos^+^ cell density across Novel versus Familiar flavor condition mice for the Retrieval timepoint (*n* = 12 mice per flavor condition). An interactive visualization of these Fos^+^ cell density maps is available at https://www.brainsharer.org/ng/?id=872. See Extended Data Table 1 for list of brain region abbreviations.

**Extended Data Fig. 2.**
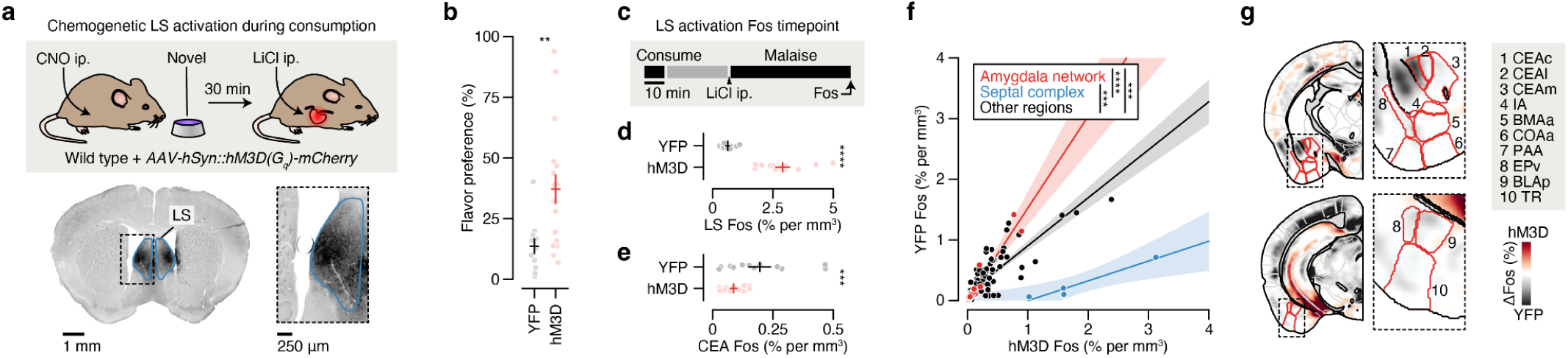
Activation of the LS during novel flavor consumption blocks malaise-driven amygdala activation and interferes with CTA acquisition. **a,** Schematic and example hM3D-mCherry expression data for the bilateral chemogenetic LS activation experiment. CNO was delivered 45-min before the experiment began to ensure that the LS was activated throughout consumption. **b**, Retrieval test flavor preference for the experiment described in **a** (*n* = 18 hM3D mice, 12 YFP mice). **c**, Schematic of the LS activation Fos timepoint (*n* = 12 mice per group). As in **a**, CNO was delivered 45-min before the experiment began, and the flavor was novel for both groups. **d**, Comparison of LS Fos for individual YFP (black) and hM3D (red) mice, confirming strong activation by hM3D. **e**, Comparison of CEA Fos for individual YFP (black) and hM3D (red) mice, showing reduced malaise-driven activation in hM3D mice. **f**, Correlation between the average Fos^+^ cell count of each brain region for hM3D versus YFP mice. The amygdala network (from Fig. 2d,e; red; *n* = 12 regions), septal complex (blue; *n* = 4 regions), and all other regions (black; *n* = 114 regions) are shown separately. **g**, Visualization of the spatially resolved difference in Fos^+^ cell density across YFP versus hM3D mice with Allen CCF boundaries overlaid. *P*-value in **b** is from a Wilcoxon rank-sum test. *P*-values in **d**,**e** are from GLMM coefficient estimate *z*-tests. *P*-values in **f** are from a one-way analysis of covariance model corrected for multiple comparisons across brain region groups. Error bars represent mean ± s.e.m. Shaded areas in **f** represent 95% confidence interval for linear fit. \*\**P* ≤ 0.01, \*\*\**P* ≤ 0.001, \*\*\*\**P* ≤ 0.0001.

**Extended Data Fig. 3.**
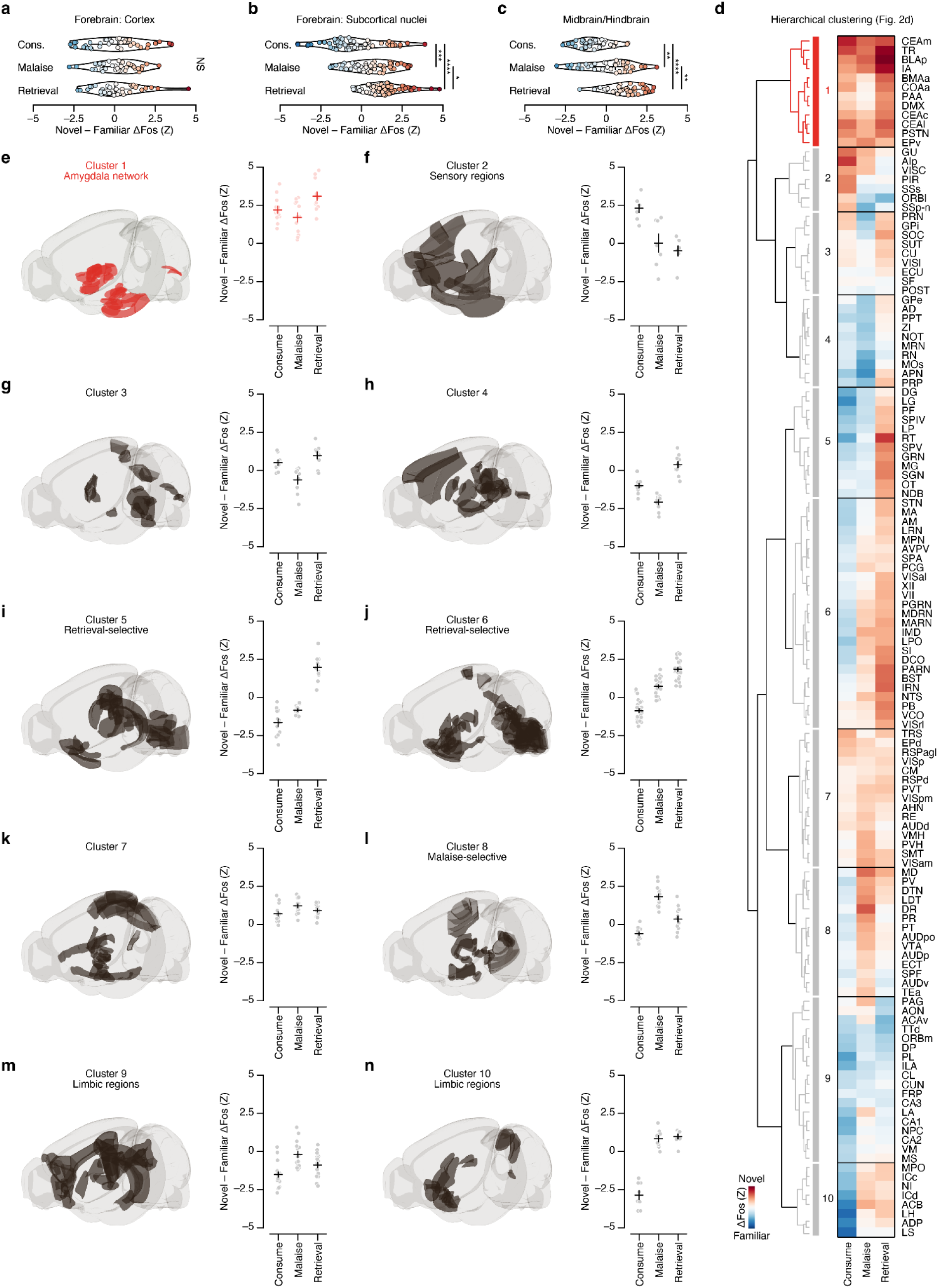
Hierarchical clustering of brain regions based on novel versus familiar flavor activation patterns. Panels **a**–**c** show that the brainwide shift towards activation by the novel flavor is primarily localized to subcortical regions. **a,** Novel – Familiar ΔFos effect distribution of all cortical regions (Cerebral Cortex in the Allen CCF) at each timepoint (*n* = 38 brain regions). **b,** Novel – Familiar ΔFos effect distribution of all subcortical forebrain (Cerebral Nuclei, Thalamus, and Hypothalamus in the Allen CCF) regions at each timepoint (*n* = 54 brain regions). **c,** Novel – Familiar ΔFos effect distribution of all midbrain and hindbrain regions (Midbrain, Pons, and Medulla in the Allen CCF) at each timepoint (*n* = 38 brain regions). **d,** Hierarchical clustering of Novel – Familiar ΔFos effects. This is an expanded version Fig. 2d showing all brain region names. **e**–**n**, Left: Illustration of the brain regions comprising each cluster from the hierarchical clustering analysis. Right: Summary of the Novel – Familiar ΔFos effect for each cluster at each timepoint, showing each brain region as an individual point. *P*-values in **b,c** are from Kolmogorov-Smirnov tests corrected for multiple comparisons across timepoints. No statistical tests were performed for **e**–**f**. Error bars represent mean ± s.e.m. Outlines in **a**–**c** represent kernel-density estimates of the empirical distributions. \**P* ≤ 0.05, \*\**P* ≤ 0.01, \*\*\**P* ≤ 0.001, \*\*\*\**P* ≤ 0.0001, NS, not significant (*P* > 0.05). See Extended Data Table 1 for list of brain region abbreviations.

**Extended Data Fig. 4.**
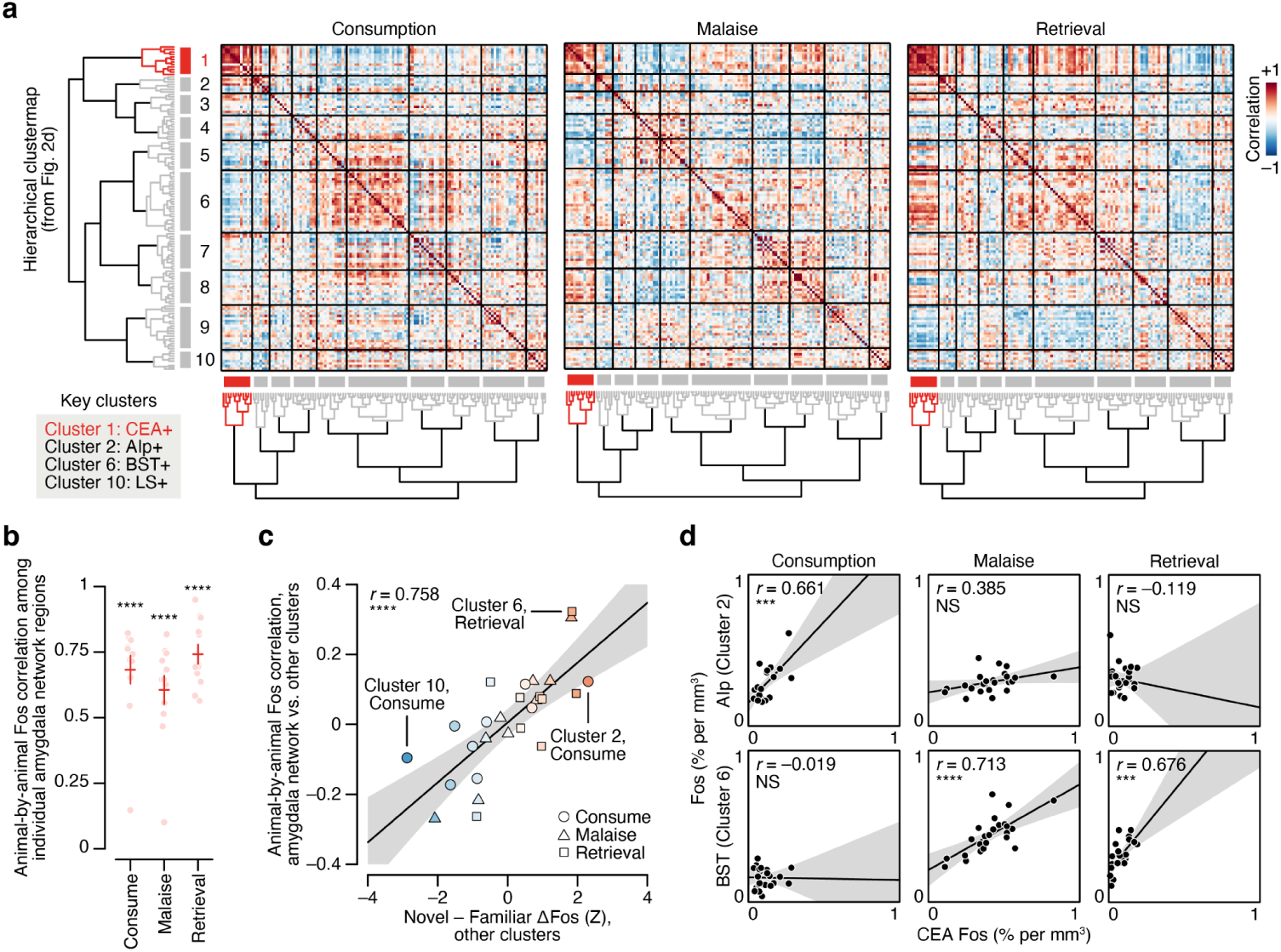
The amygdala cluster forms a functional network. **a,** Correlation matrices showing the animal-by-animal pairwise Fos correlation for every pair of brain regions during consumption (left), delayed malaise (middle), or memory retrieval (right). Brain regions are sorted using the hierarchical clustermap obtained from the Novel – Familiar ΔFos effects in Fig. 2d. **b,** Summary of the average within-cluster Fos correlation for individual amygdala network regions (Cluster 1 from Fig. 2d,e) by timepoint (*n* = 12 regions). The high animal-by-animal correlation among all of the regions in cluster 1 suggest that these regions form a functional network. Panels **c,d** show that activation of other clusters of brain regions is more correlated with amygdala network activation at experimental timepoints when those clusters are more strongly novel flavor-selective, including for clusters that were specifically engaged during the initial flavor consumption (comprising sensory cortices; cluster 2) or during retrieval (including the BST; cluster 6). This suggests that the amygdala network may play a role in orchestrating the brainwide response to novel flavors at different stages of learning. **c,** Summary of the average across-cluster Fos correlation between the amygdala network and every other cluster at each timepoint as a function of the other cluster’s standardized Novel – Familiar effect at that timepoint (*n* = 9 clusters × 3 timepoints). **d,** Scatter plots showing the pairwise correlation between AIp (top; example Cluster 2 region) or BST (bottom; example Cluster 6 region) and the CEA (*n* = 24 mice per timepoint). *P*-values in **b** are from Wilcoxon signed-rank tests corrected for multiple comparisons across timepoints. *P*-values in **c,d** are from Pearson correlation *t*-tests. Error bars represent mean ± s.e.m. Shaded areas in **c,d** represent 95% confidence interval for linear fit. \*\*\**P* ≤ 0.001, \*\*\*\**P* ≤ 0.0001, NS, not significant (*P* > 0.05). See Extended Data Table 1 for list of brain region abbreviations.

**Extended Data Fig. 5.**
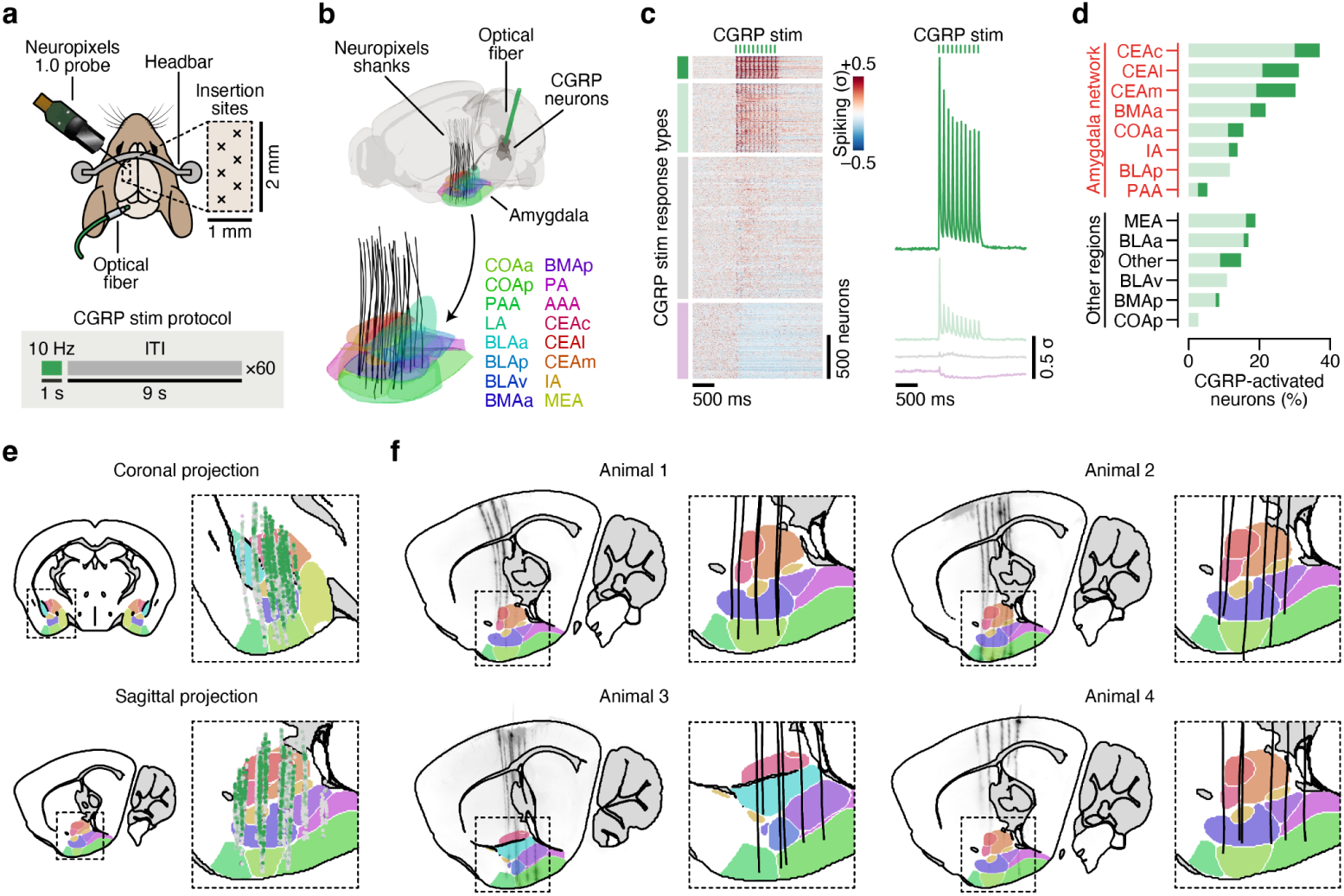
An electrophysiological atlas of the effects of CGRP neuron stimulation on amygdala activity in vivo. **a,** Schematic of the acute Neuropixels recording experiment. **b,** Reconstruction of recording trajectories registered to the Allen CCF. Each line represents one insertion of a single-shank Neuropixels 1.0 probe targeting the amygdala (*n* = 24 insertions from 4 mice). **c,** Left: PETHs of neural activity time-locked to CGRP neuron stimulation trains (*n* = 3,524 amygdala neurons from 24 insertions). Neurons were divided into four response types using a GMM model (see Methods): two CGRP neuron stimulation-activated response types (7.3% strongly activated, dark green; 22.4% weakly activated, light green), one CGRP neuron stimulation-inhibited response type (24.6%, purple), and one unmodulated response type (45.8%, gray). Right: Average PETHs for each GMM response type. **d,** Percentage of recorded neurons that were CGRP neuron stimulation-activated based on the GMM across amygdala subregions (*n* = 339 CEAc, 272 CEAl, 717 CEAm, 526 BMAa, 129 COAa, 133 IA, 41 BLAp, 30 PAA, 182 MEA, 354 BLAa, 54 Other (AAA, LA, PA), 44 BLAv, 250 BMAp, and 58 COAp neurons). Regions in the CTA amygdala network (Cluster 1 from Fig. 2d,e) are shown in red, and other amygdala regions are shown in black. **e,** Anatomical distribution of all CGRP-activated (green), CGRP-inhibited (purple), and unmodulated (gray) neurons projected onto a single coronal or sagittal section of the Allen CCF. **f,** Left: Light sheet imaging data for each animal of Neuropixels probe trajectories aligned to the Allen CCF with amygdala subregions overlaid. Right: Reconstruction of recording trajectories in the amygdala for each animal. For each animal, a single sagittal section corresponding to the center-of-mass of all active recording sites is shown. The colormap for amygdala regions in **b** is also used in **e** and **f**. See Extended Data Table 1 for list of brain region abbreviations.

**Extended Data Fig. 6.**
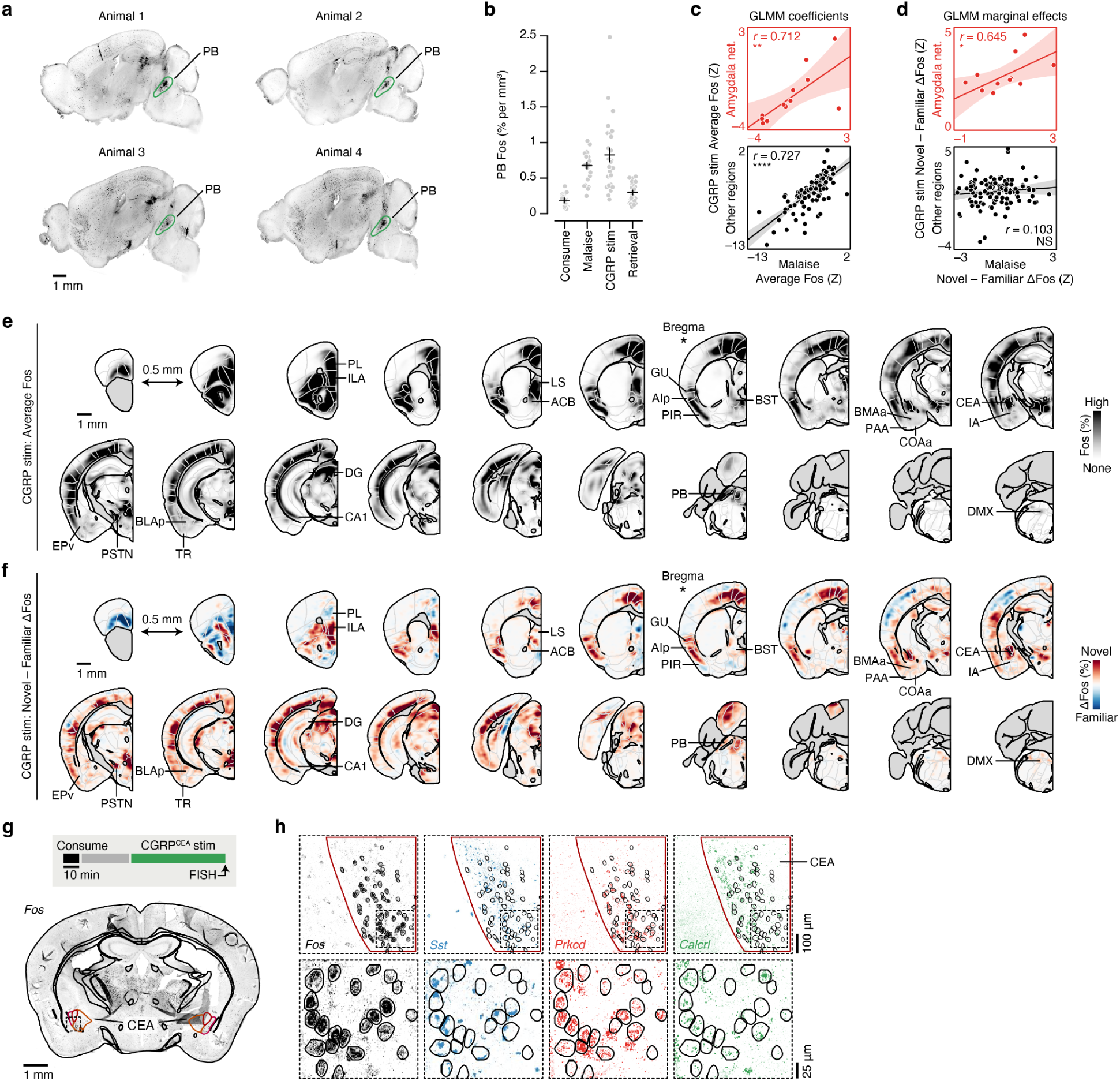
Brainwide novel versus familiar flavor activation pattern during CGRP neuron stimulation. **a,** Example brainwide Fos imaging data (200-µm maximum intensity projections) for four example CGRP stim timepoint animals. Strong Fos expression in the PB driven by optogenetic stimulation of CGRP neurons is highlighted in green. **b,** Summary of Fos^+^ cell counts in the PB at each experimental timpoint, confirming high levels of neural activation during both LiCl-induced malaise and CGRP neuron stimulation (*n* = 24 mice for Consumption, 24 mice for Malaise, 27 mice for CGRP stim, and 24 mice for Retrieval). **c,** Analysis analogous to Fig. 3i but using the Fos GLMM from Equation 3. Here, the correlation among the standardized coefficients (*Z* = estimate/standard error) for the main LiCl effect and main CGRP stim effect on Fos^+^ cell counts from the regression **`Fos counts`** ∼ **`Timepoint`** + **`Sex`** + (1|**`Batch`**) + ln(**`PB Counts`**) are plotted (*n* = 12 amygdala network regions, 117 other regions). **d,** Analysis analogous to Fig. 3j but using the Fos GLMM from Equation 4. Here, the correlation among the average marginal effect (Novel – Familiar ΔFos, in standardized units) of flavor on Fos^+^ cell counts from the regression **`Fos counts`** ∼ **`Novel*Timepoint`** + **`Sex`** + (1|**`Batch`**) + ln(**`PB Counts`**) are plotted (*n* = 12 amygdala network regions, 117 other regions). **e,** Map of average Fos^+^ cell density across all mice for the CGRP stim timepoint (*n* = 27 mice). The Allen CCF is overlaid. Coronal sections are spaced by 0.5 mm, the section corresponding to Bregma is marked with a *, and key brain regions are labeled. **f,** Map of the difference in average Fos^+^ cell density across Novel versus Familiar flavor condition mice for the CGRP stim timepoint (*n* = 14 novel flavor mice, 13 familiar flavor mice). **g**, Top: Schematic of the CGRP^CEA^ projection stimulation RNAscope FISH experiment. Bottom: Example slide scanner image of *Fos* expression from one coronal section with the Allen CCF overlaid. **h**, Example confocal image showing *Fos* expression (as a marker of neural activation) and *Sst*, *Prkcd*, and *Calcrl* expression (as markers of known CEA cell types) in the CEA following CGRP^CEA^ projection stimulation. This is an expanded version of Fig. 3l. The top row shows the full field-of-view for each gene, and the bottom row is magnified with *Fos*^+^ cell outlines overlaid in black. *P*-values in **c,d** are from Pearson correlation *t*-tests. No statistical tests were performed for **b**. Error bars represent mean ± s.e.m. Shaded areas in **c,d** represent 95% confidence interval for linear fit. \**P* ≤ 0.05, \*\**P* ≤ 0.01, \*\*\*\**P* ≤ 0.0001, NS, not significant (*P* > 0.05). See Extended Data Table 1 for list of brain region abbreviations and for GLMM statistics.

**Extended Data Fig. 7.**
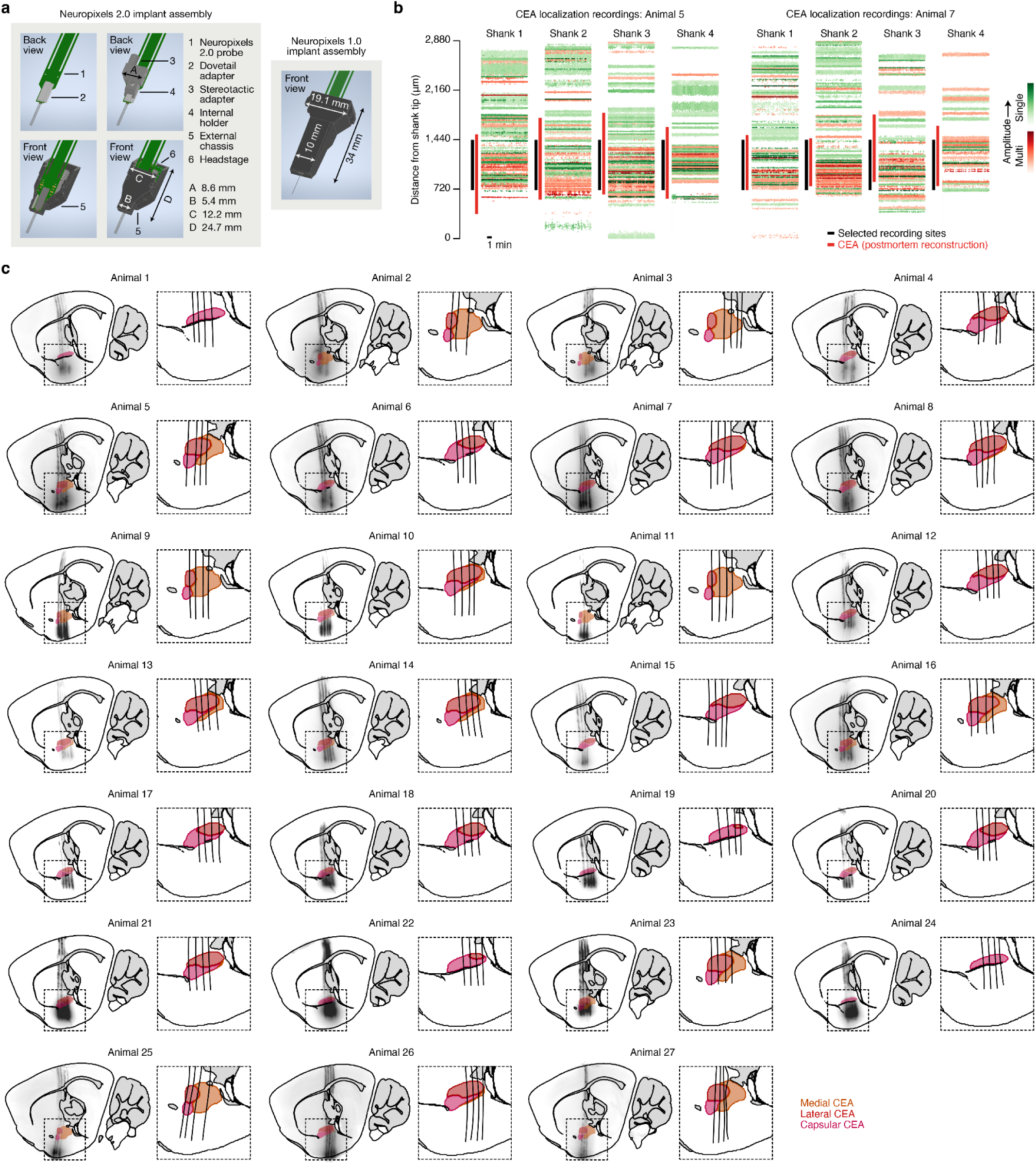
Chronic Neuropixels electrophysiology in the amygdala of freely moving mice. **a,** Left: A schematic illustration of the chronic Neuropixels 2.0 implant assembly at progressive stages of construction from top left to bottom right. Right: A schematic illustration of the chronic Neuropixels 1.0 implant assembly^69^ upon which the 2.0 implant design is based, shown for size comparison. **b,** Test recordings used to select recording sites (black bars) properly targeting the CEA (red bars, based on postmortem reconstruction) for two example animals. The shanks are arranged from anterior (1, left) to posterior (4, right) and span 750-µm total. We found that we could distinguish the CEA from nearby brain regions along the vertical axis of the probe shanks as a band of dense neural activity. **c,** Left: Light sheet imaging data for each animal of Neuropixels shank trajectories aligned to the Allen CCF with CEA subregions overlaid. Right: Reconstruction of trajectories in the CEA. For each animal, a single sagittal section corresponding to the center-of-mass of all active recording sites is shown. See Methods for a description of which animals were included in each experiment.

**Extended Data Fig. 8.**
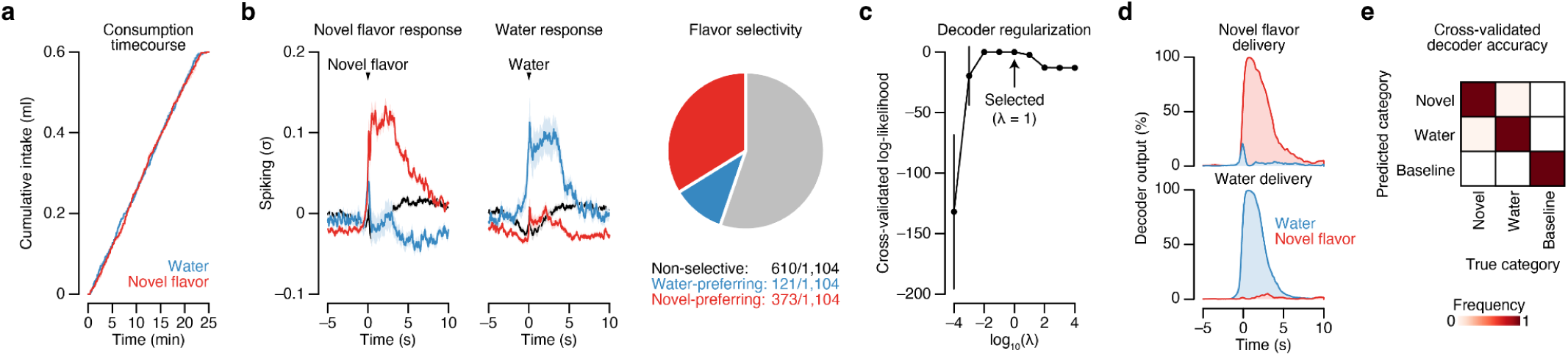
The amygdala is strongly activated by a novel flavor and near-perfectly discriminates flavor. **a,** Cumulative intake of the novel flavor (red) and water (blue) in the two-reward CTA paradigm in Fig. 4b (*n* = 8 mice). We used a randomized, trial-based structure to ensure that mice consumed the two options at an equal rate (see Methods). **b,** Left: PETHs to novel flavor delivery for the novel flavor-preferring (*n* = 373 neurons from 8 mice), water-preferring (*n* = 121 neurons), and non-selective (*n* = 610 neurons) CEA neurons in Fig. 4c–l. Middle: PETHs to water delivery for the novel flavor-preferring, water-preferring, and non-selective CEA neurons. Right: Pie chart visualizing the proportion of novel flavor-preferring, water-preferring, and non-selective CEA neurons. Panels **c–e** provide additional characterization of the multinomial logistic regression decoder using CEA population activity. **c,** Cross-validated log-likelihood for the decoder classifying periods of novel flavor consumption versus water consumption versus baseline activity across a range of regularization parameter (λ) values (*n* = 6 mice). **d,** Decoder output time-locked to novel flavor delivery (top) and water delivery (bottom) in the initial consumption period (mean across 6 mice). The decoder’s predicted probability for novel flavor consumption (red) and water consumption (blue) are both shown. **e,** Confusion matrix summarizing the cross-validated decoder performance across all mice (*n* = 6 mice). The overall misclassification rate was 0.56% (4 out of 720). Error bars and shaded areas represent mean ± s.e.m.

**Extended Data Fig. 9.**
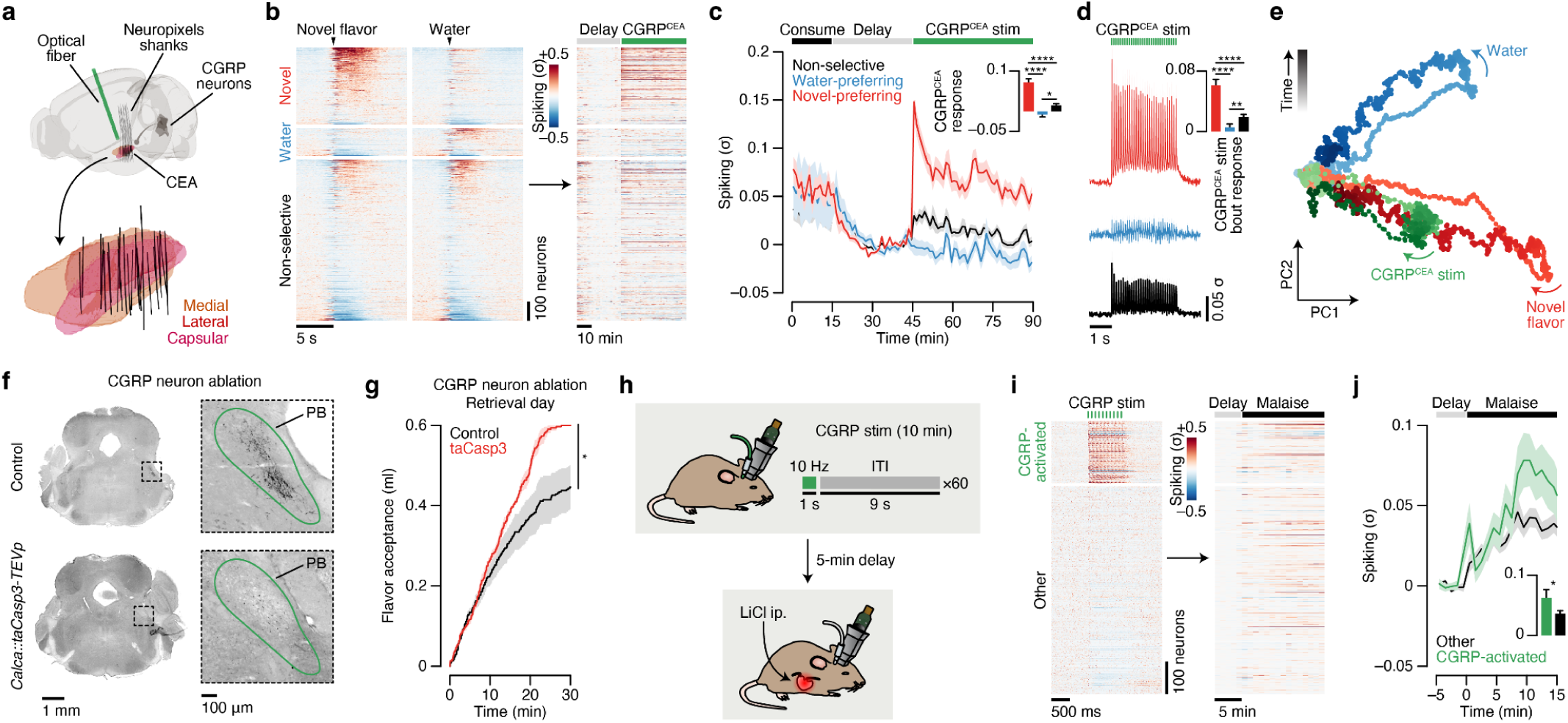
CGRP^CEA^ projection stimulation reactivates novel flavor representations in the amygdala, and CGRP neuron ablation impairs delayed CTA learning. Panels **a**–**e** show that postingestive CGRP^CEA^ projection stimulation reactivates novel flavor representations in the amygdala. **a**, Reconstruction of recording trajectories registered to the Allen CCF for mice with CGRP^CEA^ projection stimulation (*n* = 32 shanks from 8 mice). **b**, Heatmap showing the trial-average spiking of all recorded CEA neurons (*n* = 1,221 neurons from 8 mice) to novel flavor and water consumption during the consumption period (left) and during the delay and CGRP^CEA^ projection stimulation periods (right). Neurons are grouped by their novel flavor/water preference and then sorted by consumption response magnitude. Consumption PETHs are time-locked to delivery of the flavor or water. **c**, Average spiking of the novel flavor-preferring (red; *n =* 354 neurons), water-preferring (blue; *n* = 129 neurons), and non-selective (black; *n* = 738 neurons) populations across the entire experiment. The inset quantifies the average response of each population during the entire 45-min CGRP^CEA^ projection stimulation period. **d**, Average spiking of the novel flavor-preferring (red), water-preferring (blue), and non-selective (black) populations during individual 3-s bouts of 10-Hz CGRP^CEA^ projection stimulation. The inset quantifies the average response of each population within bouts of CGRP^CEA^ projection stimulation. **e,** Neural trajectories for novel flavor consumption (red), water consumption (blue), and CGRP^CEA^ projection stimulation in PC-space from a population activity dimensionality reduction analysis analogous to Fig. 4k. Panels **f**,**g** show that genetic ablation of CGRP neurons by taCasp3-TEVp impairs delayed CTA learning. **f**, Example CGRP immunoreactivity data confirming genetic ablation of CGRP neurons by taCasp3-TEVp. This is an expanded version of Fig. 4p. **g**, CGRP neuron ablation mice (taCasp3) show significantly higher acceptance of the conditioned flavor following LiCl-induced CTA when compared to wild type controls (*n* = 6 taCasp3 mice, 7 control mice). This data was collected during the retrieval session in the two-reward CTA paradigm, where 0.6-ml is the maximum possible flavor acceptance. Panels **h**–**j** show that CGRP-activated CEA neurons are also activated by LiCl injection. **h,** Schematic of the paradigm for tracking neurons across CGRP neuron stimulation and LiCl-induced malaise. **i,** Heatmap showing the trial-average spiking of all recorded CEA neurons (*n* = 833 neurons from 4 mice) to CGRP neuron stimulation (left) and then during LiCl-induced malaise (right). Neurons were classified as CGRP neuron stimulation-activated using the GMM trained on the data in Extended Data Fig. 5c. **j,** Average spiking of the CGRP neuron stimulation-activated neurons (green; *n* = 189 neurons) and other neurons (black; *n* = 634 neurons) during LiCl-induced malaise. The inset quantifies the average response of each population. *P*-values in **c,d,g,j** are from Wilcoxon rank-sum tests corrected for multiple comparisons when appropriate (across neuron groups in **c,d**). Error bars and shaded areas represent mean ± s.e.m. \**P* ≤ 0.05, \*\**P* ≤ 0.01, \*\*\*\**P* ≤ 0.0001, NS, not significant (*P* > 0.05).

**Extended Data Fig. 10.**
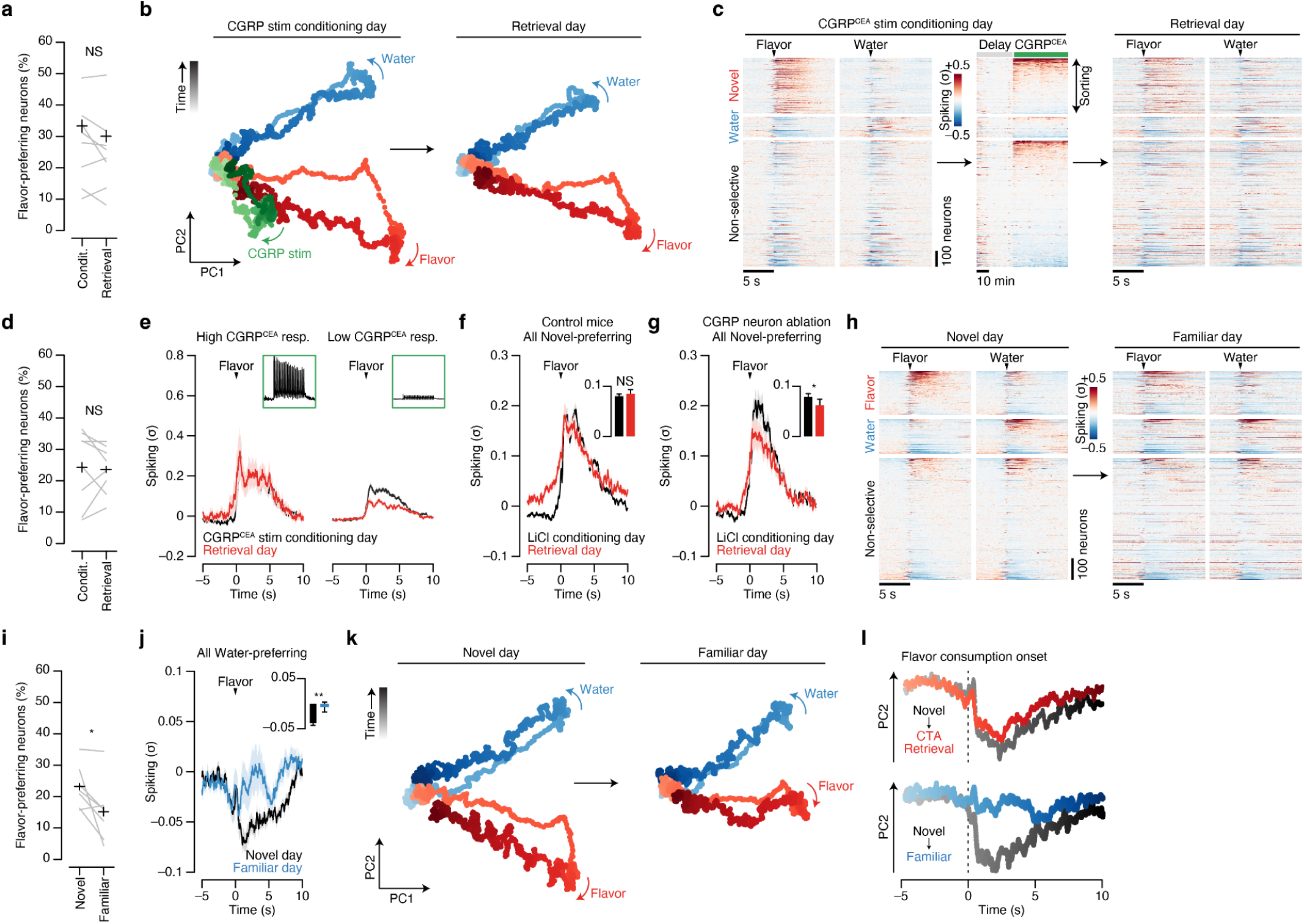
Postingestive CGRP neuron activity is necessary and sufficient to stabilize novel flavor representations in the amygdala upon memory retrieval. Panels **a**,**b** present further analysis of the CGRP neuron stimulation conditioning experiment in Fig. 5b–d. **a,** Summary of the proportion of flavor-preferring units classified separately on conditioning day or on retrieval day (*n* = 8 mice). **b,** Neural trajectories for flavor consumption (red), water consumption (blue), and CGRP neuron stimulation in the PC-space on conditioning day (left) and retrieval day (right). Trajectories for both days were calculated using the PCA loadings from conditioning day, and the novel flavor trajectory remained largely stable. Panels **c**–**e** present further analysis of the CGRP^CEA^ projection stimulation conditioning experiment in Fig. 5e. **c**, Heatmap showing the average spiking of all recorded neurons (*n* = 1,042 neurons from 8 mice) to novel flavor and water consumption during the consumption period (left) and during the delay and CGRP^CEA^ projection stimulation periods (middle) on conditioning day, and the responses of the same neurons to flavor and water consumption on retrieval day. Neurons are grouped by their novel flavor/water preference on pairing day and then sorted by CGRP^CEA^ response magnitude. **d**, Analogous to **a**, but for mice with CGRP^CEA^ projection stimulation (*n* = 8 mice). **e**, Trial-average spiking of the novel-preferring neurons with the highest 10% of CGRP^CEA^ response magnitudes (High CGRP^CEA^ resp.; left) and of the remaining novel-preferring neurons (Low CGRP^CEA^ resp.; right) on conditioning day and retrieval day. The insets show the average CGRP^CEA^ projection stimulation response profile of each subpopulation. Panels **f**,**g** show that LiCl-induced malaise stabilizes CEA novel flavor representations during memory retrieval, and that this stabilization is impaired in mice with CGRP neuron ablation. **f**, For control mice, trial-average spiking of the novel flavor-preferring population (*n* = 279 neurons) of CEA neurons during flavor consumption on conditioning day (black) and retrieval day (red). The inset quantifies the average response on each day. **g**, Analogous to **f**, but for mice with CGRP neuron ablation (*n* = 109 neurons from 4 mice). Panels **h**–**k** present further analysis of the familiarization experiment in Fig. 5f. **h,** Heatmap showing the average spiking of all recorded neurons (*n* = 924 neurons from 7 mice) to flavor and water consumption on novel day (left) and, two days later, on familiar day (right). Neurons are grouped by their novel flavor/water preference on novel day and then sorted by reward response magnitude. **i,** Summary of the proportion of flavor-preferring units classified separately on novel day or on familiar day for the familiarization experiment (*n* = 7 mice). **j,** Trial-average spiking of the initially water-preferring population (*n* = 160 neurons from 7 mice; classified on novel day) during flavor consumption on novel day (black) and familiar day (blue). The inset quantifies the average response on each day. **k,** Neural trajectories for flavor consumption (red) and water consumption (blue) in the PC-space on novel day (left) and familiar day (right). Trajectories for both days were calculated using the PCA loadings from the novel day, and the flavor trajectory was strongly degraded following familiarization. **l,** Comparison of the time-courses specifically along the PC2 axis during consumption for the conditioning with CGRP neuron stimulation (from the data in **b**) and familiarization (from the data in **k**) experiments. Top: The flavor consumption trajectory on conditioning day (“Novel”) is shown in grays and the stable trajectory on retrieval day (“CTA Retrieval”) is shown in reds. Bottom: The flavor consumption trajectory on novel day (“Novel”) is shown in grays and the degraded trajectory on familiar day (“Familiar”) is shown in blues. *P*-values in **a,d,f,g,i,j** are from Wilcoxon signed-rank tests. Error bars and shaded areas represent mean ± s.e.m. \**P* ≤ 0.05, \*\**P* ≤ 0.01, NS, not significant (*P* > 0.05).

**Extended Data Table 1.**
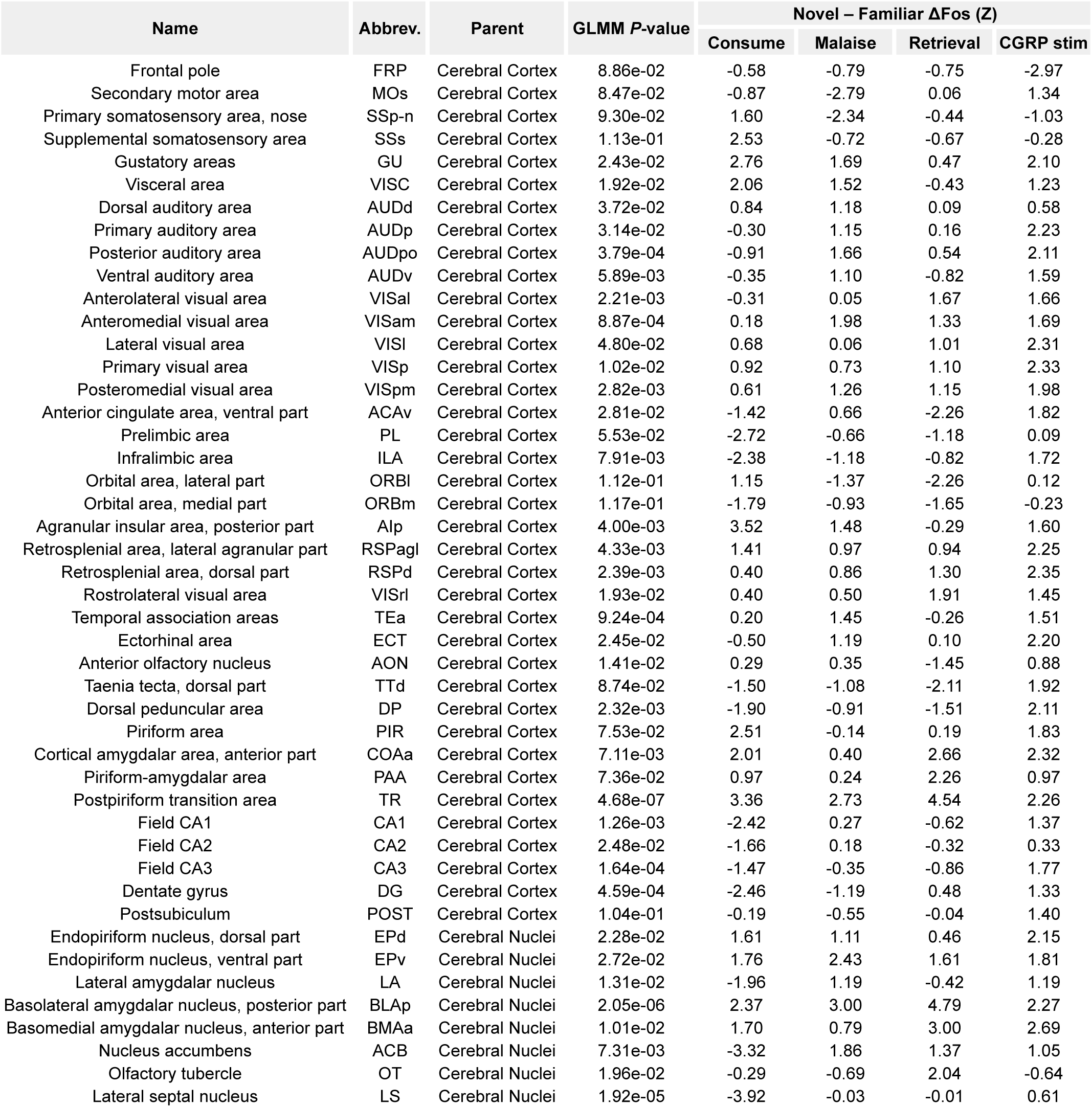
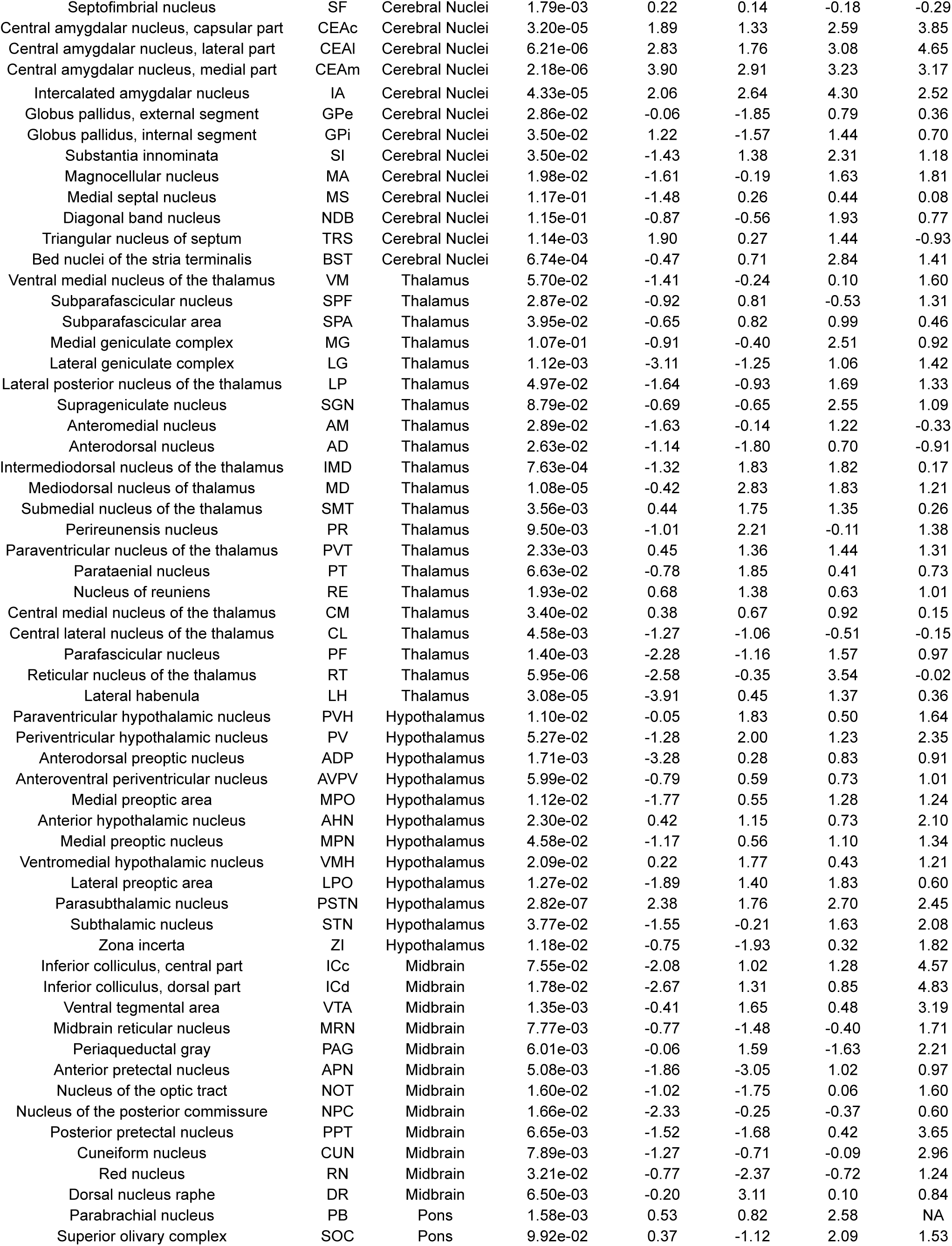
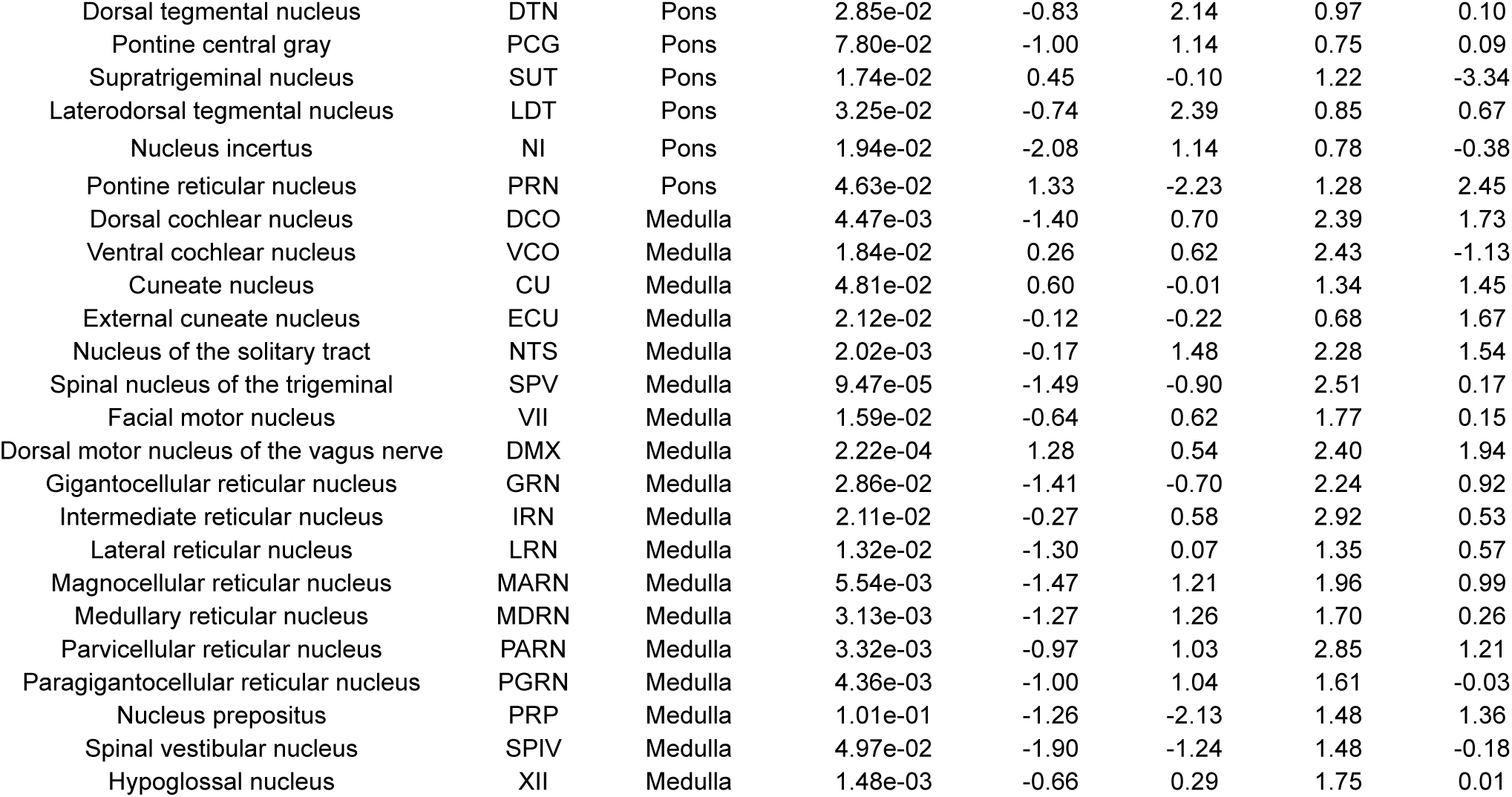
Fos GLMM statistics for individual brain regions. Summary information for each brain region in the Allen CCF that met the following criteria: (1) total volume ≥0.1 mm^3^; (2) lowest level of its branch of the ontology tree (cortical layers not included); and (3) **`Flavor*Timepoint`** predictors (all main effects and interactions) significantly improved overall Fos GLMM performance (Equation 2; see Methods). *P*-values are from likelihood-ratio χ^2^-tests comparing full and reduced models. Uncorrected *P*-values are reported here. The critical *P*-value for significance while permitting a 10% FDR using the Benjamini-Krieger-Yekutieli two-step procedure^132^ was 0.1172. Of 200 regions that met criteria (1) and (2), 130 also met criterion (3) and are listed below. Novel – Familiar ΔFos values for the Consumption, Malaise, and Retrieval timepoints were calculated using the GLMM from Equation 2, and values for the CGRP stim timepoint were calculated using the GLMM from Equation 4.

